# Dirichlet-multinomial modelling outperforms alternatives for analysis of microbiome and other ecological count data

**DOI:** 10.1101/711317

**Authors:** Joshua G. Harrison, W. John Calder, Vivaswat Shastry, C. Alex Buerkle

## Abstract

Molecular ecology regularly requires the analysis of count data that reflect the relative abundance of features of a composition (e.g., taxa in a community, gene transcripts in a tissue). The sampling process that generates these data can be modeled using the multinomial distribution. Replicate multinomial samples inform the relative abundances of features in an underlying Dirichlet distribution. These distributions together form a hierarchical model for relative abundances among replicates and sampling groups. This type of Dirichlet-multinomial modelling (DMM) has been described previously, but its benefits and limitations are largely untested. With simulated data, we quantified the ability of DMM to detect differences in proportions between treatment and control groups, and compared the efficacy of three computational methods to implement DMM—Hamiltonian Monte Carlo (HMC), variational inference (VI), and Gibbs Markov chain Monte Carlo. We report that DMM was better able to detect shifts in relative abundances than analogous analytical tools, while identifying an acceptably low number of false positives. Among methods for implementing DMM, HMC provided the most accurate estimates of relative abundances, and VI was the most computationally efficient. The sensitivity of DMM was exemplified through analysis of previously published data describing lung microbiomes. We report that DMM identified several potentially pathogenic, bacterial taxa as more abundant in the lungs of children who aspirated foreign material during swallowing; these differences went undetected with different statistical approaches. Our results suggest that DMM has strong potential as a statistical method to guide inference in molecular ecology.

## Introduction

In many scientific disciplines, data from both manipulative experiments and surveys of natural variation are often counts of observations that are assigned to categories. Given some total level of observational effort, the counts of the different features in the sample (e.g., taxa or transcripts) reflect the underlying proportions of those features in the sampled composition (e.g., an assemblage of organisms or collection of molecules). In molecular ecology, such sampling can take the form of detecting and counting taxa based on observed DNA sequences (e.g., in molecular barcoding or microbial ecology) or counting the reads assigned to specific transcripts in studies of gene expression (Fernandes et al. 2014, Gloor et al. 2017, Tsilimigras and Fodor 2016). For these applications, sampling effort corresponds to the total number of sequence reads, and the count of reads assigned to a taxon or gene supports inference of their true proportion in the composition. Moreover, the total number of reads that can be obtained is constrained by the sequencing instrument, with reads ascribed to samples and features within each sample. Due to this constant sum constraint, compositional data have the important quality that as the relative abundance of one feature in the composition increases, other features must decrease.

Molecular ecologists often rely on compositional count data to define differences between sampling groups. As an example, we may wish to know how the foliar and root microbiomes of a particular plant taxon differ. To answer this question, an understanding of how each feature shifts in relative abundance among sampling groups is required. In our view, if even a single feature shifts in relative abundance among groups, then this demonstrates an effect of sampling group that could be biologically interesting, albeit subtle. Such effects will go unnoticed if analyses rely on techniques such as ordination and PERMANOVA, which can provide insight into overall differences between sampling groups (McKnight et al. 2019), but provide no statistical model to identify those features that may differ in relative abundance among groups. Accordingly, a variety of methods have been developed to perform the seemingly simple task of determining treatment-induced shifts in relative abundance, which is often referred to as “differential relative abundance testing” or “differential expression” testing (the latter phrase arises because the roots of many of these methods lie within the field of functional genomics; Bullard et al. 2010, Dillies et al. 2013, Paulson et al. 2013, Thorsen et al. 2016, Weiss et al. 2017).

Methods for detecting shifts in relative abundance vary tremendously—and the benefits and drawbacks of various methods are the subjects of an ongoing dialogue (e.g., Bullard et al. 2010, McMurdie and Holmes 2014, Weiss et al. 2017). Early approaches typically relied on repeated frequentist tests after transforming count data to account for differences in sampling effort among replicates or sampling groups, typically via rarefaction, conversion to proportions, or, for transcriptomic data, reads per kilobase per million mapped reads (Bullard et al. 2010). More recently, rarefaction has been criticized because it can amplify the variation present within replicates and thus reduce statistical power (McMurdie and Holmes 2014; but see McKnight et al. 2019 and Weiss et al. 2017 for counterarguments). Numerous statistical modelling approaches have arisen to account for the challenges imposed by compositional data, while avoiding rarefaction. These methods often model feature relative abundance and typically involve some form of normalization followed by repeated frequentist testing. Methods most often differ in the choice of distribution(s) utilized for modelling and normalization method employed. For example, the software DESeq2 (Love et al. 2014) and edgeR (Robinson et al. 2010) are widely-used for analysis of gene expression data and, more recently, for microbiome analysis (Weiss et al. 2017). These tools model feature relative abundances using a negative binomial distribution (a reparameterization of the Poisson distribution to allow for overdispersion), which is scaled to account for variation in sequencing depth among samples (each tool uses different normalization methods). Next a generalized linear model is used to determine if features differ in relative abundance between sampling groups. By comparison, the popular ANCOM software applies a centered log ratio transformation (Aitchison 1982) to the data followed by repeated parametric or non-parametric testing (depending on the data) with multiple comparison correction. These few examples serve to illustrate the variety of approaches available for performing differential expression testing. However, we are unaware of any popular method that allows estimates of feature relative abundance to be easily extracted while preserving the uncertainty in those estimates for propagation to downstream analyses. This perceived need led us to consider modelling feature relative abundances using the Dirichlet and multinomial distributions (Box 1) in a Bayesian framework.

The multinomial and Dirichlet probability distributions are the relevant models of the aforementioned sampling process that commonly leads to compositional data. Statistical modelling using these distributions has proven successful in a number of biological studies. For instance, Fordyce et al. (2011) rely on Dirichlet-multinomial modelling (DMM) to analyze ecological count data, such as counts of behavioural and dietary choices of animals (also see Coblentz et al. 2017). Similar models have been applied to large counts of DNA sequences—for instance, Fernandes et al. (ALDEx2, 2014), Nowicka and Robinson (DRIM-Seq, 2016), and Rosa et al. (HMP, 2012) use DMM to estimate and compare feature-specific relative abundances in transcriptomes and microbiomes. Additionally, DMM has been used to model mixtures of compositions, a situation that could arise in a laboratory-derived microbial assemblage occurring as a contaminant within samples, or in mixtures of different communities in nature (MicrobeDMM, Holmes et al. 2012; SourceTracker, Knights et al. 2011; BioMiCo, Shafiei et al. 2015; FEAST, Shenhav et al. 2019; ecostructure, White et al. 2019). Likewise, DMM has been used to estimate association networks among microbial taxa (SparCC, Friedman and Alm 2012; mLDM, Yang et al. 2017).

These models represent important advances and demonstrate the utility of DMM, but it remains unclear how data attributes, such as rank-abundance profiles and dimensionality, affect the accuracy and precision of parameter estimates. Moreover, compared to models that rely on other distributions or are based on different statistical methods (likelihood and frequentist methods), Bayesian DMM can be computationally demanding. Recent advances in computational statistics such as Hamiltonian Monte Carlo (HMC) sampling and variational inference (VI, see Methods; Blei et al. 2017, Monnahan et al. 2017) may improve model runtime, but the accuracy and performance of these new methods remains to be evaluated in different modelling contexts.

Consequently, we conducted a simulation experiment to learn the limits and benefits of DMM through the analysis of data that encompass much of the variety in attributes encountered across scientific domains (e.g. replication, number of observations, and so on; Fig. 1). Notably, included in simulated data, were those emulating the results of high-throughput sequencing of microbial assemblages, as these are analytically challenging due to their dimensionality, high among-replicate variation, and extreme rank-abundance skew—often several microbial taxa are orders of magnitude more abundant than the numerous marginal taxa that typically compose the bulk of biodiversity within a sample (e.g., see Lynch and Neufeld 2015, Sachdeva et al. 2019). Our primary analytical goal was to measure the sensitivity and accuracy of DMM for comparing feature relative abundance between compositions and to compare the performance of DMM with competing approaches. Also, we provide a primer on the requisite algorithmic methods (e.g., VI and HMC) for Bayesian implementation of DMM and explore how different algorithms affect model accuracy and computational expense. Finally, we analyzed a data set published by Duvallet et al. (2019) that describes the lung microbiomes of children experiencing aspiration of foreign material and evaluated to what extent DMM recapitulated the published analyses or detected additional differences among microbiomes.

**Figure 1:**
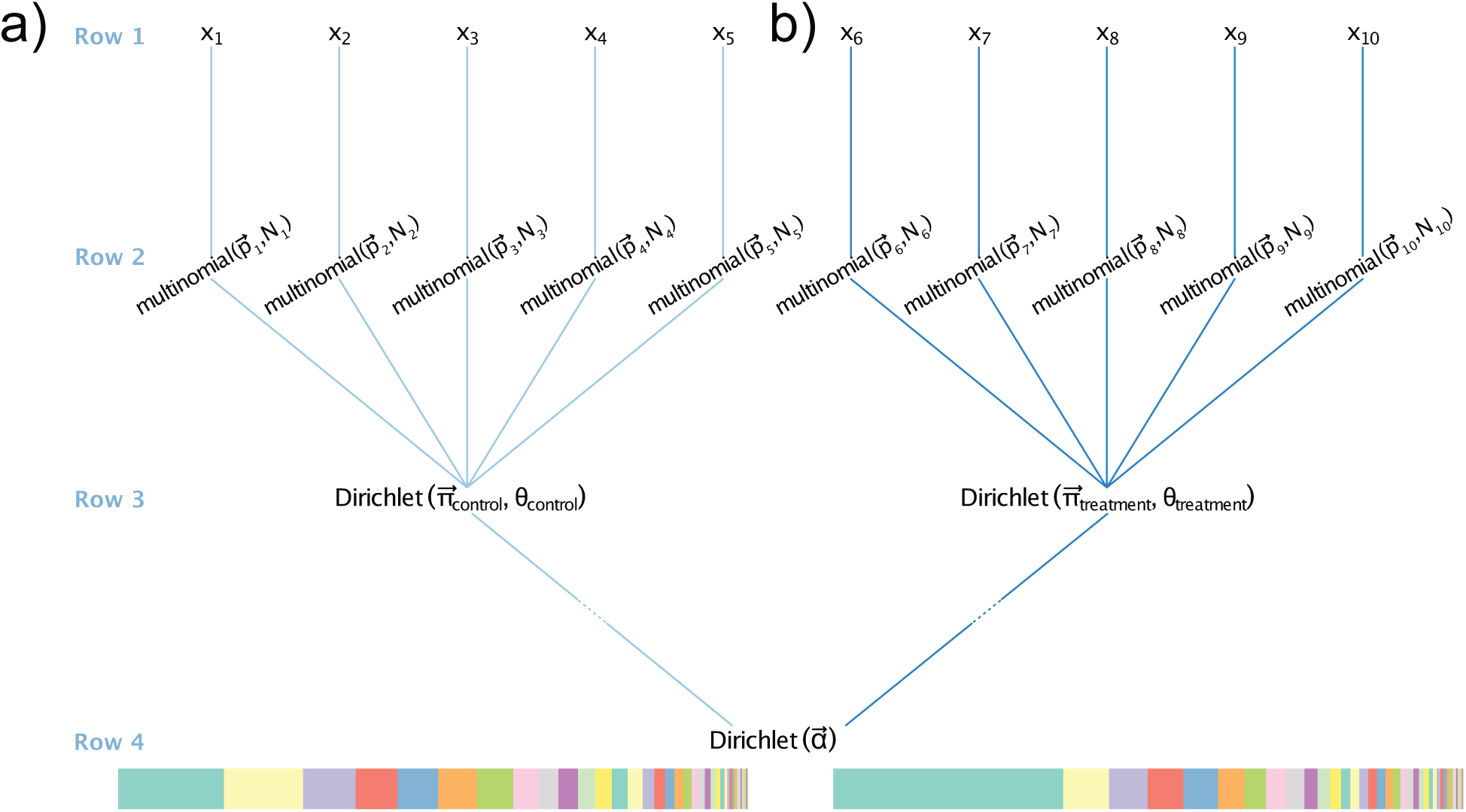
Visual depiction of hierarchical Bayesian modelling of the relative abundance of features within compositional data. Panels (a) and (b) represent two different sampling groups–a treatment group and a control group. The colored bars at the bottom of the plot show two hypothetical compositions that differ between those sampling groups. These compositions differ by virtue of the first feature, shown in pastel green, shifting dramatically in relative abundance, thus all other features are shifted in relative abundance as well (because these are proportion data and must sum to one). This interdependency represents an opportunity for statistical modelling because parameters that describe relative abundance are mutually informative. However, interdependency also poses many challenges (see main text). Replicates within sampling groups are denoted as *x*_*i*_, where *i* is in an integer in the range [1,10] (row 1). Replicates consist of data that are multinomially distributed (see Box 1). Therefore, each replicate is modeled using a unique multinomial distribution with parameters 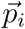 and *N*_*i*_ (row 2), where the vector 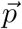 describes the probabilities that an observation would be assigned to a particular feature and the *N* parameters denote the total number of observations per replicate. Multinomial parameters are modeled as a deviate from a Dirichlet distribution unique to the treatment group (row 3). The 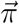 parameters of the Dirichlet are estimates of proportional abundance for all features within the group. The *θ* parameter is a scalar intensity parameter that describes the amount of among-replicate variation present within each sampling group. The prior imposed on the Dirichlet distributions of both sampling groups has the expectation 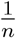 for each feature, where *n* is the number of features. If desired, additional Dirichlet distributions could be added between rows three and four to share information as dictated by more complexly nested experimental designs.

#### Box 1. A brief explanation of the multinomial and Dirichlet distributions

The multinomial distribution is the multivariate generalization of the binomial distribution. The binomial distribution can be used to describe counts of binary outcomes, with respective probabilities *p* and 1 − *p*. For instance, with a finite sample of observations, the binomial distribution would be useful for estimating the frequency of females (*p*) in a dioecious population. The multinomial distribution extends this concept to encompass more than two unique outcomes. For instance, a composition comprising three equally abundant features would have the the following multinomial parameter vector: 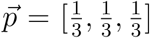. As an example, consider data from a sequencing machine. The counts of sequences that fall into each category (e.g., transcripts or taxa) are multinomially distributed, with a probability that corresponds to its relative abundance. For three equally abundant features (i.e. microbial taxa), there would be an equal chance of sampling a sequence from each of the features and on average we would expect to obtain the same number of sequences from each (for this example, we assume no laboratory-technique imposed bias).

To share information among samples in the same sampling group (e.g. treatment group, host population, or sampling location) and recover group-level estimates of the proportion of each feature in a composition, the Dirichlet distribution can be appropriately parameterized. The Dirichlet distribution is the multivariate generalization of the beta distribution. Deviates from a standard beta distribution fall in the range of [0, 1], and the distribution can be parameterized with expectation *π* (the expected frequency of the reference category, with 1 − *π* for the alternative category) and a parameter, *θ*, that affects the variation among deviates. Likewise, the Dirichlet distribution can be parameterized by a vector of expected frequencies of each feature 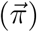, and an intensity parameter, *θ*. When drawing deviates from the Dirichlet distribution, the intensity parameter influences the amount of among-deviate variation in the frequencies observed—for a given 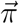, larger intensity parameters induce less among-deviate variation. This parameterization of the Dirichlet thus allows modelling of the variation among experimental replicates (the “noise” within the data).

Information about the frequencies of features within replicates 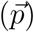 is shared to estimate frequencies for each feature within that sampling group 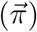, forming a hierarchical model (Fig. 1) that is analogous to how replicates can be used in an analysis of variance to learn about marginal, grand means associated with treatments. Estimates of frequencies of compositional features at the sampling group level 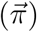 are the basis of inferences about which features differ among sampling groups (e.g., treatment versus control) and by how much (on an absolute or normalized scale).

## Methods

### Dirichlet multinomial modelling approach

Our specification of the Dirichlet-multinomial model generally follows that of Fordyce et al. (2011, implemented in the bayespref software) and takes as input a matrix of counts (X). The rows of this matrix correspond to different replicates (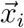; the superscripted arrow denotes a vector) and the columns correspond to features of the composition (the format of an OTU or transcript table). Each count *x*_*ij*_ in this matrix corresponds to the *j*^th^ feature (of *n* features in total) in the composition observed in the *i*^th^ replicate sample. Replicates are grouped into *k* groups, corresponding to treatment conditions, sampling locations, or some other stratification that specifies which replicates share information (parameters shared among replicates for the group). Counts in each row of the matrix are multinomially distributed:

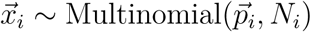

Each value *p*_*ij*_ in 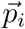 is the probability of observing a particular feature *j* in sample *i* and 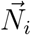 is a vector of the total counts in each sample. The product across *i* replicates of the *i* multinomial distributions forms the likelihood in the model and can be written:

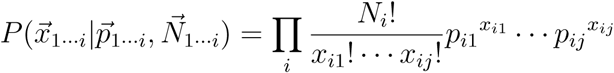

The prior probability for the vector of feature proportions 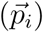 is a Dirichlet distribution, with parameters that are specific to the *k*^th^ group of replicates and that are learned from the data:

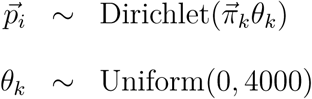

In this parameterization of the Dirichlet distribution for 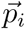, the 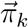 parameters correspond to the expected proportions of each of the *n* features (e.g., a particular transcript or taxon) in group *k*, and *θ* is an intensity parameter that is shared among all features (see Box 1). For a given 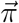, larger *θ* means less variation among deviates from the Dirichlet expectation 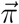. The probability density function of this distribution, across *i* replicates within the *k*^th^ group, is given by,

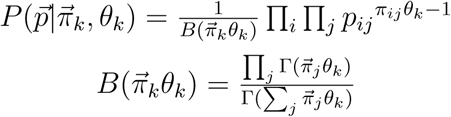

where 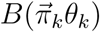 is a normalizing function that ensures the Dirichlet distribution integrates to one. The hyperprior for the 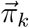 parameters at the “topmost”, or most inclusive, level of the model hierarchy is another Dirichlet distribution with equal prior probability for each feature within the composition. For this Dirichlet distribution we use *α*_1*…n*_ = 10^−7^ as a prior that will contribute little information, gives an expected value of 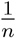, and has a high variance on the expectation:

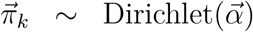

The overall model for the posterior distribution for parameters of a sampling group is:

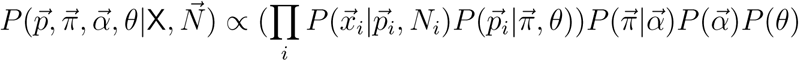

To quantify differences in proportions of features between two sampling groups (often referred to as “differential relative abundance testing”; Thorsen et al. 2016, Weiss et al. 2017), posterior probability distributions (PPDs) for *π*_*j,k*=1_ − *π*_*j,k*=2_ (Fig. 2d) can be obtained. Consistent with convention, if 95% of the samples of this PPD of differences are either greater or less than zero, then there is a high certainty of a non-zero effect of sampling group on feature relative abundance. One can also observe where zero occurs in the PPD of differences to quantify the probability of no effect of sampling group on feature relative abundance.

**Figure 2:**
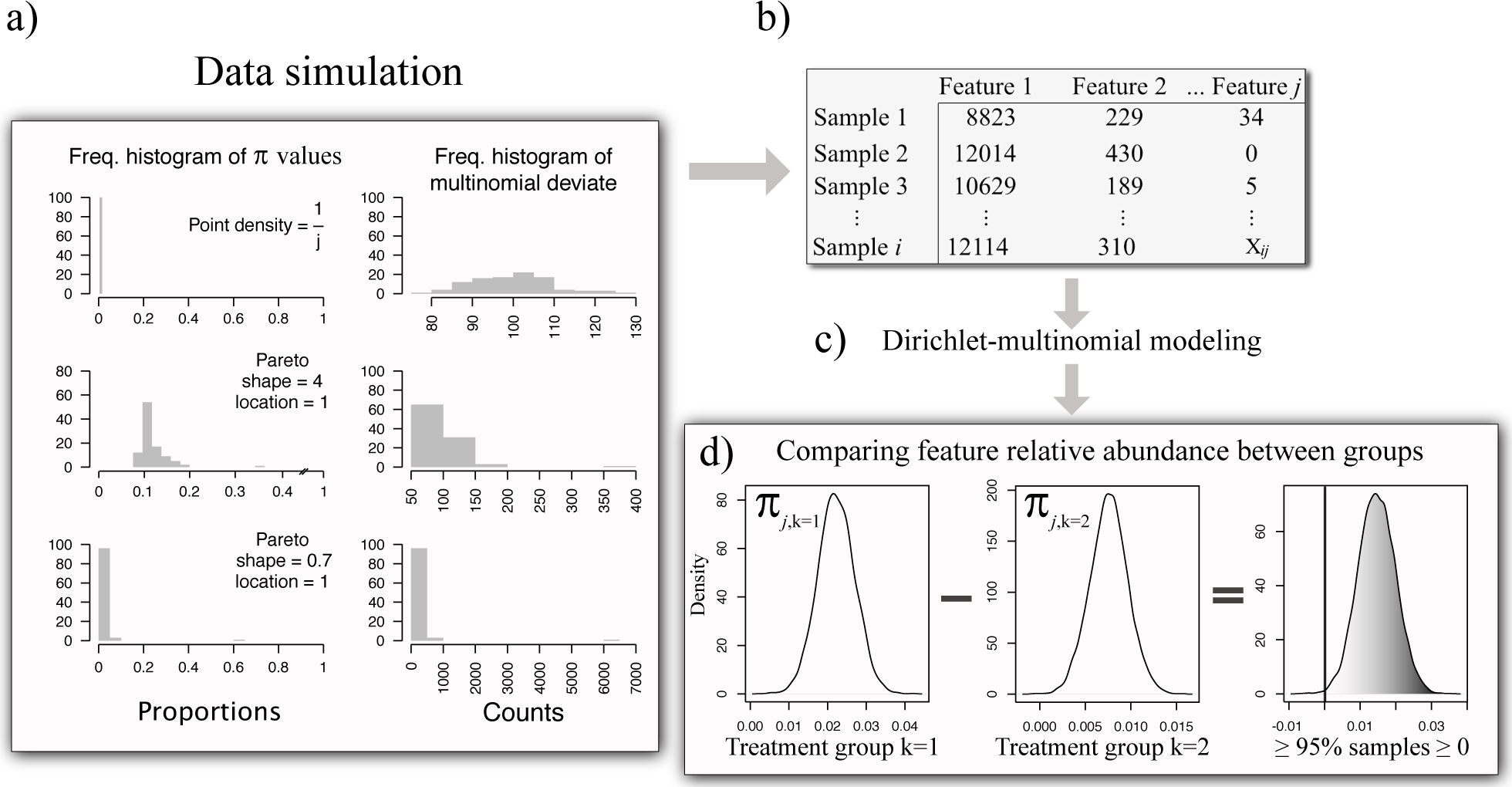
Visual description of simulation approach. (a) Deviates from either of two Pareto distributions, or a point density, defined as one divided by the number of features (*j*), were used to simulate values used to parameterize Dirichlet distributions. The use of these three approaches generated deviates with parameters that differed dramatically in rank abundance profiles, as shown in the left portion of panel (a). These deviates were, in turn, used to parameterize a Dirichlet distribution (with intensity parameter *θ*). A deviate of this Dirichlet distribution served as the parameter vector of a multinomial distribution that was sampled (b) to generate a feature (*j*) by replicate (*i*) matrix that emulated an OTU or transcript table (see the right portion of panel a for an example frequency distribution of multinomial deviates). This matrix encompassed samples belonging to two sampling groups. Dirichlet parameters for each group were made to differ such that certain features varied in relative abundance between groups by a known effect size. (c) Hierarchical Bayesian modelling (Fig. 1) was used to estimate the Dirichlet parameters (*π*_*j,k*_) describing the relative abundance of each feature (*j*) in each sampling group (*k*). (d) To determine if a feature (*π*_*j*_) differed in relative abundance between treatment groups (*k*), the posterior probability distribution (PPD) for the feature of interest from one treatment group, *π*_*j,k*=1_, was subtracted from the PPD for that feature from the second treatment group, *π*_*j,k*=2_. If the resulting PPD of differences indicated zero difference was improbable, then there was high certainty that *π*_*j*_ differed between treatment groups. Additionally, the location of zero within the PPD quantified the certainty of a non-zero effect of treatment.

If a sampling scheme was used that induces dependence among replicates via a more nested hierarchical structure then the model described above, then the model hierarchy could be extended to include inference of the Dirichlet distributions describing the relative abundances of features within each additional stratum of the sampling scheme. For example, consider a study design where subjects are provided one of several diets and gut microbiome samples are taken from both sexes. In this case, one would want to account for non-independence among the data due to both sex and diet treatment. This can be accomplished through incorporation of additional Dirichlet distributions into the model, 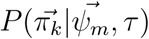, where 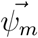 describes the relative abundances of features within each diet treatment (*m*), *τ* is the intensity parameter for that Dirichlet distribution, and 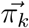 describes relative abundances of features within each sex that is nested within each diet treatment. In this way, the model can be extended to encompass as many hierarchical layers as desired, given suitable sampling and replication (Coblentz et al. 2017).

#### A primer of the algorithms to perform DMM

One goal of statistical modelling is to estimate values for parameters that could correspond with directly observable variables (i.e. the data) or with latent, unobservable, variables (i.e. those that are inferred from observable variables). Bayesian modelling attempts to estimate parameters of interest, while explicitly quantifying the uncertainty in those estimates and allowing for the influence of prior knowledge on estimates. Much of Bayesian statistical modelling relies on Markov chain Monte Carlo (MCMC) sampling (Gelman et al. 2013). A Markov chain is a series of states where each state depends upon the immediately preceding state. Monte Carlo refers to repeated, random sampling. MCMC is a process by which values are suggested randomly from a probability distribution and substituted into the functions that define the model. Over MCMC iterations, sampling converges on the most supported parameter space (the PPDs for model parameters) and samples in the chain occur with probability defined by the PPD.

There are several MCMC algorithms and they primarily differ in how they choose or propose new values and their criteria for inclusion of those values in the chain (Gelman et al. 2013). A standard MCMC tool is the Metropolis algorithm (Gelman et al. 2013, pg. 289). To perform Metropolis sampling, a value (*x*_*t*_) is proposed from some distribution *Q*(*x*_*t*_|*x*_*t*−1_), where *t* is iteration (a suitable initial value, *x*_0_, is required). Once *x*_*t*_ is chosen a ratio of 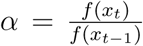 is calculated, where *f* (*x*) is a function that is proportional to the probability density to be estimated. The new value *x*_*t*_ is accepted into the chain with probability *α*, otherwise *x*_*t*_ = *x*_*t*−1_. The Metropolis algorithm relies on a symmetric proposal distribution, such that *Q*(*x*_*t*_|*x*_*t*−1_) = *Q*(*x*_*t*−1_|*x*_*t*_). The Metropolis-Hastings (MH) algorithm extends this concept through relaxing the assumption of symmetry regarding the proposal probability distribution.

Gibbs sampling (Geman and Geman 1987, Kruschke 2015) is a special case of the MH algorithm (because the proposal acceptance criterion is always met; see pg. 289 in Gelman et al. 2013) and is suited for cases when the distributions used within the model are conditionally conjugate, such as when the prior and likelihood distributions are conjugate and, consequently, their product has a well defined form. At each iteration of Gibbs sampling (*t*), each parameter is sampled from the conditional distribution defined by the other parameters in the model, which are held constant at values chosen at iteration *t* − 1. Parameters are typically updated one at a time, in a predefined order.

The probabilistic programming language JAGS (Plummer 2003) implements Gibbs and Metropolis-Hastings MCMC as required to obtain samples from the distributions in our DMM. Henceforth, we refer to parameter estimation via Gibbs, Metropolis, and Metropolis-Hasting sampling as MCMC. These algorithms can be slow to converge for complex models; indeed in our experience, in a JAGS implementation, convergence may not be observed for the majority of parameters over a week of runtime for DMM with high dimensional data (such as transcriptomic data), even with sensible chain initialization values (a bespoke software implementation of MCMC tuned to the data and model would likely be faster, but would require greater care in programming and use).

Hamiltonian Monte Carlo (HMC) seeks to improve upon the efficiency of MCMC through the use of a physics inspired algorithm (for an excellent description of HMC see Monnahan et al. 2017). The sampling method can be envisioned by considering a ball dropped into a bowl and allowing the ball to roll about the curvature of the bowl. The bowl is the PPD and is frictionless, so the ball will roll back and forth in the bowl forever. After repeated drops of the ball into the bowl, from different angles and with different potential energies, the shape of the PPD is determined from the combined paths the ball took across all iterations. The benefit of this approach is that samples from nearly anywhere in the PPD can be generated at each iteration (HMC does not use a Markov chain process, but does rely on a Metropolis ratio to determine acceptability of updates), whereas MCMC typically chooses values based on the previous state space and thus cannot quickly move throughout the PPD, which can slow chain mixing and time to convergence. The probabilistic programming language and software Stan allows the use of an improved version of HMC called the “no U-turn” sampler that avoids redundant sampling of parameter space (Hoffman and Gelman 2014). To continue the previous analogy, when the ball starts to make a U-turn due to the curvature of the bowl, the sampler is stopped, and the ball dropped again—thus avoiding spending sampler time in previous explored parameter space.

HMC often improves model runtime (Monnahan et al. 2017) over MCMC, but can still be quite time consuming. Variational inference is a class of optimization methods from the machine learning literature that can rapidly approximate PPDs (Blei et al. 2017), and thus holds great promise for statistical modelling of complex data where the speed of MCMC or HMC is insufficient. Variational inference (VI) has yet to be widely applied by biologists, but it has been used to estimate population genetic structure (e.g. Raj et al. 2014, Scordato et al. 2017), genotype-phenotype associations (Carbonetto and Stephens 2012, Logsdon et al. 2010), phylogenetic relationships (Jojic et al. 2004), and in a generalized latent linear modelling context (Niku et al. 2019).

The idea behind VI is that the exact PPD need not be estimated, but can be approximated through optimization of parameters of more tractable distributions. Briefly, a density is chosen from a family of distributions and optimized so that the Kullback-Leibler (KL) divergence between that density and the PPD is minimized. KL divergence relies on the definition of entropy. Entropy is a measure of the information present within a distribution and can be expressed (for a discrete probability distribution):

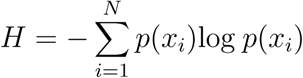

where *p*(*x*) is a function that outputs a probability contingent upon an input value *x*, which is indexed by *i*. It is perhaps easiest to intuit entropy using log_2_, in which case *H* is the minimum number of bits needed to encode the data. KL divergence extends this idea to quantify the amount of information necessary to explain the divergence (||) between two probability distributions *p* and *q*, which, in this example, are discrete:

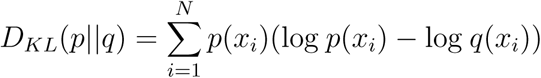

Because this measure of divergence is based on the quantification of entropy, when *p* and *q* differ greatly, then more information is required to explain how they differ and KL divergence increases. For VI we wish to minimize the KL divergence between the probability distribution 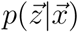 and some density 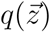 chosen from a family of distributions *Q*. To avoid computation of 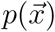 (see Blei et al. 2017, for more), minimizing the KL divergence can be solved by maximizing the “evidence lower bound” (ELBO; the 𝔼 used below refers to expectation):

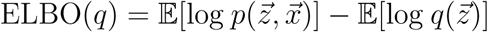

The ELBO is the negative of KL divergence after adding the constant log *p*(*x*). Thus maximizing the ELBO is equivalent to minimizing the KL divergence, up to the added constant. This also means that:

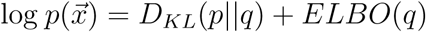

The ELBO describes the lower bound of the evidence, because when the ELBO is subtracted from the evidence (log *p*(*x*)) the result must be ≥ 0, because KL must be ≥ 0. Because maximizing the ELBO does not require computing log 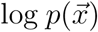 it is easier than minimizing KL divergence. Maximization techniques can then be used to find the density 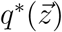 that best approximates 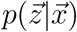.

Choosing *Q* such that the family of densities includes a 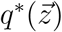 that provides a good approximation, while being easily optimized, is the challenge of VI. Stan solves this problem through a method called “automatic differentiation variational inference” (Kucukelbir et al. 2015) by first transforming the data that are the support of the latent variables to lie within the real numbers (ℝ) and then suggesting a Gaussian distribution, which can be optimized to fit the data, and which induces a non-Gaussian approximation to the untransformed data. Stan’s default approach uses the “mean-field” algorithm, which treats latent variables (*z*_*j*_) as independent and assigns a unique density, *q*_*j*_(*z*_*j*_), to each of these *j* variables. Since Stan transforms the data such that latent variables have support on ℝ and then fits Gaussian distributions to those data, this statement becomes the product of many Gaussian distributions, each of which are optimized to minimize the ELBO. Following the notation of Blei et al. (2017), this can be written:

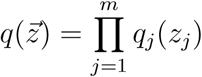

VI is an attractive technique because it can be many orders of magnitude faster than MCMC (e.g. Raj et al. 2014). However, it is unclear how well VI works across analytical tasks and model specifications (Blei et al. 2017).

### Model implementation

We performed DMM in the R statistical computing environment (R Core Team 2019) using models specified for the JAGS and Stan (Carpenter et al. 2017) software programs, and used the models through the rjags (Plummer 2015) and rstan (Stan Development Team 2018) R packages, respectively. JAGS uses MCMC (Gibbs and MH), whereas Stan implements HMC (no U-turn sampling) and VI. Model specification for use in Stan was slightly modified from that described above in that we used an exponential distribution as the form of the prior for *θ*_*k*_:

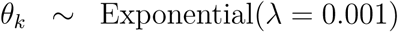

This change in model specification followed the recommendation to avoid uniform priors provided in the Stan documentation.

For HMC and MCMC implementations of DMM, we used two chains to explore parameter space. Initial values for 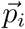 in each chain were the vector of proportions observed from the data in replicate *i*, and values for 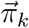 were initialized using the vector of observed proportions for each feature across replicates within *k* (i.e., the maximum likelihood estimates for 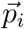 and 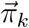). *θ* was left to be initialized internally by rjags and rstan. In rjags, the model was subjected to an adaptation period long enough for the sampler to approach optimal efficiency as determined via internal heuristics, or for 20,000 iterations, whichever came first. Models were updated (“burned in”) for 300,000 steps for rjags and 1000 steps for rstan (with a maximum tree depth of 10). This discrepancy in burn in time was needed because in preliminary work we observed much quicker convergence with HMC than MCMC sampling. We obtained 1000 samples from PPDs by saving every second sample for HMC, and 2000 samples from PPDs for MCMC by saving every fourth sample.

Preliminary inspections of samples showed higher auto-correlation of parameter estimates for MCMC sampling, hence we discarded more samples (higher thinning rate) from the MCMC-derived chains. MCMC convergence was evaluated via the Gelman-Rubin and Geweke statistics (Geweke 1991, Gelman and Rubin 1992). We note that the runtime of MCMC could likely be improved by optimizing adaptation, burn in, and sampling steps within JAGS, or by implementing a custom MCMC procedure in the C (or an equivalent) programming language. Data with different dimensions and variance among samples would likely require different optimizations, so we have not further pursued optimization of the MCMC herein. To perform variational inference we used the functionality included within Stan (the “vb” function; Kucukelbir et al. 2015) and collected 1000 samples from the estimated posterior distributions.

The ability of models to recover true simulation parameters was estimated via root mean square error (RMSE) and the percentage of times the true simulation parameters were within the 95% high density intervals (HDIs) of PPDs (as per Kruschke 2015, pg. 727). For unimodal, symmetric PPDs, the HDI and equal-tailed probability interval should be identical (Gelman et al. 2013, pg. 38). We measured model bias as the average difference between estimated parameters and the truth and we measured model precision as 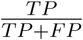, where TP refers to true positives and FP to false positives. False positive rate was calculated as 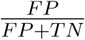, where TN is true negatives. Additionally, we calculated Matthew’s Correlation Coefficient (MCC; Matthews 1975), which provides a measure of classifier performance in terms of both true and false positives and negatives. MCC is the correlation between actual and predicted classifications and varies from one (perfect classification) to negative one (completely incorrect classification). An MCC value of zero denotes a classifier that performs no better than expected from random guessing.

### Data simulation

To evaluate the performance of DMM implementations and alternative statistical methods (see below), we simulated and analyzed data with two sampling categories (*k*), corresponding to treatment and control groups, or some other blocking factor of interest (Fig. 2). We simulated data that possessed three different rank abundance profiles that were meant to correspond to the variety of data encountered by practitioners (Fig. 2). We considered simulations in which all features were equally abundant 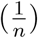, and two sets of simulations in which features were sampled from Pareto distributions with differing shape parameters. The Pareto distribution describes data with few abundant features and many rarer features (Krishnamoorthy 2006). The skew towards low abundance in this distribution is controlled by the shape parameter, with smaller parameters increasing skew (Fig. 2); the location parameter defines the minimum value of the distribution. For each simulation, we sampled one of these distributions to populate a vector 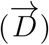 of length corresponding to the approximate desired number of features (*n*) within the simulated data:

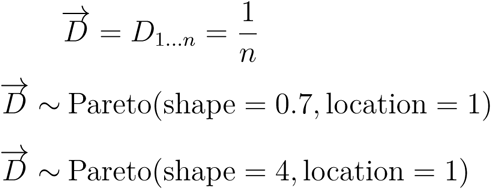

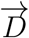 was duplicated to make a second vector, 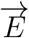. Selected features within these vectors were multiplied by an effect size (either 1.1, 1.5, or 2, to simulate 10%, 50%, or 100% shifts in feature relative abundance), such that those elements differed between 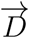 and 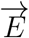. Features that varied between vectors were chosen randomly from within each of three broad abundance classes (abundant, rare, and intermediate; see Electronic Supplementary Material) present within 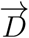 and 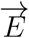. Only features of intermediate abundance were available when constraining all relative abundances to be equal. Effect sizes were applied so that 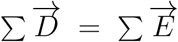. These two vectors were multiplied by a specified intensity parameter *S* and used as the parameters for two Dirichlet distributions that were sampled to create 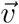 parameter vectors for multinomial distributions corresponding with each replicate. In this way, we simulated a replicate by feature matrix where replicates were split into two treatment groups and known features differed between treatment groups. Simulated data sets often had fewer features than the originally specified value for *n*, because when drawing deviates from multinomial distributions with many rare features, all features would not be observed in each deviate (for a visual depiction of simulation approach see Fig. 2).

Using this approach, we simulated data from each sampling distribution that varied in dimensionality (number of features, ∈ {500, 2000}), number of replicates (∈ {10, 50}), the total number of observations per replicate (e.g., the number of reads per sample for sequencing data; ∈ {10000, 50000}), the variation (noise) among replicates (∈ {0.5, 3}; the intensity parameter in notation provided above), and the effect size applied to features that differed between sampling groups (∈ {1.1, 1.5, 2}; to apply the effect size transformation, these values were multiplied by the original proportion. In total, we created and analyzed 144 data sets. Because the same number of observations were used for each replicate, transformation of the data to account for unequal sampling effort was not required. After simulating data matrices, we added a one to every datum, and thereby avoided numerical errors in JAGS that arise with Dirichlet parameters approaching zero.

For our main simulation, we did not vary read counts among replicates for the sake of simplicity, however to ensure that this did not bias our results we simulated data where replicates differed by up to two orders of magnitude in total observations (read count). To accomplish this, multinomial deviates were obtained as described above, however the total number of draws from the multinomial distribution was randomly selected from ∈ {1000, 10000, 100000}. Data used for this additional analysis were simulated using a representative subset of the aforementioned attributes. Additionally, to better understand the false positive rate of DMM, we simulated and analyzed data where no features were expected to differ between treatment groups, again using a representative subset of the attributes presented above to simulate data.

We competed our implementations of DMM against ALDEx2 v1.14.1 (Fernandes et al. 2014), ANCOM v2.0 (Mandal et al. 2015), DESeq2 v1.18.1 (Love et al. 2014), edgeR v3.20.9 (Robinson et al. 2010), mvabund v4.0.1 (Wang et al. 2019) and a frequentist approach using repeated Wilcoxon rank sum tests with a Benjamini-Hochberg false discovery rate (FDR) correction (Weiss et al. 2017). We used multiple comparison correction and typical settings for all software (see the Supplemental Material). Of the aforementioned methods, only ALDEx2 relies upon DMM. ALDEx2 estimates posterior probability distributions of Dirichlet parameters, which are subsequently transformed via the centered log ratio (Aitchison 1982). Transformed MCMC samples are subjected to a frequentist test of differential relative abundance between sampling groups, *p* values calculated, and the distribution of *p* values across MCMC samples obtained (with multiple comparison correction applied as desired by the user). The mean of this distribution is used as a point estimate of the significance of treatment. mvabund relies on a generalized linear model, in our case using a negative binomial distribution, to determine differential relative abundance. Each feature in the simulated data was a response variable and treatment group was the categorical predictor variable in the model. If the effect of the predictor was significant then the feature differed between treatment groups in relative abundance. mvabund is thus quite similar to edgeR and DESeq2, however those methods use different normalization strategies.

Our implementation of DMM differs from these methods in several important ways: 1) most competing methods do not rely on the Dirichlet and multinomial distributions, which explicitly model compositions (except ALDEx2); 2) we use a more complex hierarchical structure than the other methods tested to share information among replicates and sampling groups; 3) we do not perform repeated frequentist tests to determine differences in feature relative abundance, but instead directly subtract posterior probability distributions for parameters of interest and observe the location of zero in the resulting distribution of differences.

For all methods, we evaluated how data attributes (e.g. number of replicates, features, etc.) influenced model performance via multiple regression, with either the proportion of true positives recovered or false positive rate as the response variable.

### Analyses on empirical data

To understand how DMM could affect inferences made using previously published, empirical data, we analyzed data from Duvallet et al. (2019) describing the lung microbiomes of children with and without oropharyngeal dysphagia (swallowing difficulties) induced aspiration (when a foreign substance enters the lungs). These authors characterized the bacterial assemblages in the lungs (obtained via bronchoalveolar lavage; BAL), gastric fluid, and oropharyngeal region (OR) of each subject via sequencing of the 16S locus. Aspiration is linked to pneumonia in both adults and children (Holas et al. 1994, Marik 2001, Thomson et al. 2016), but the provenance of aspirated microbes is poorly understood. Duvallet et al. (2019) showed that the lung microbiome of patients with difficulty swallowing is more similar to the microbiome of the oropharyngeal region than that of gastric fluid. These authors performed differential relative abundance testing using Kruskal-Wallis tests with a multiple comparison correction to determine whether certain bacterial taxa shifted in relative abundance between aspirating and non-aspirating patients. The authors did not find any taxa that differed in relative abundance, regardless of substrate examined (BAL, gastric fluid, or OR), though they did detect shifts in prevalence (presence across subjects within a sampling group) with phenotype, and suggested that microbial exchange between the lungs and oropharyngeal region is greater than between the lungs and stomach. Using DMM (both VI and HMC; implemented as described above) and all aforementioned competing analyses, we reanalyzed the publicly available BAL data from aspirators and non-aspirators. The data we analyzed were obtained from 66 patients (33 aspirators, and 33 non-aspirators) and included 4006 OTUs (for details of sequence processing see Duvallet et al. 2019).

## Results

Dirichlet-multinomial modelling (DMM) provided a good compromise between true positive recovery and false positive generation (Fig. 3 & S2), as shown through analysis of data simulated in the context of a treatment-control experimental design. DMM consistently detected many more true positives than competing methods (Fig. 4) and this sensitivity facilitated detection of subtle shifts in relative abundance between sampling groups. For instance, when analyzing data with a skewed rank abundance profile, DMM detected approximately 15–20% of features that were shifted by treatment by just 10% of their relative abundance. None of the other methods that we employed were able to reliably detect these subtle effects (Fig. 3). When effect sizes were larger, DMM recovered more than 80% of true positives on average, which was 20–40% more true positives than were recovered by DESeq2, the next best model in terms of sensitivity.

**Figure 3:**
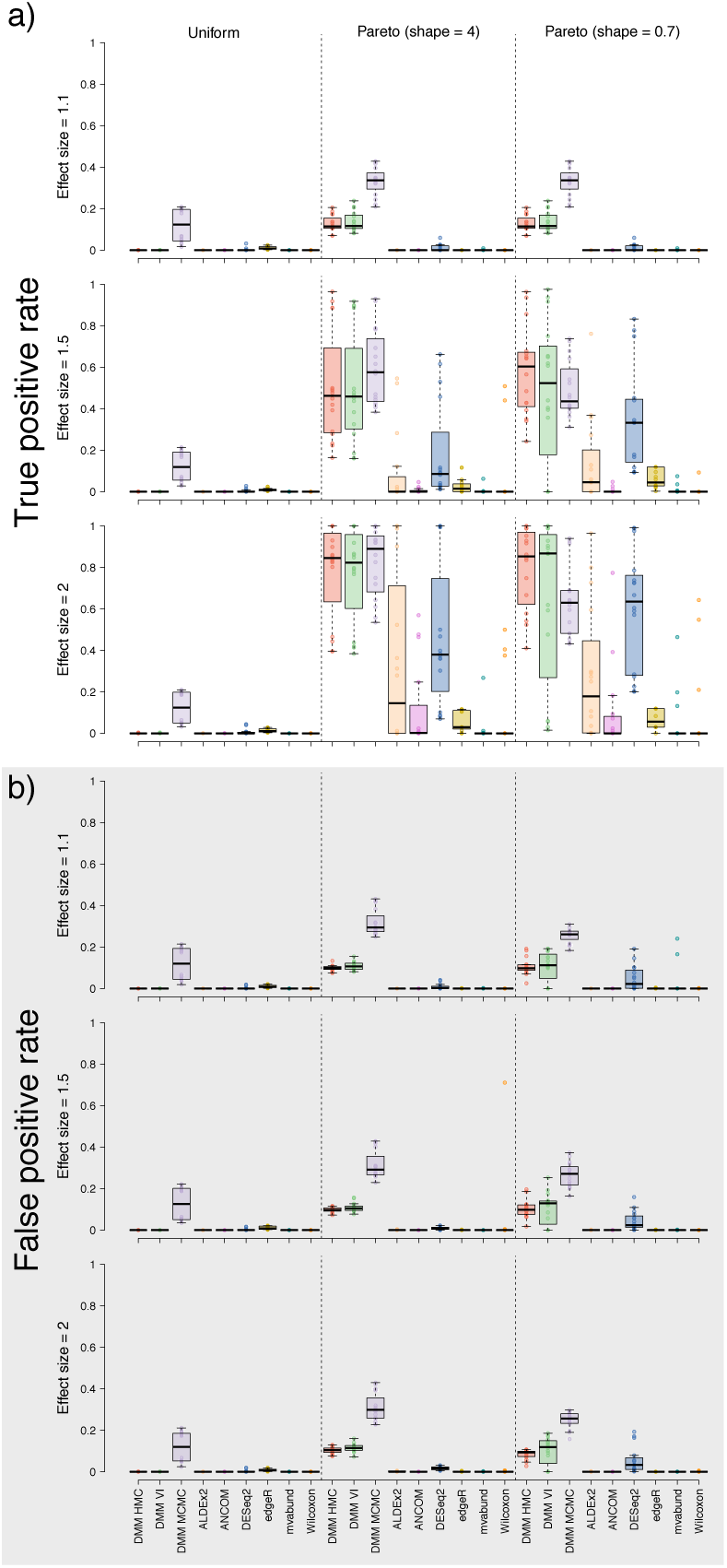
Performance of Dirichlet-multinomial modelling (DMM) and competing methods when confronted with simulated data from a treatment-control experimental design. Each point denotes the results from analysis of a simulated dataset. Panel a depicts true positive rate and panel b depicts false positive rate. The x axis describes the methods competed, which are each given a unique color. Each panel is split into three sections that correspond with the three rank abundance profiles used to simulate data (see Fig. 2). “Uniform” refers to data where the expected relative abundance of all features was equivalent; “pareto (shape = 4)” refers to data with an intermediate rank abundance skew; “pareto (shape = 0.7)” were highly skewed data with very few abundant features and many rare features. Features were made to shift in relative abundance between treatment groups by different effect sizes (an effect size of 1.1 corresponded with a 10% shift in relative abundance). Panels are split by row to show results for a specified effect size. Rectangles in the boxplots delineate the central 50% of the data (1st to 3rd quartiles, also called the interquartile range) and contain the median (delineated by a horizontal line). Whiskers extend an additional 1.5 times the interquartile range beyond the first and third quartiles. These are the defaults for boxplots in base R.

**Figure 4:**
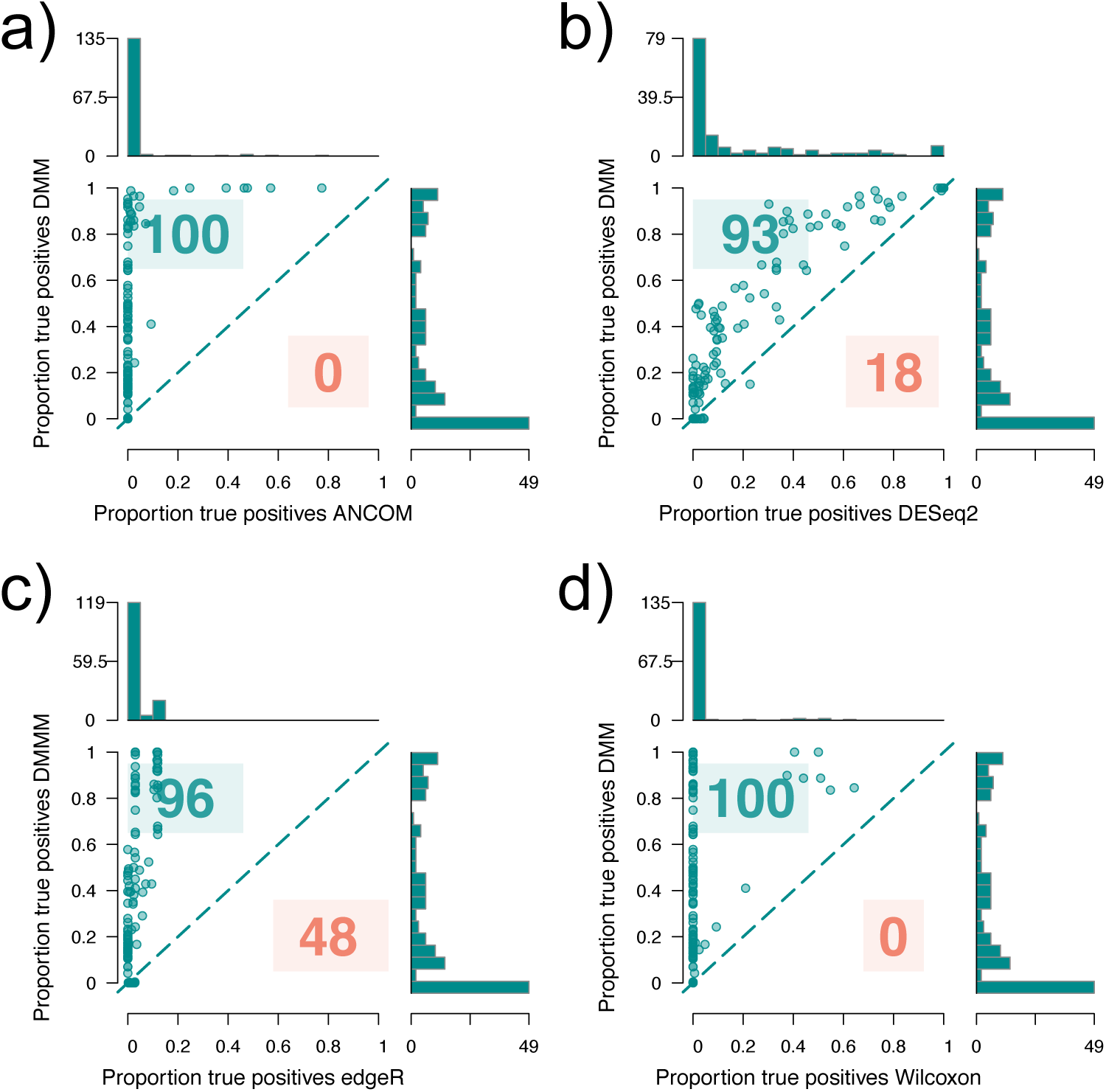
Relative ability of competing methods to detect true positives within simulated data. Each point represents the results from a simulated data set and each panel compares the proportion of true positives identified by Bayesian Dirichlet-multinomial modelling (using HMC) to the proportion of true positives identified by a competing method: (a) ANCOM, (b) DESeq2, (c) edgeR, (d) Wilcoxon rank sum test. The line bisecting each plot denotes equal performance of both models—so if a point lies above this line then HMC detected more true positives than the competing method for that data set. The summed numbers of points on either side of this line are shown to demonstrate relative performance of methods across datasets. For instance, in panel (a), the Dirichlet-multinomial model (DMM) detected more true positives than ANCOM for 100 data sets, while ANCOM was the more sensitive model for zero data sets. The sum numbers of simulations for each panel differ (and do not always reflect the 144 total data sets analyzed) because in some cases both DMM and the competing method exhibited equal performance. This was mostly the case for extremely challenging data when neither method was able to detect any true positives. Marginal histograms in each plot denote frequency distributions of results along the parallel axis.

The sensitivity of DMM came at the cost of a slightly higher false positive rate and a loss of precision compared to other methods (Figs. 3 & S1). Precision was generally high for uniformly distributed data and when the effect size that described the shift in relative abundance of a feature was large, however for data with skewed rank abundance profiles the precision of DMM was lower than competing methods. When considering the Matthew’s correlation coefficient (MCC), DMM typically performed as well or better than competing approaches examined (Fig. S2). MCC is a more holistic index of classifier performance than precision because it encompasses true and false positives and negatives. mvabund, ANCOM, and, for some data sets, Wilcoxon tests also performed quite well by this metric.

We observed that the FPR was adversely affected by the rank abundance skew within the data. Analysis of data that was simulated such that no features were expected to differ among treatment groups revealed that for data simulated from a uniform distribution FPR was negligible (0%, Fig. S3). However, FPR for HMC increased to 5.4% on average for data simulated such that they had a highly skewed rank abundance profile (Pareto shape parameter of 0.7). When data were of intermediate skew (Pareto shape of 4) then FPR increased to 8.2%. We also found that high among-replicate variation in sampling depth tended to increase FPR by a few percentage points (Fig. S4). On average, FPR of VI was only slightly higher than HMC. By comparison, FPR was often much higher when DMM was implemented via MCMC. Indeed, in many cases, MCMC generated an unacceptably high FPR of over 20%. This high FPR is at least partially due to the lack of convergence we observed for many parameters when using MCMC, even when we employed lengthy run times. We observed broadly comparable results from our primary simulation experiment, which spanned data with a broader variety of attributes and for which features differed in relative abundance among sampling groups (Fig. 3).

Of the analytical tools examined, DESeq2 and edgeR were the next most sensitive behind DMM. DESeq2 maintained a lower false positive rate than DMM. ANCOM, ALDEx2, and Wilcoxon tests all exhibited negligible false positive rates, but were only able to identify a small fraction of the features that shifted in relative abundance between sampling groups. All methods, including DMM, performed poorly when confronted with data where all features were equally abundant (denoted as “uniform” in figures). This was unsurprising, because, for these data, the expectation of *π* was approximately one divided by the number of features present and large, marginal shifts in relative abundance between sampling groups (such as doubling) still resulted in very small differences in proportions (e.g. 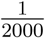 versus 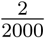), which were difficult to estimate.

We used multiple regression to test how data attributes influenced true positive detection and false positive rate (Tables S2, S3). For all methods competed, the degree of rank abundance skew within the data had, by far, the largest effect on model performance. Surprisingly, all methods were quite insensitive to variation in other data attributes. Data dimensionality (number of features), number of replicates, number of observations, and among-replicate variation had very minor influences on true positive detection and false positive rate for most methods tested (Tables S2, S3).

While our primary goal was ascertaining the relative merits of DMM for detecting differences in feature abundance, we also asked how well DMM could recover the relative abundances (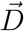 and 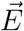) that were used to simulate data. We report very low average root mean square error (RMSE) for estimates of simulated relative abundances (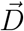 and 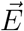) obtained through DMM (Fig. 5). As a complementary test of model performance, we determined how often the parameters used to simulate data fell within the high density interval (HDI) of PPDs. When feature relative abundances were equal, or modestly skewed (“Equal” or “Pareto, shape = 4”), the HDI of PPDs encompassed the value used to simulate data for nearly all parameters of interest, regardless of estimation method employed (MCMC, VI, or HMC; Fig. S5). Parameter estimation was much more difficult for highly skewed data—when using MCMC or VI, the true values for the parameters did not lie within the estimated HDIs in some cases. By comparison, HMC did better when confronting these challenging data—on average ∼90% of simulation parameters fell within the HDI, though there was wide variation in model performance depending upon data set (Fig. 3). We observed that the width of credible intervals for *π* parameters was not associated with relative abundance regardless of implementation method or dataset (Fig. S16– S18). Bias of DMM differed among implementations, with HMC having negligible bias (Figs. S7, S8, S9) and VI and MCMC exhibiting comparatively more bias. We observed that, for all implementations, bias, when present, was typically limited to the most abundant and rarest features within the dataset. Specifically, *π* parameters were occasionally slightly underestimated for abundant features and overestimated for rare features. This pattern was more noticeable for highly skewed data and can be explained given the prior we used for *π* parameters, which corresponded to 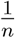, where *n* was the number of features. For skewed data with high among-replicate variation, the strength of the prior was not overcome by the likelihood, thus leading to slight overestimation of marginal features and underestimation of abundant features. If among-replicate variation was reduced, then DMM was able to accurately recover true parameters even for highly skewed data. The prior we chose was agnostic to rank-abundance curves and thus suitable for a wide-range of applications, but could be substituted for a prior with a specific rank-abundance profile if desired by the user.

**Figure 5:**
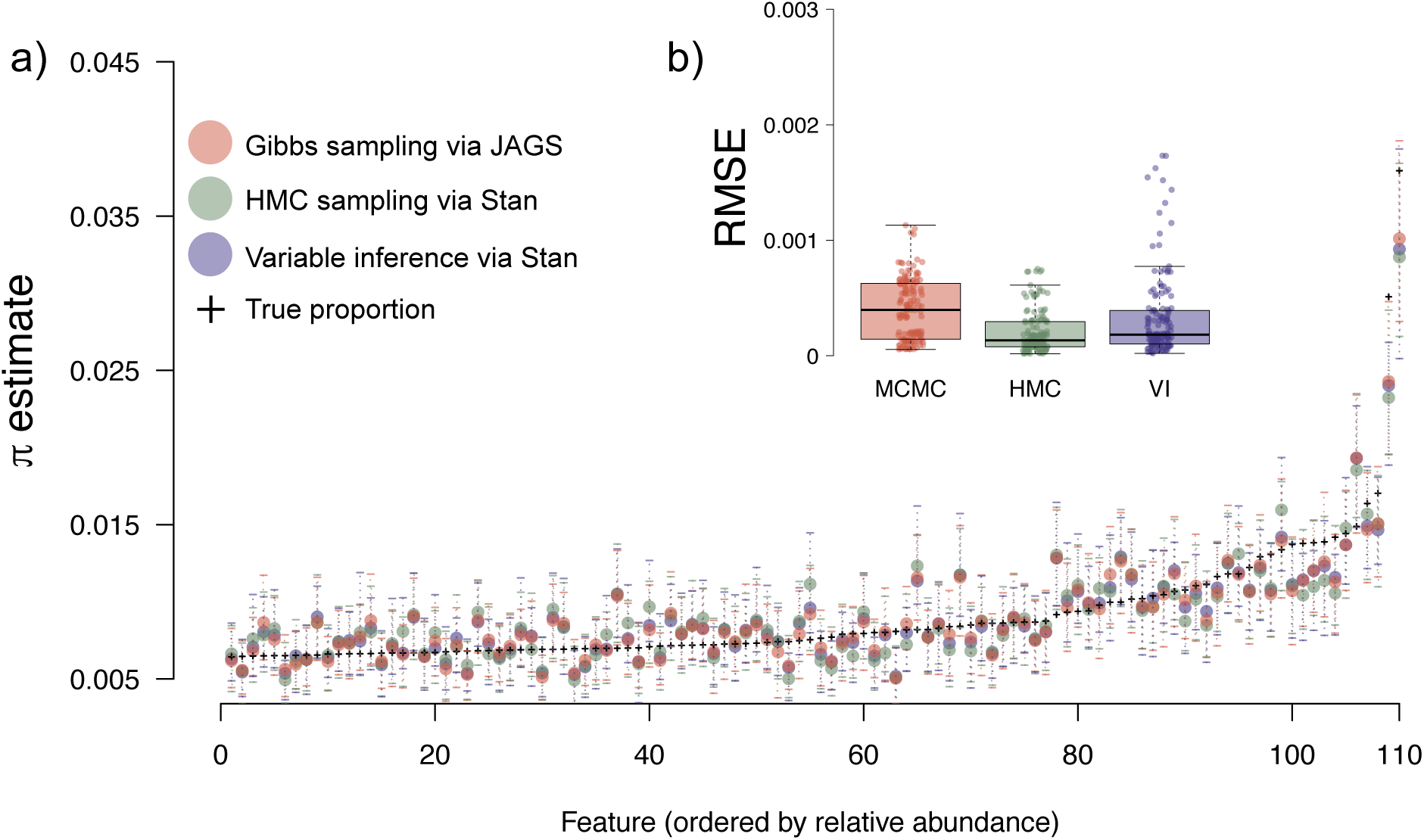
Comparison of DMM performance when using different methods to estimate posterior probability distributions (PPDs) of parameters describing feature relative abundance within sampling groups (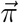; see Fig. 1). The results shown in panel (a) are from a single, illustrative simulation. Features are indexed along the horizontal axis and associated 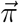 estimates are shown on the vertical axis. The means of PPDs are shown as shaded circles and the 95% high density interval (HDI) of the PPD is delineated by dotted lines. The true proportions (+ symbols) fall within the HDI of the PPD for almost all features, regardless of PPD estimation method. Average root mean square error (RMSE) for *π* parameters for all simulated data sets for each method is shown in panel (b).

### Inferences on empirical data

Reanalysis of data provided by Duvallet et al. (2019) demonstrated the sensitivity of DMM. Using HMC, we found that 53 taxa within the lung microbiome (samples were obtained via bronchoalveolar lavage) shifted in relative abundance between aspirating and non-aspirating children (Fig. S19). This contrasts dramatically with the results we obtained from repeated Wilcoxon tests with a Benjamini-Hochberg false discovery rate correction, mvabund, and ALDEx2, which suggested no taxa significantly shifted in relative abundance between sampling groups. By comparison, DESeq2 suggested 17 taxa differed, edgeR suggested ten taxa, and ANCOM four taxa.

Analysis of lung microbiome data using VI and HMC based implementations of DMM provided largely similar results; however, VI did report five fewer taxa shifted in relative abundance than did HMC. The majority of taxa identified by HMC were also identified by VI; the two methods did not agree regarding true positive status for only nine taxa. Of the 53 taxa that we found shifted between sampling groups, the most dramatic change was in a *Streptococcus* taxon, which was much more abundant in aspirating children (Fig. S19). An increase in this taxon has previously been reported in adult humans with pneumonia by Akata et al. (2016). We also found an increase in *Haemophilus* (Norman M. Jacobs and Harris 1979), *Moraxella* (Claesson and Leinonen 1994), *Neisseria* (Johnson et al. 1981), and *Prevotella* (El-Solh et al. 2003), all of which have previously been associated with pneumonia (see citations for examples), but may be present in healthy lung tissue as well (Beck et al. 2012). We also observed an increase in *Enterobacter, Lactococcus, Leuoconostoc*, and *Acinetobacter* taxa in the lungs of non-aspirating subjects.

## Discussion

Over the past decade, there has been considerable discussion regarding how molecular ecologists should process and analyze compositional data, particularly those generated by high-throughput sequencing instruments (e.g., see Knight et al. 2018, Thorsen et al. 2016, Weiss et al. 2017). This dialogue has been motivated by the constraints of modern laboratory equipment (e.g., the constant sum constraint of sequencers) coupled with a pressing need for consensus involving appropriate, sensitive tools to analyze data generated by such instruments. Through analysis of simulated data spanning the variation in attributes expected across many scientific domains, we report that new computational statistical techniques have made Dirichlet-multinomial modelling (DMM) an approach that can be applied efficiently in many settings. Specifically, we report that DMM is much more sensitive than the competing approaches we examined, making DMM particularly well suited to identification of subtle shifts in relative abundance among features, such as what might be required in the study of rare, but consequential, microbes or metabolites (Lynch and Neufeld 2015, Sachdeva et al. 2019). Indeed, for some data, DMM identified many times more true positives then certain competing methods (up to approximately eight times more in extreme cases; Fig. 3). The sensitivity of DMM does, however, come at the cost of an increase in false positive rate (FPR) and a loss of precision compared to competing methods, particularly for data with skewed rank abundance profiles and large variation in sampling depth among replicates. For such challenging data, FPR increased to between 5.5–10% (Fig. S3), which we suggest may be acceptable for those practitioners tasked with analyzing challenging data and that wish to avoid missing features that truly differ among compositions. The tradeoff between sensitivity (also referred to as “recall”) and precision is well known (Buckland and Gey 1994) and we suggest that the suitability of DMM will depend on the particular needs of the practitioner. If practitioners are interested primarily in sensitivity, then our results suggest DMM is an appropriate method to choose. If, on the other hand, practitioners wish to avoid false positives, even at the expense of considerable loss of sensitivity, then other methods may be more suitable.

Aside from sensitivity, DMM provides several important ancillary benefits including the estimation of parameters that describe the data under consideration and the ability to propagate uncertainty in those estimates to downstream analyses. Propagation of uncertainty allows for a precise statement regarding the credibility of an inference and is a particular benefit of Bayesian techniques over frequentist methods. For example, to determine the extent that specific features shifted from one simulated sampling group to another, we obtained the difference between PPDs of Dirichlet 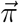 parameters from each group (Fig. 2d). A PPD is a distribution that explicitly describes the probability of certain values for a particular model parameter; thus, in the model described here, the mean of the PPD for a specific *π* parameter is a sensible point estimate for that feature’s relative abundance and the variation around that mean describes the certainty in that estimate. By subtracting PPDs for *π* parameters obtained from different sampling groups for a focal taxon, we obtain a PPD of differences, thus propagating uncertainty in relative abundance estimates through to differential relative abundance testing (Fig. 1). This provides a great deal of flexibility to practitioners, because the location of zero in this distribution of differences quantifies the probability that the two original PPDs differed—in other words, that the feature differed in relative abundance between sampling groups. We assumed that, for some feature *i* present in two sampling groups *k*, if 95% of the PPDs for *π*_*ik*_ did not overlap, then that feature differed in relative abundance between groups (see methods). If a more conservative analysis is desired, then a more strict criterion could be employed to determine if PPDs of focal features are sufficiently divergent, for instance 98% or 99%. Similarly, a less strict criterion could be used (e.g., 90%) for exploratory analyses. Moreover, because we precisely quantify uncertainty in parameter estimates derived from a single model, multiple comparison testing is unneeded for our implementation of DMM. A final benefit of quantifying uncertainty for each feature of interest is that, with some creativity, this uncertainty can be propagated to other downstream analyses, including those using derived parameters of interest such as diversity entropies (see Supplemental Material and Marion et al. 2018). The benefits provided by uncertainty propagation are primary differences between DMM as we describe it here and the competing approaches we tested that rely on some form of frequentist testing.

Another important benefit of the approach to DMM we describe is the hierarchical sharing of information among replicates from sampling groups (also see Fordyce et al. 2011). Hierarchical models make thorough use of the information present within the data, which can improve parameter estimates and propagate uncertainty, particularly when sampling effort is inconsistent among replicates and sampling groups (Coblentz et al. 2017). As described in the methods, hierarchical modelling can be used in a way analogous to frequentist, mixed effects modelling to account for non-independence among replicates through the use of a random effect (Bates et al. 2015, Björk et al. 2018). Hierarchical modelling also allows for novel inferential opportunities, given sufficient data, because parameter estimates can be extracted from any level in the model hierarchy.

### Additional considerations pertaining to Dirichlet-multinomial modelling

A downside to Bayesian modelling is its computational expense. While JAGS (Plummer 2003), BUGS (Lunn et al. 2012), Stan (Carpenter et al. 2017), and PyMC3 (Salvatier et al. 2016) have greatly simplified Bayesian model specification and implementation, Bayesian analysis can require much more computational time then frequentist methods. Users should be aware that as the number of parameters to estimate increases, so too does modelling time. For data sets of low to moderate dimensionality (i.e. less than a thousand features), the model described herein can be run on a desktop computer within several hours using any of the three PPD estimation methods (VI may take only a few seconds to run for such small data). However, for larger data sets of many thousand features, convergence when using MCMC or HMC may require a multiple days. For larger data, MCMC sampling should probably be avoided because HMC, as implemented in Stan is much faster and results in convergence for more parameters and, thus, a lower false positive rate (Fig. S6). For extremely large data, VI may be the only viable option for efficient parameter estimation. Unfortunately, we observed heightened variation in the performance of VI compared to MCMC or HMC when confronting data with a dramatic rank abundance skew—in some cases VI did as well as HMC, but in other cases it was unable to recover a high proportion of the true positives present (Fig. 3). Computational implementations of VI are a topic of current research and will undoubtedly improve over coming years (Blei et al. 2017). For most users, we suggest performing an initial analysis using both HMC and VI. If parameter estimates are largely congruent between techniques (as we generally observed), then VI could be used for subsequent analyses using similar data, thus taking advantage of VI’s efficiency.

For HMC or MCMC sampling, time to convergence can be improved through initializing the chains at sensible values for all parameters. We initialize chains for multinomial and Dirichlet parameters at their maximum likelihood values (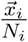, the proportion of each feature within a sampling group). Additional performance gains can be achieved by combining features that are consistently infrequent across replicates to form a composite feature. This composite feature should be included in modelling, otherwise proportion estimates will be distorted and incorrect. This approach could be particularly appropriate for analysis of high-throughput sequencing of microbiomes and transcriptomes, which often rely on data sets characterized by many features of extremely low relative abundance. Estimates of the relative abundance of very infrequent features will be imprecise, thus precluding effective comparison of relative abundances among sampling groups. Therefore, for some questions, combining these features will not lessen inferential opportunity and can greatly reduce computation time.

Some authors have suggested that the expected negative covariance of feature proportions 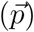 in a Dirichlet distribution is a drawback that makes this distribution undesirable (Grantham et al. 2017, Mandal et al. 2015, Weiss et al. 2016). Specifically, the elements of 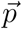 in a deviate from a Dirichlet distribution are expected to negatively covary (Mosimann 1962) according to: 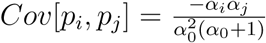, where 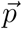 is the vector of expected proportions for features in the composition and 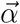 represents the Dirichlet parameter vector. Indexing of 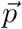 and 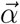 across features is achieved via *i* and *j*, and 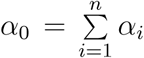, where *n* is the number of features. For even modest values of *α*_0_, the expected negative covariance between elements in 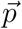 is small and diminishes rapidly with increasing *α*_0_, approaching zero in the limit of large *α*_0_. The negative covariance structure is a fundamental limitation of compositional data, as one or more features increase, other features must decline to maintain a constant sum. Thus, the Dirichlet distribution assumes a reality that mirrors the data.

There are many problems associated with the analysis of compositional data that cannot be handled by DMM alone (see Aitchison and Egozcue 2005, Gloor and Reid 2016, Quinn et al. 2017, Tsilimigras and Fodor 2016, van den Boogaart and Tolosana-Delgado 2013). The most intuitive challenge posed by compositional data is that spurious correlations among features can arise because of the data’s inherent covariance structure (Pearson 1897). For instance, shifts in the relative abundance of a dominant microbial taxon along an abiotic gradient causes shifts in the relative abundance of co-occurring taxa, even if the actual abundances of those taxa are invariant across the gradient (Fig. 1). In such a scenario, compositionality could induce associations between the relative abundances of certain taxa and the gradient that are not biologically supported. Other issues that can arise when analyzing compositional data include “sub-compositional incoherence”, which means that omission of features from the composition necessarily changes the relative abundances of the remaining features after they are renormalized to their constant sum (e.g., one for proportions; Pawlowsky-Glahn and Egozcue 2006).

The technique most relied upon to address these problems is log ratio transformation: 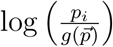, where *p*_*i*_ is the *i*^th^ feature within 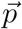, which is composed of either counts or proportions, and 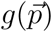 is a function. When 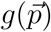 is the geometric mean of all feature abundances, this transformation is called the “centered log ratio” (CLR; Aitchison 1982). Division by the geometric mean places all replicates on the same scale and, therefore, is useful when variation in sampling effort exists among replicates. Alternatively, 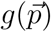 can be an indexing function and output the value of a feature, *p*_*j*_, that has a constant absolute abundance among replicates. This approach is called the “additive log ratio” (ALR) transformation (Aitchison 1982) and can be useful when an internal standard can be added to samples prior to data generation (e.g. during library preparation for next-generation sequencing; Jiang et al. 2011, Munro et al. 2014, Tourlousse et al. 2017, Tkacz et al. 2018) or when certain features are expected to be invariant among replicates (e.g. “housekeeping genes”; Eisenberg and Levanon 2013). By converting information from each feature into a ratio, both ALR and CLR avoid the sub-composition incoherence problem (Morton et al. 2019). To understand this, consider conducting the ALR transformation on replicates that each include a feature with identical absolute abundance that is used as the denominator in the transformation (it does not matter whether we consider counts or proportions for this example). The ratio between any specific feature within a replicate and the denominator will not be affected by removing other features from the composition (i.e., if the ratio is 2:1 it will remain so after omitting features from the composition and re-normalizing to maintain a constant sum). Either the CLR or ALR transformation can be applied to each MCMC sample of parameters of interest to obtain transformed PPDs for analysis (see Fernandes et al. 2014, for an example).

### Conclusions

The challenges posed by many modern molecular ecology data sets—extreme dimensionality, compositionality, and, often, stark differences in the abundance of features—have motivated the rapid development of new analytical tools and techniques. Indeed, new methods and software are published on a near monthly basis and practitioners are left to wonder which tool is best suited for the job at hand. While we do not claim DMM addresses all the challenges associated with compositional data, we do report that it is a sensitive, flexible technique that facilitates feature-specific analyses and should be added to ecologist’s toolkits (Fordyce et al. 2011). It is likely to be broadly useful and sensitive for analyses of microbiomes, other DNA barcoding, gene expression, metabolomics, and other applications in molecular ecology (Table S1). To facilitate use of DMM, we have provided an expository vignette in the Electronic Supplemental Material that provides an example of how to perform DMM using both Stan and JAGS in the R environment.

The success of DMM for relative abundance estimation, as demonstrated herein, coupled with the aforementioned benefits of hierarchical Bayesian modelling, justifies extension of the DMM to determine the effects of covariates on relative abundances and to characterize mixtures of compositions (sensu Chen and Li 2013, Holmes et al. 2012, Knights et al. 2011, Shafiei et al. 2015, Tang and Chen 2018). We look forward to continued method development along these lines.

## Acknowledgments

We wish to thank Claire Duvallet and the co-authors of Duvallet et al. (2019) for making their well-curated data available to the public. Specific thanks to Dr. Duvallet for helpful interpretation regarding our reanalysis of her and her co-author’s data. Additional thanks to helpful comments from James Fordyce and two anonymous reviewers. This research was supported by the Microbial Ecology Collaborative at the University of Wyoming with funding from NSF award #EPS-1655726. Computing was performed in the Teton Computing Environment at the Advanced Research Computing Center, University of Wyoming, Laramie (https://doi.org/10.15786/M2FY47).

## Data Accessibility

All scripts and processed data used for this manuscript are available at https://github.com/JHarrisonEcoEvo/DMM Harrison et al. 2019 and a snapshot corresponding to the status at publication at Zenodo (10.5281/zenodo.3558682). Data from Duvallet et al. (2019) can be downloaded from (DOI: 10.5281/zenodo.2678108).

### Author contributions

All authors contributed to model development and manuscript preparation.

## Supplementary Material

### Supplemental Methods

#### Determination of feature abundance class

Deviates used to simulate data were divided into abundance classes to ensure that features of each abundance class were made to differ between sampling groups. All features were assigned to the intermediate abundance class when Dirichlet 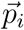 parameters were assigned a constant value. When the Pareto distribution with shape parameter of four was used, deviates greater than or equal to five were assigned to the abundant class, deviates in the intermediate class were between two and five, and deviates within the rare class were less than two. When the Pareto distribution with shape parameter of 0.7 was used, deviates greater than 1000 were assigned to the abundant class, deviates between 1000 and 100 to the intermediate class, and deviates less than 100 to the rare class. These thresholds were chosen through visual examination of frequency distributions of deviates from distributions. Recall that the location parameter (minimum value) of the Pareto distributions was set to one.

#### Implementation of competing software

For analyses conducted using ALDEx2 v1.14.1 (Fernandes et al. 2014) we drew 1000 MCMC samples, which were transformed using the CLR (denom = all). Welch’s t tests and general linear models were used to determine differential relative abundance. We used a *p* value threshold of 0.05 to determine significance after applying a Benjamini-Hochberg FDR correction.

Options used for ANCOM v2.0 (Mandal et al. 2015) included a significance value of 0.05, a “less stringent” multiple comparison correction (multcorr = 2), “prev.cut” was set to 0.99 (meaning features that were not observed in 99% or more of samples were omitted), and “repeated” was set to “False”. During analysis we uncovered an apparent error in the ANCOM v1.1-3 software. On occasion, ANCOM would suggest that all features within a data set differed significantly between groups. This error was not stable, though errors did seem to only occur when data were generated using the Pareto distribution. Upon further research, we found others have reported this error on the QIIME forums (Caporaso et al. 2010). To work around this problem, during the very rare cases when ANCOM reported ≥ 90% of features were significant, we identified significantly differing features as those with non-zero *w* parameters (the test statistic used by ANCOM). This resulted in very similar results among replicate analyses of data simulated using the same parameters, but that did not trigger the aforementioned error. Subsequently, we shifted analyses to rely on ANCOM v2.0, but we left this solution in place in the event that v2.0 suffered from the same error we observed in v1.1-3.

We used default options for DESeq2 v1.18.1 (Love et al. 2014). The “nbinomWaldTest” function was used to determine differential relative abundance. Significant differences were defined at *p* ≤ 0.05 after a multiple comparison correction that was calculated by DESeq2.

Default options were used for edgeR v3.20.9 (Robinson et al. 2010). After dispersion estimates were calculated using the “estimateDisp” function, the “glmQLFit” and “glmQLFTest” functions were used to determine differential relative abundance. Features differing in relative abundance were determined using the “topTags” function with a Benjamini-Hochberg FDR correction and *p* ≤ 0.05 threshold.

Default options were used for mvabund v4.0.1 (Wang et al. 2019). Simulated data were converted into an mvabund object using the “mvabund” function. The “manyglm” function was used to implement a non-hierarchical linear model where each taxon was the response and treatment group was a categorical predictor variable. A negative binomial distribution was used for the GLM and the parameter “cor.type” set to “shrink” to account for correlation among response variables. Results from the GLM were determined using the “anova” function with a Wald test and a multiple comparison correction using a step-down resampling algorithm described in Wang et al. (2012) and Westfall and Young (1993).

R was used to implement all software.

#### Examples of possible derived parameters

Derived parameters can be calculated from the output of Dirichlet-multinomial modelling while preserving the uncertainty quantified by the model. For example, many microbial and community ecologists wish to compare diversity indices among sampling groups (Jost 2007, Marion et al. 2015). Diversity indices can be calculated for each sample of the Dirichlet’s 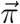 parameter vector, thus generating a PPD of diversity statistics for each sampling group. PPDs of diversity could then be compared between sampling groups through subtraction (see Harrison et al. 2019 for an example). This conceptual approach was first described by Marion et al. (2018), though the model in that study relied upon a multivariate normal prior with softmax transformation, instead of the Dirichlet prior we use here.

---

**Table S1:** Scientific domains that often rely on compositional data with brief examples of such data with associated methods (if applicable). This table is not meant to be exhaustive, but to draw awareness to the ubiquity of compositional data across the sciences. Asterisks denote data often consisting of proportions rather than counts.

| Domain | Example data |
| --- | --- |
| Molecular biology | gene expression characterization (RNA sequencing)<br>chromatin immunoprecipitation sequencing (ChIP-Seq)<br>flow cytometry* |
| Analytical chemistry | chemical concentrations as determined through various methods<br>(e.g., mass spectrometry)*<br>elemental composition* |
| Microbiology/Microbial ecology | colony/cell counts*<br>amplicon sequencing |
| Ecology | species counts*<br>DNA barcode based community characterization<br>foraging preference assays*<br>allele frequencies*<br>haplotype counts<br>chemical concentrations*<br>elemental composition* |
| Paleolimnology | pollen counts*<br>foraminifera counts* |
| Geology | mineral composition*<br>sediment composition* |
| Psychology | behavioral characterization* |
| Economics | budget composition<br>portfolio composition |

**Table S2:**
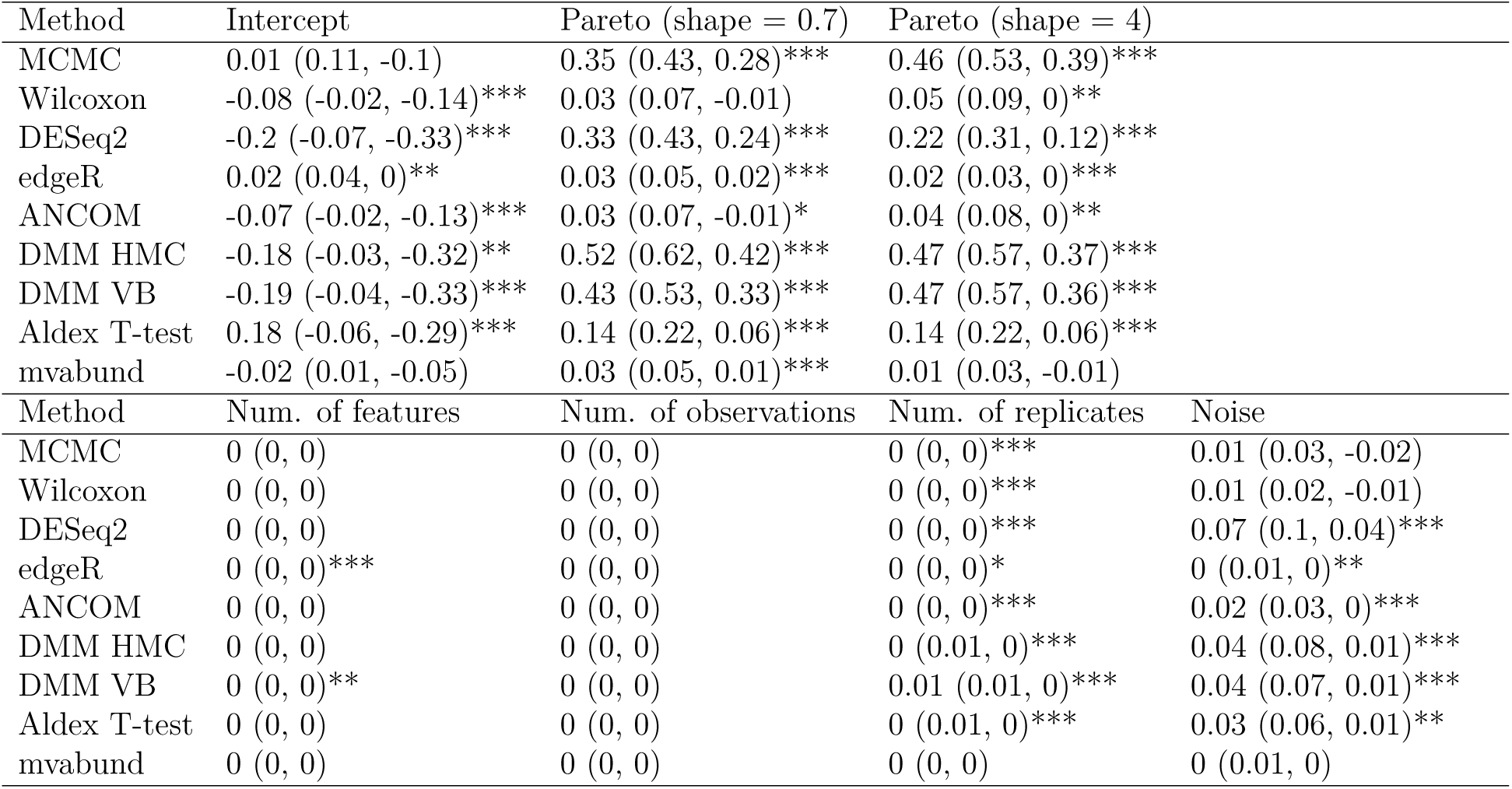
Influence of data attributes on true positive recovery. Results shown are beta coefficients and 95% confidence intervals from a multiple regression analysis. The response variable was the proportion of true positives recovered. The rank abundance profile of the data was a categorical variable, with point mass as the reference condition. The table is split into two panels of results to aid visualization. *** denotes p < 0.01, ** p < 0.05, * p < 0.1.

**Table S3:**
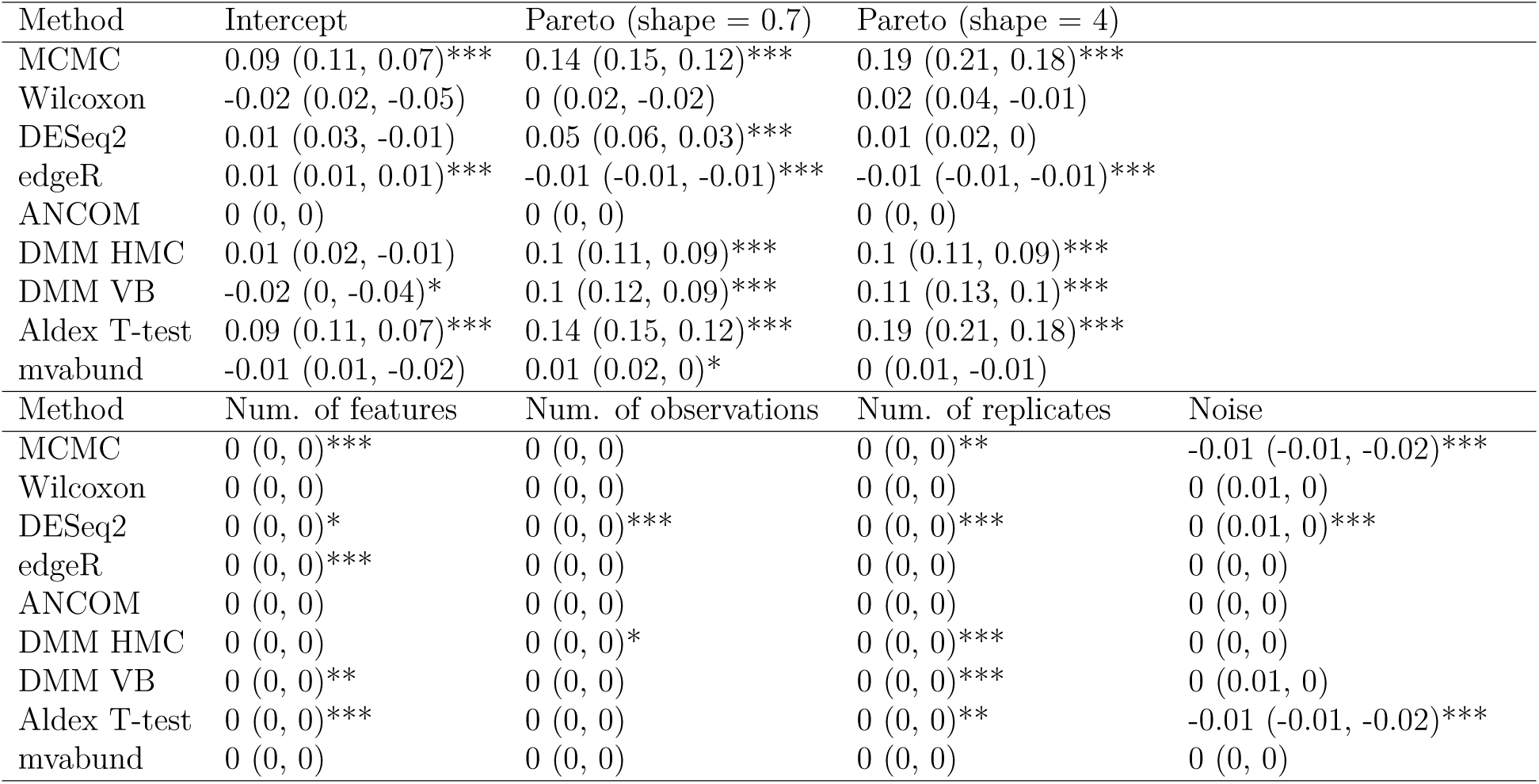
Influence of data attributes on false positive rate. Results shown are beta coefficients and 95% confidence intervals from a multiple regression analysis. The response variable was the average false positive rate among the three replicates for a unique combination of data attributes. The rank abundance profile of the data was a categorical variable, with point mass as the reference condition. The table is split into two panels of results to aid visualization. *** denotes p < 0.01, ** p < 0.05, * p < 0.1.

**Figure S1:**
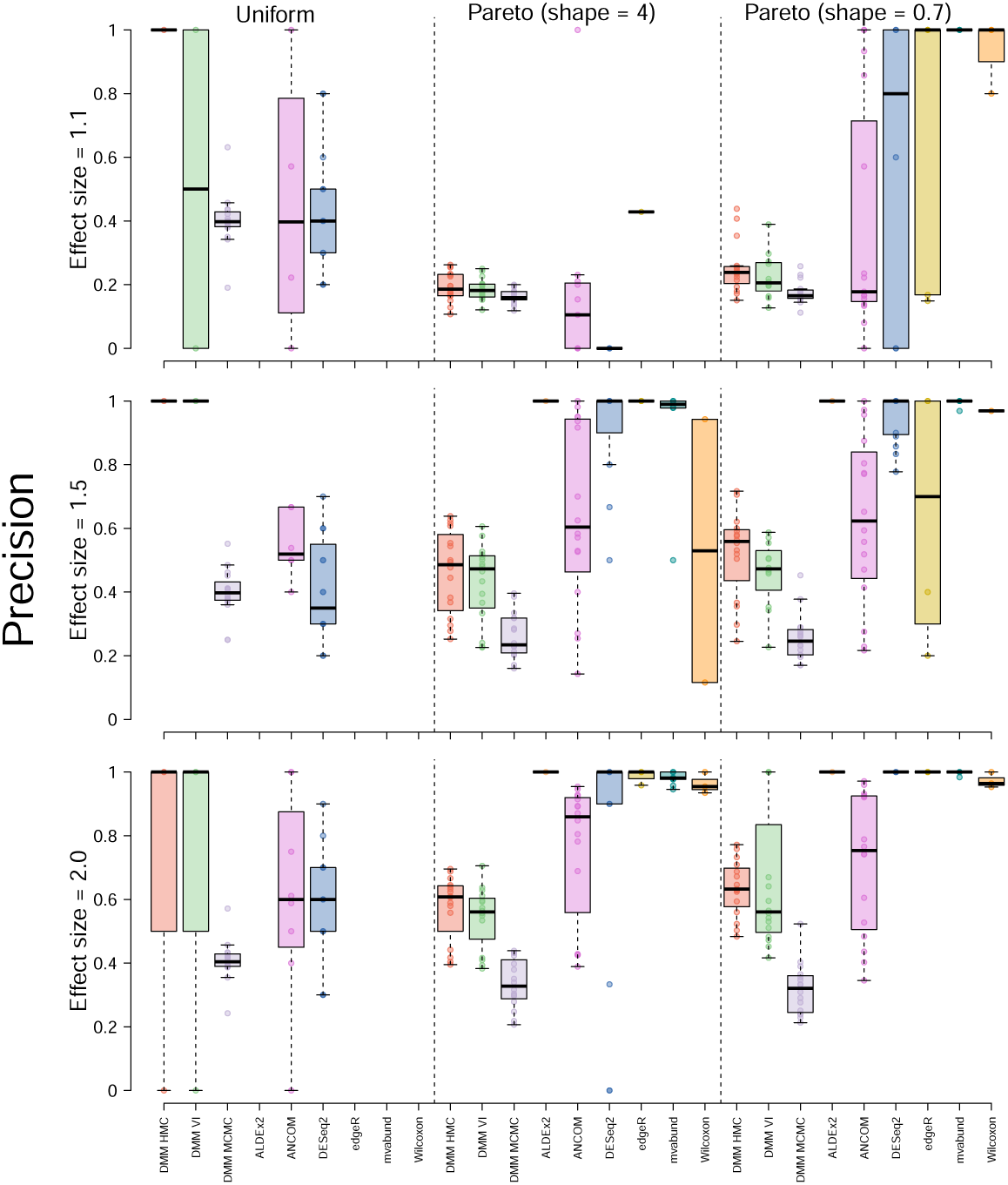
Precision of all methods competed. Precision was calculated as 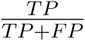, where TP stands for true positives and FP for false positives. This is a measure of how many of the positives suggested by the model are actually true. Precision is shown on the y axis and model type on the x axis. Rows describe model performance when identifying features that differ between treatment groups by a certain effect size (e.g., 1.1 in the top row). Each row is broken up into three columns that are separated by a dotted line. These columns show results for data with differing rank abundance profiles.

**Figure S2:**
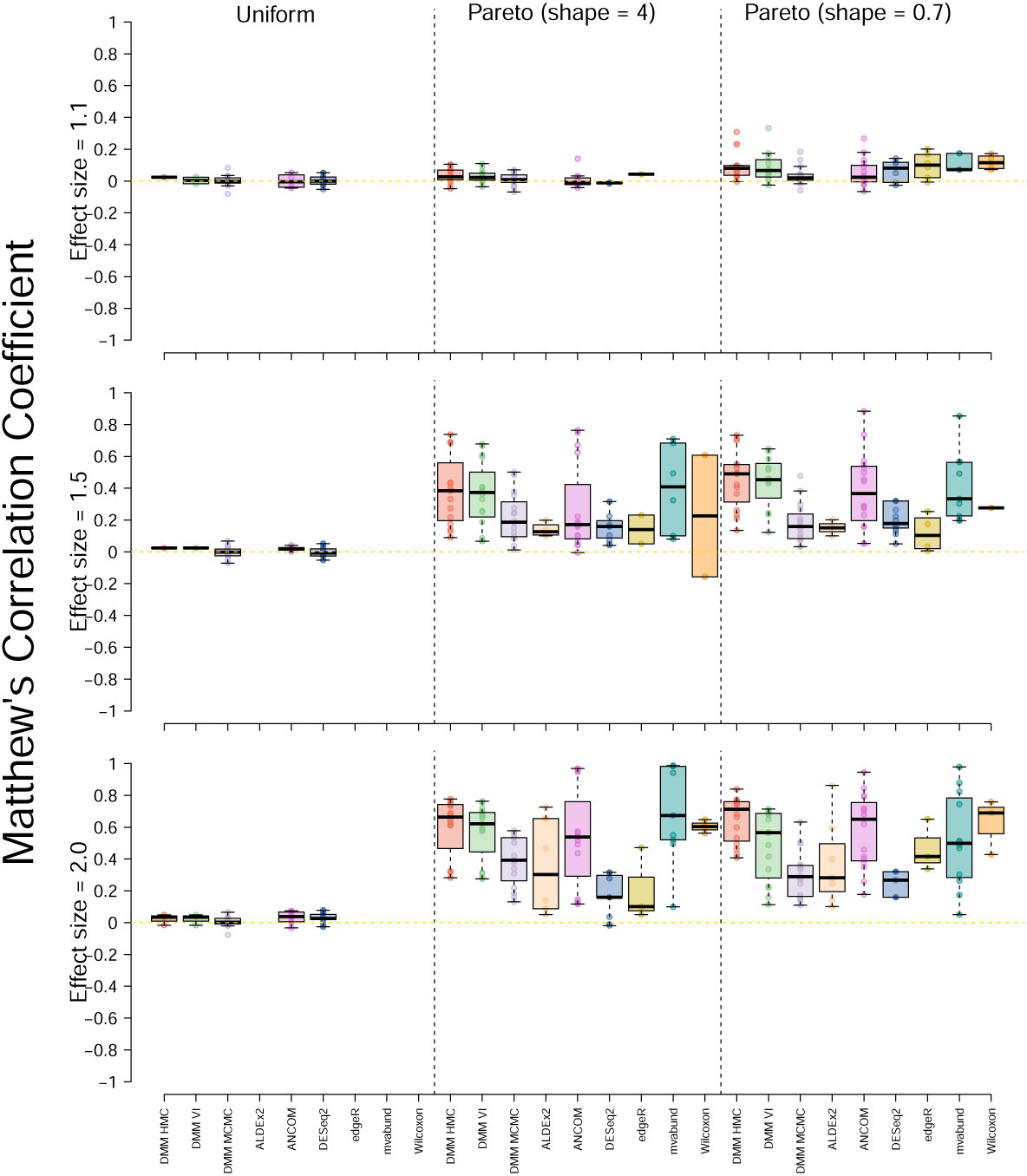
Matthew’s Correlation Coefficient (MCC) of all methods competed. MCC is the correlation between actual and predicted classifications and varies from one (perfect classification) to negative one (completely incorrect classification). An MCC of zero denotes the classifier performed no better than random guessing. MCC was calculated as 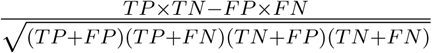, where TP stands for true positives, FP for false positives, TN for true negatives, and FN for false negatives. MCC is shown on the y axis and model type on the x axis. Rows describe model performance when identifying features that differ between treatment groups by a certain effect size (e.g., 1.1 in the top row). Each row is broken up into three columns that are separated by a dotted line. These columns show results for data with differing rank abundance profiles.

**Figure S3:**
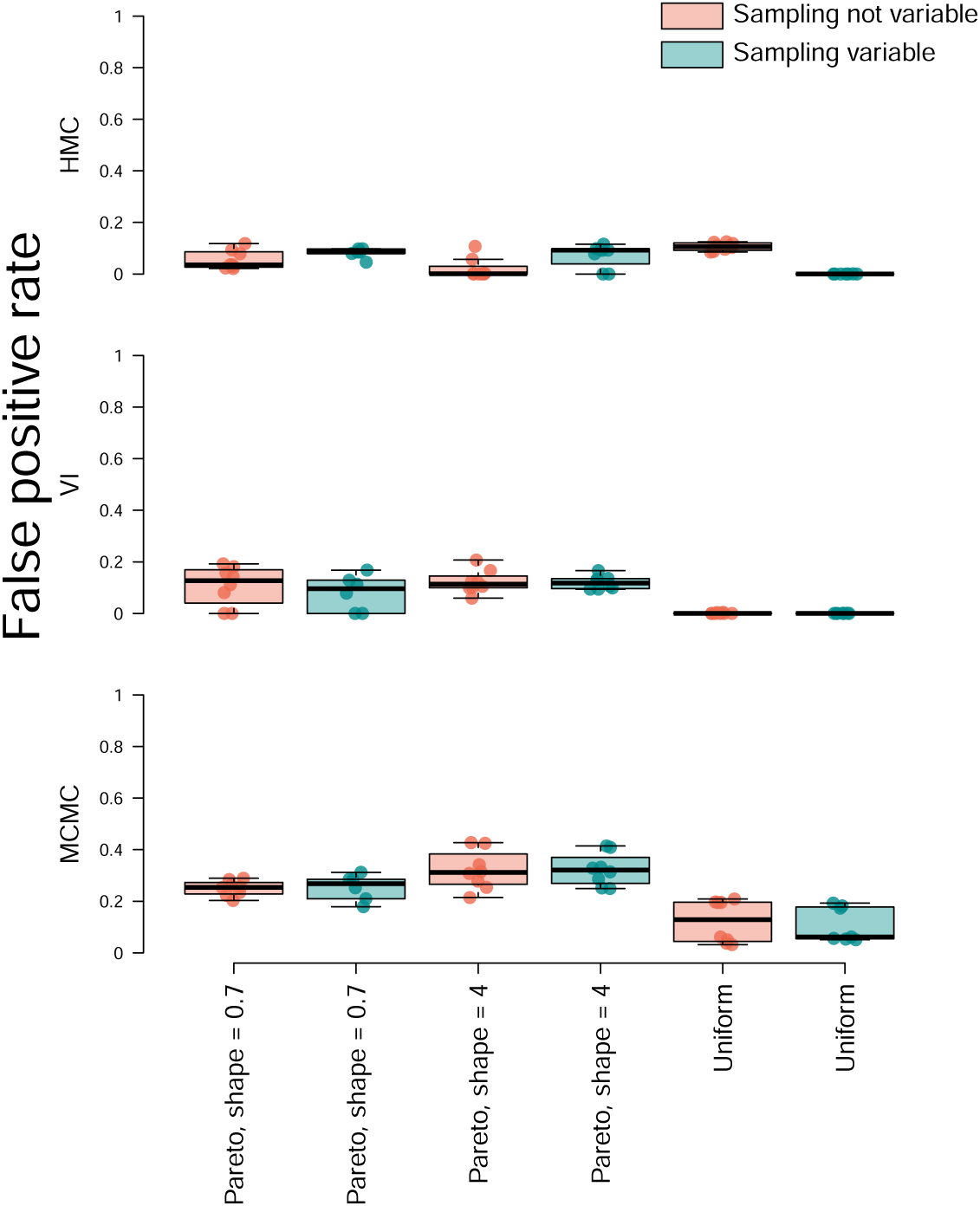
False positive rate of DMM as implemented via HMC (top row), VI (middle row), and MCMC (bottom row) when confronted with data where no features were expected to differ between sampling groups. The distribution used to simulate the data is shown on the x axis (see main text). Each point is the result from a different simulated data set; data were simulated using a variety of parameters encompassing a representative subset of the attributes considered in our main simulation (number of features ∈ {500, 2000}, 10000 samples per replicate, number of replicates ∈ {10, 50}, intensity parameter ∈ {0.5, 3}). FPR was calculated as the proportion of features that were incorrectly estimated to vary between treatment groups (false positives divided by the sum of false positives and true negatives). For a subset of the simulated data, sampling depth was made to vary by as much as two orders of magnitude between replicates (i.e., a sum of 1000 in one replicate and a sum of 100,000 in another replicate within the same sampling group). The results from analysis of data with variation in sampling depth are shown in blue.

**Figure S4:**
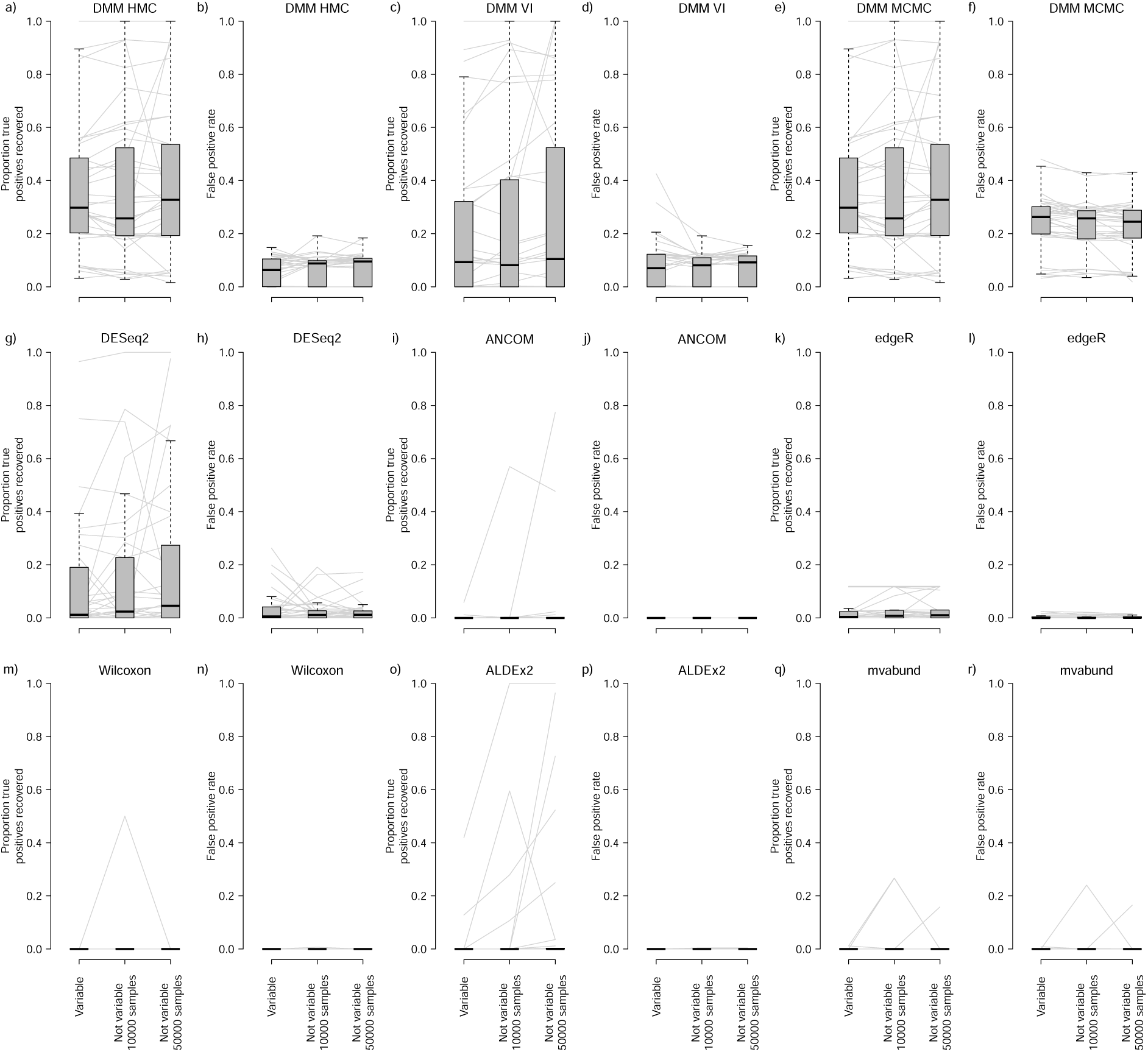
Effect of variation in sampling effort on model performance. Replicates were made to differ in sampling effort by up to two orders of magnitude (see main text) and the results from analysis of these data are shown in the first box of each panel (“variable”). Subsequent boxes show results from data where among replicate sampling effort was fixed at either 10,000 or 50,000 samples. Gray lines connect results from data sets that were simulated using identical parameters, except for sampling effort. Data were simulated using a representative subset of the parameters used for our main simulation experiment. Boxplots follow the format described in Fig. 3.

**Figure S5:**
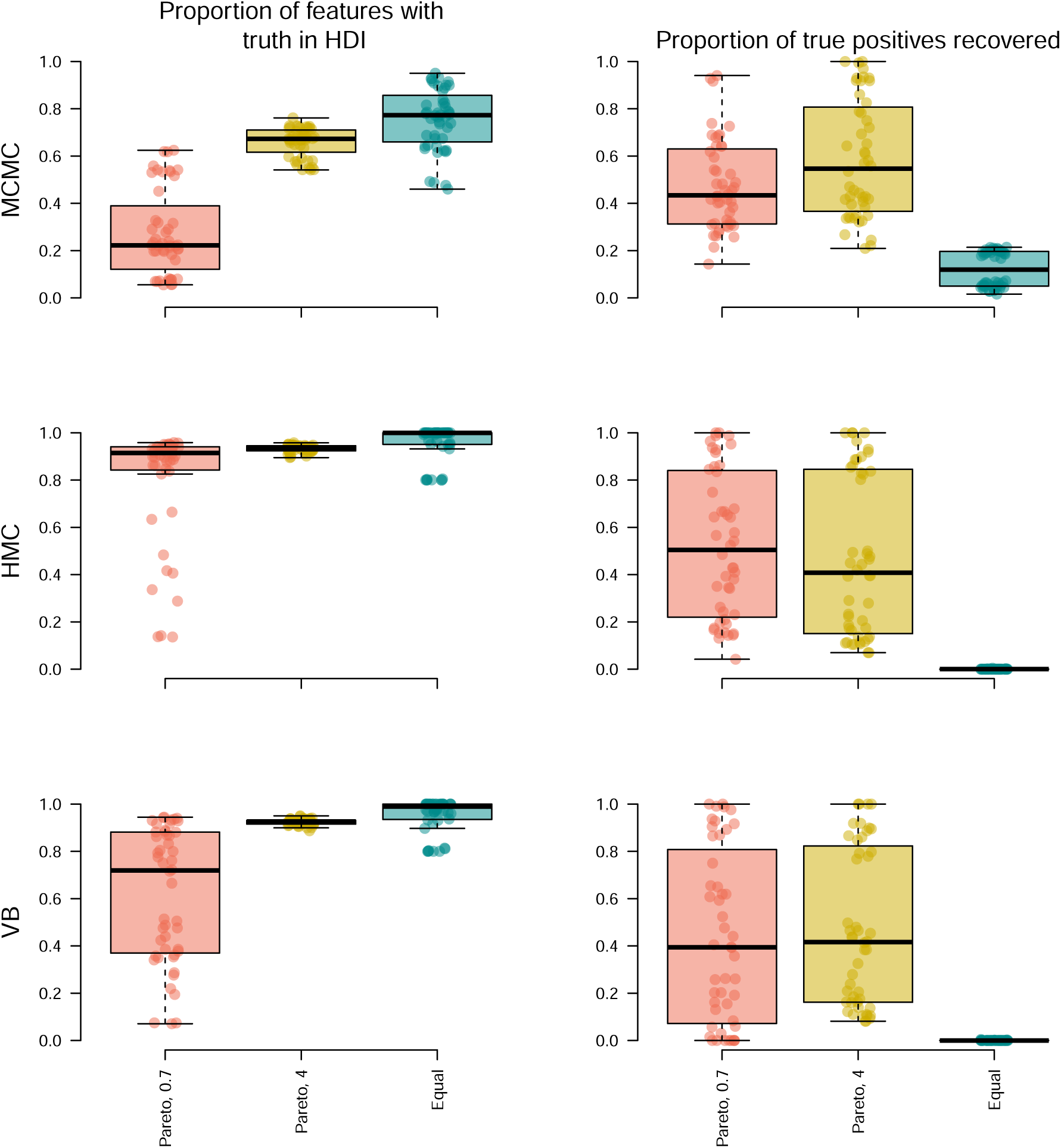
Proportion of times that high density intervals of posterior probability distributions for Dirichlet parameters (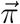 parameters) included the true, simulated parameters (left column). In the right column, the proportion of true positives recovered is shown. Boxplots are shown for results from data simulated to have differing rank abundance curves (“Pareto, 0.7” was most skewed, “Equal” was least skewed; see main text). Boxplots follow the format described in Fig. 3.

**Figure S6:**
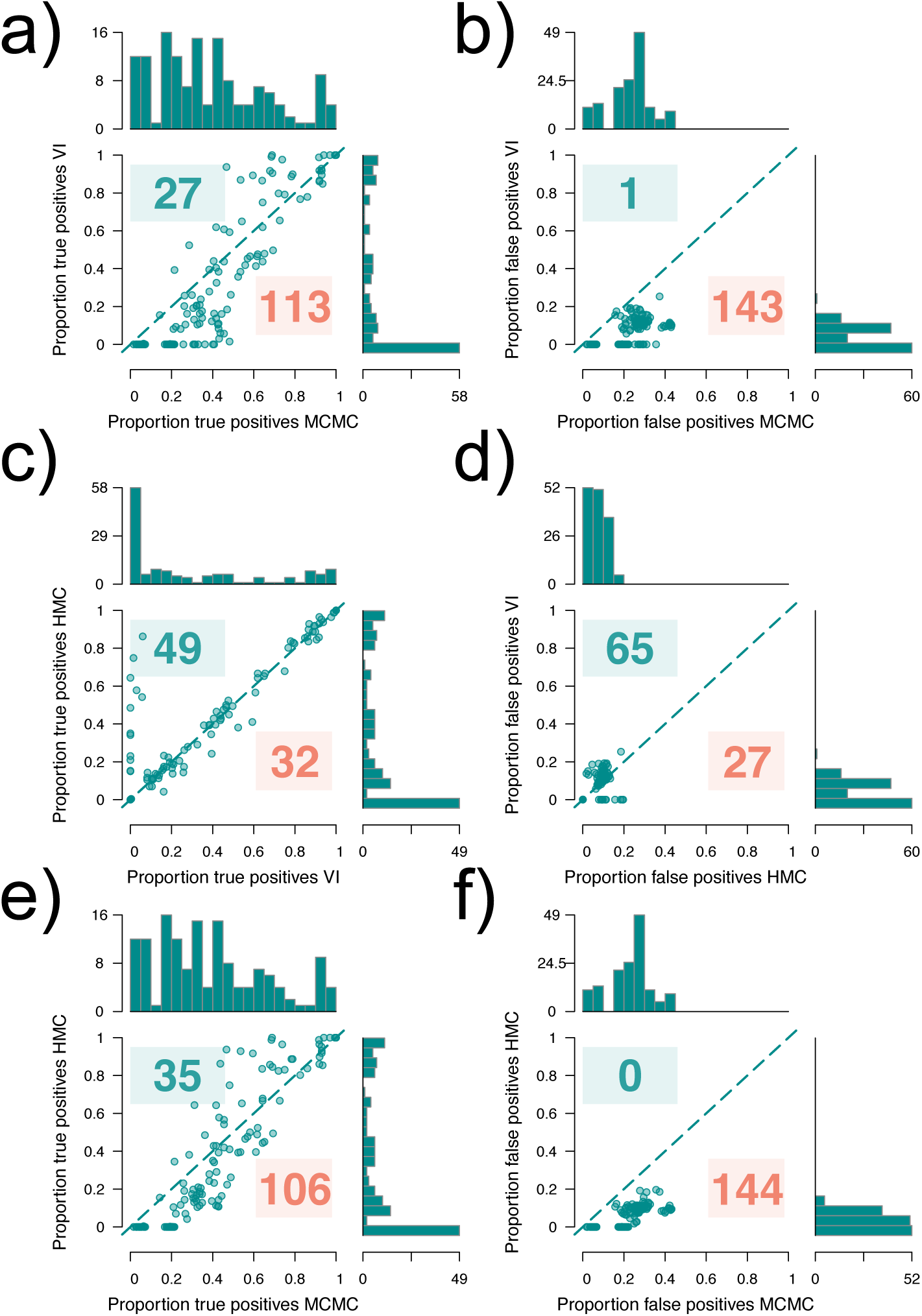
Comparison of true positive rate among the three parameter estimation methods tested (panels a,c,e). Comparison of false discovery rate between the same three methods (b,d,f). This figure corresponds in format to Fig. 4. Marginal histograms are provided to aid visualization. For details of model implementation and parameter estimation methods, see the main text (VI: variational inference; HMC: Hamiltonian Monte Carlo; MCMC: JAGS model implementation).

**Figure S7:**
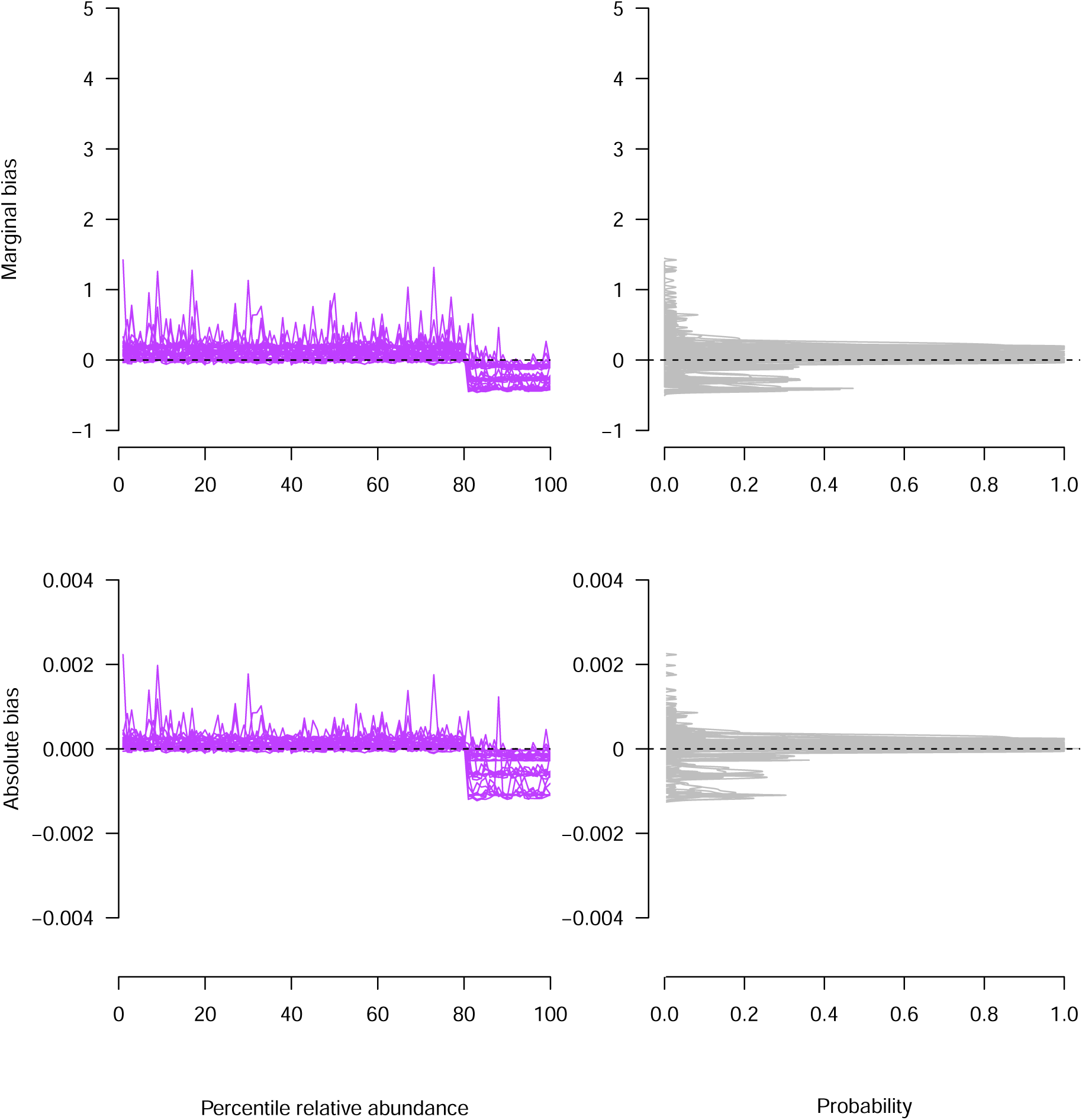
Bias of DMM as implemented via HMC as a function of feature relative abundance for data sets simulated using a uniform rank abundance distribution. Percentiles of relative abundances for each dataset were calculated and are shown on the x axis of bias plots (left column), thus normalizing for the differences in numbers of features among datasets. Bias (defined as the difference between predicted values and the truth) in *π* parameters is shown on the y axis. Marginal bias was calculated as absolute bias divided by the relative abundance of the focal parameter. Plots in the right column show probability densities for the different bias values shown in the left column. Each line denotes results from a simulated dataset. Results shown here are from a representative subset of the datasets simulated as part of our main experiment including 99 datasets with variable counts among replicates (see Methods) and 45 datasets with invariant counts among replicates. All values that we considered as part of our main experiment for number of features, number of observations, number of replicates, precision (*θ*), and effect sizes were included in the subset of the datasets analyzed here.

**Figure S8:**
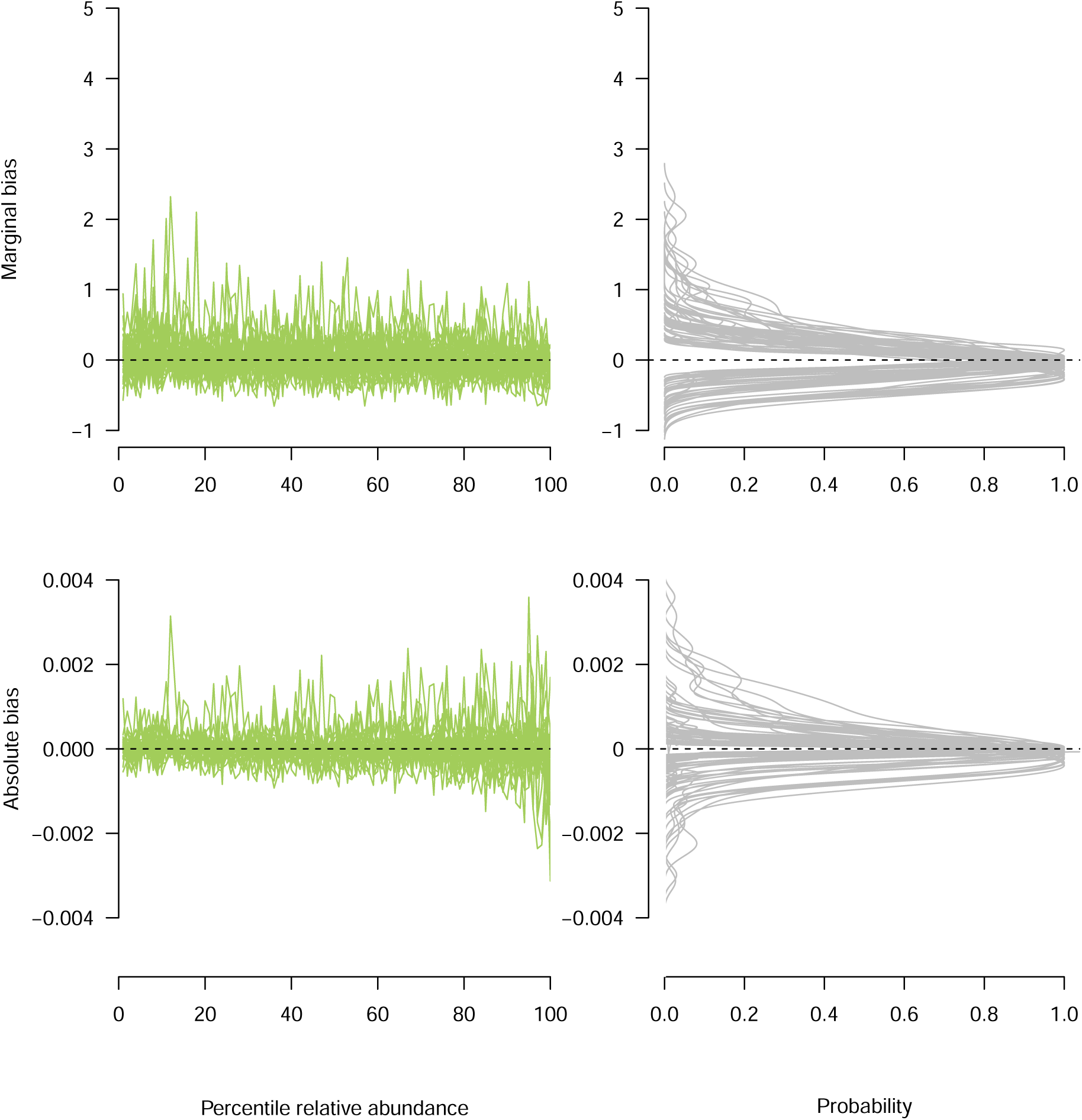
Bias of DMM as implemented via HMC as a function of feature relative abundance for data sets simulated using a moderately skewed rank abundance distribution (Pareto = 4). Percentiles of relative abundances for each dataset were calculated and are shown on the x axis of bias plots (left column), thus normalizing for the differences in numbers of features among datasets. Bias (defined as the difference between predicted values and the truth) in *π* parameters is shown on the y axis. Marginal bias was calculated as absolute bias divided by the relative abundance of the focal parameter. Plots in the right column show probability densities for the different bias values shown in the left column. Each line denotes results from a simulated data set. Results shown here are from a representative subset of the datasets simulated as part of our main experiment including 99 datasets with variable counts among replicates (see Methods) and 45 datasets with invariant counts among replicates. All values that we considered as part of our main experiment for number of features, number of observations, number of replicates, precision (*θ*), and effect sizes were included in the subset of the datasets analyzed here.

**Figure S9:**
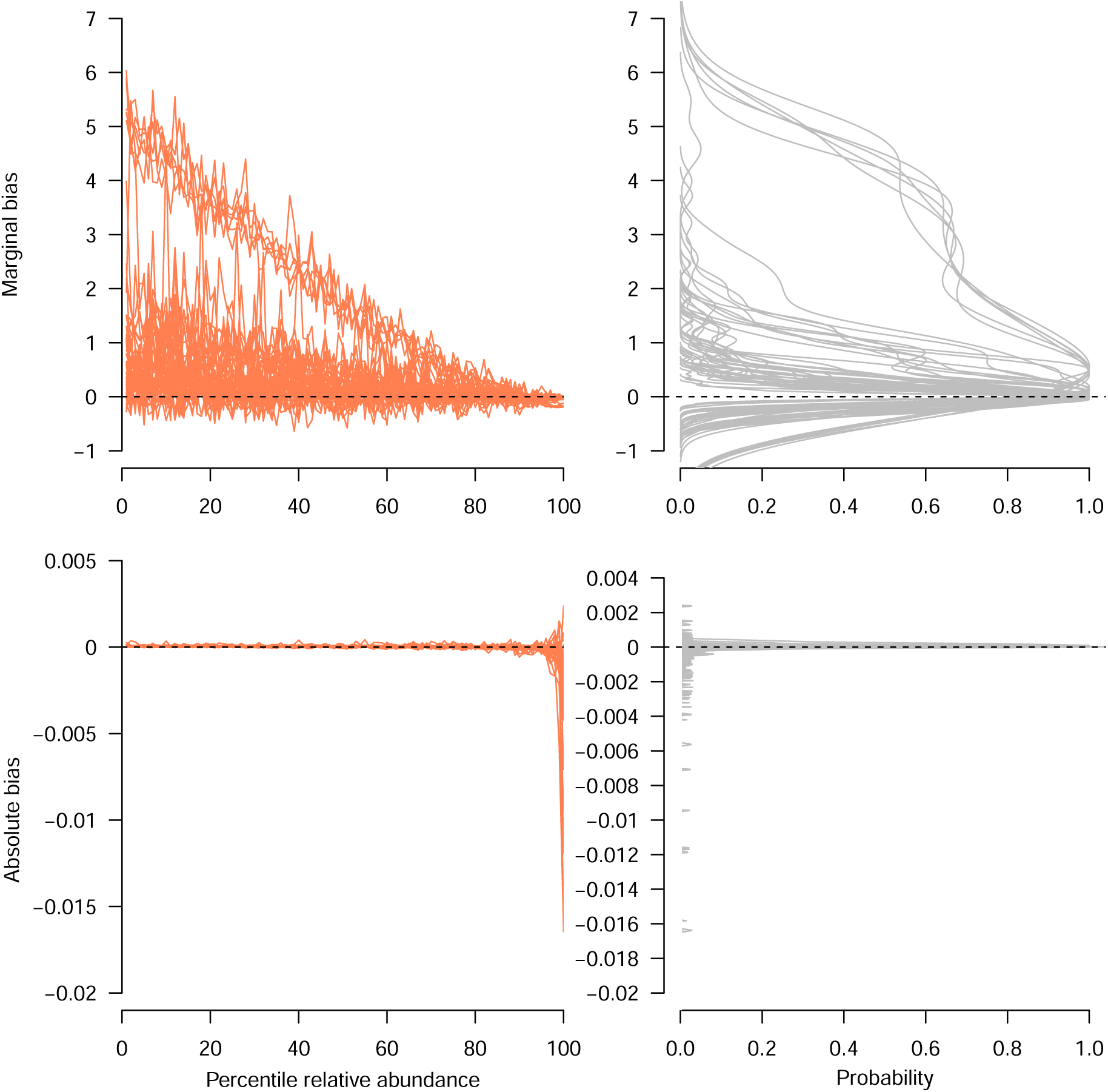
Bias of DMM as implemented via HMC as a function of feature relative abundance for data sets simulated using a highly skewed rank abundance distribution (Pareto = 0.7). Percentiles of relative abundances for each dataset were calculated and are shown on the x axis of bias plots (left column), thus normalizing for the differences in numbers of features among datasets. Bias (defined as the difference between predicted values and the truth) in *π* parameters is shown on the y axis. Marginal bias was calculated as absolute bias divided by the relative abundance of the focal parameter. Plots in the right column show probability densities for the different bias values shown in the left column. Each line denotes results from a simulated data set. Results shown here are from a representative subset of the datasets simulated as part of our main experiment including 99 datasets with variable counts among replicates (see Methods) and 45 datasets with invariant counts among replicates. All values that we considered as part of our main experiment for number of features, number of observations, number of replicates, precision (*θ*), and effect sizes were included in the subset of the datasets analyzed here.

**Figure S10:**
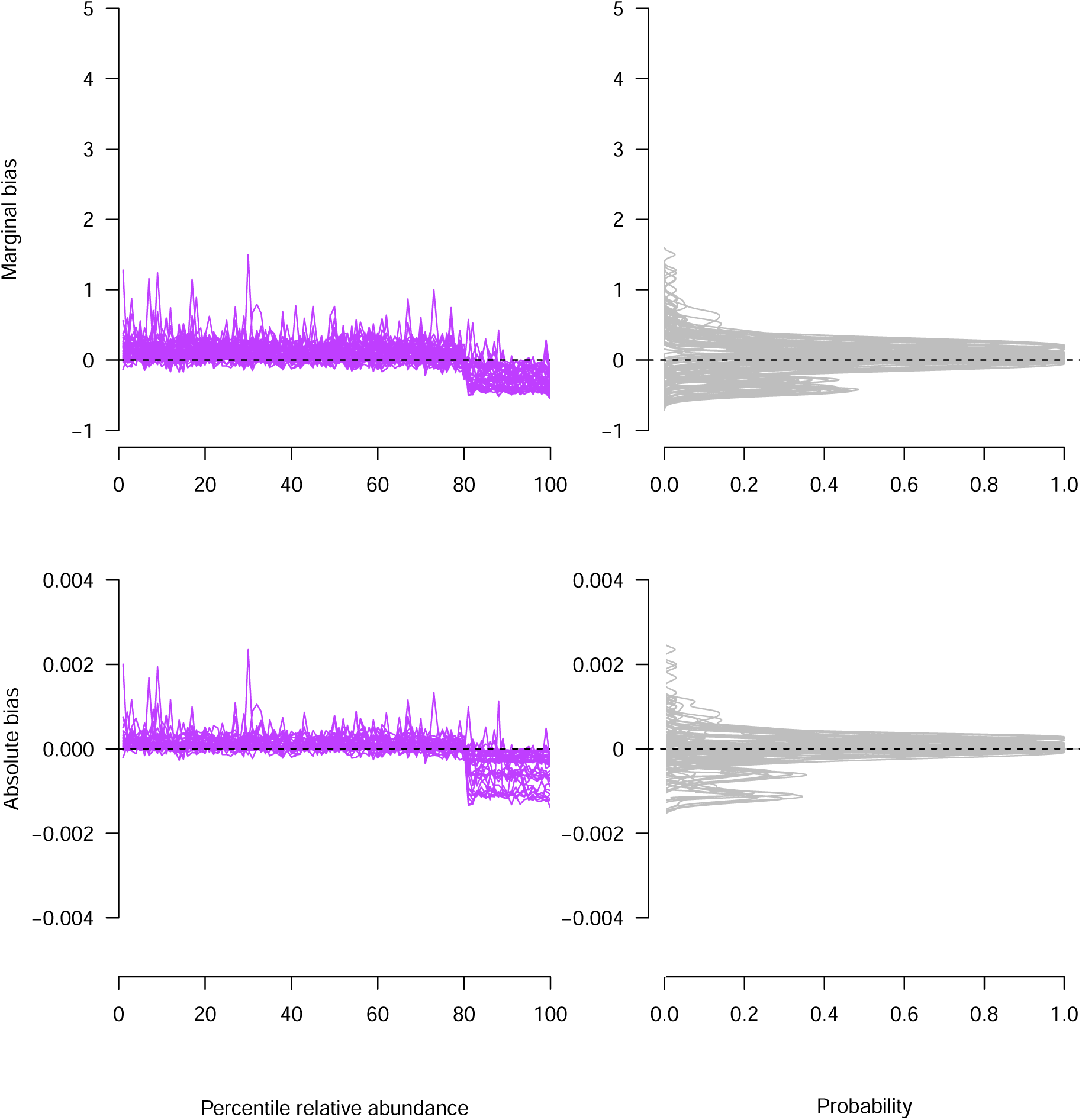
Bias of DMM as implemented via VI as a function of feature relative abundance for data sets simulated using a uniform rank abundance distribution. Percentiles of relative abundances for each dataset were calculated and are shown on the x axis of bias plots (left column), thus normalizing for the differences in numbers of features among datasets. Bias (defined as the difference between predicted values and the truth) in *π* parameters is shown on the y axis. Marginal bias was calculated as absolute bias divided by the relative abundance of the focal parameter. Plots in the right column show probability densities for the different bias values shown in the left column. Each line denotes results from a simulated data set. Results shown here are from a representative subset of the datasets simulated as part of our main experiment including 99 datasets with variable counts among replicates (see Methods) and 45 datasets with invariant counts among replicates. All values that we considered as part of our main experiment for number of features, number of observations, number of replicates, precision (*θ*), and effect sizes were included in the subset of the datasets analyzed here.

**Figure S11:**
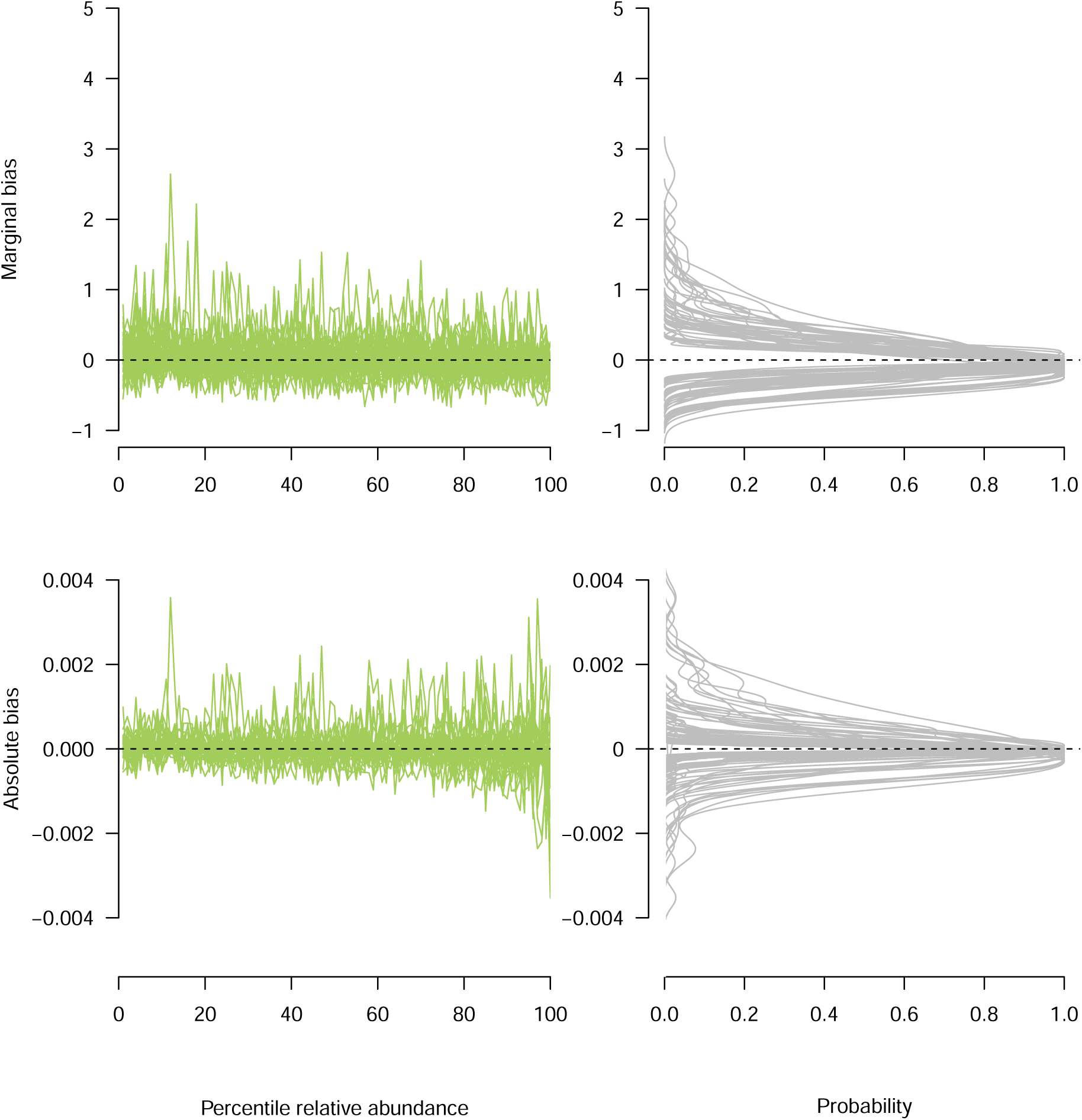
Bias of DMM as implemented via VI as a function of feature relative abundance for data sets simulated using a moderately skewed rank abundance distribution (Pareto = 4). Percentiles of relative abundances for each dataset were calculated and are shown on the x axis of bias plots (left column), thus normalizing for the differences in numbers of features among datasets. Bias (defined as the difference between predicted values and the truth) in *π* parameters is shown on the y axis. Marginal bias was calculated as absolute bias divided by the relative abundance of the focal parameter. Plots in the right column show probability densities for the different bias values shown in the left column. Each line denotes results from a simulated data set. Results shown here are from a representative subset of the datasets simulated as part of our main experiment including 99 datasets with variable counts among replicates (see Methods) and 45 datasets with invariant counts among replicates. All values that we considered as part of our main experiment for number of features, number of observations, number of replicates, precision (*θ*), and effect sizes were included in the subset of the datasets analyzed here.

**Figure S12:**
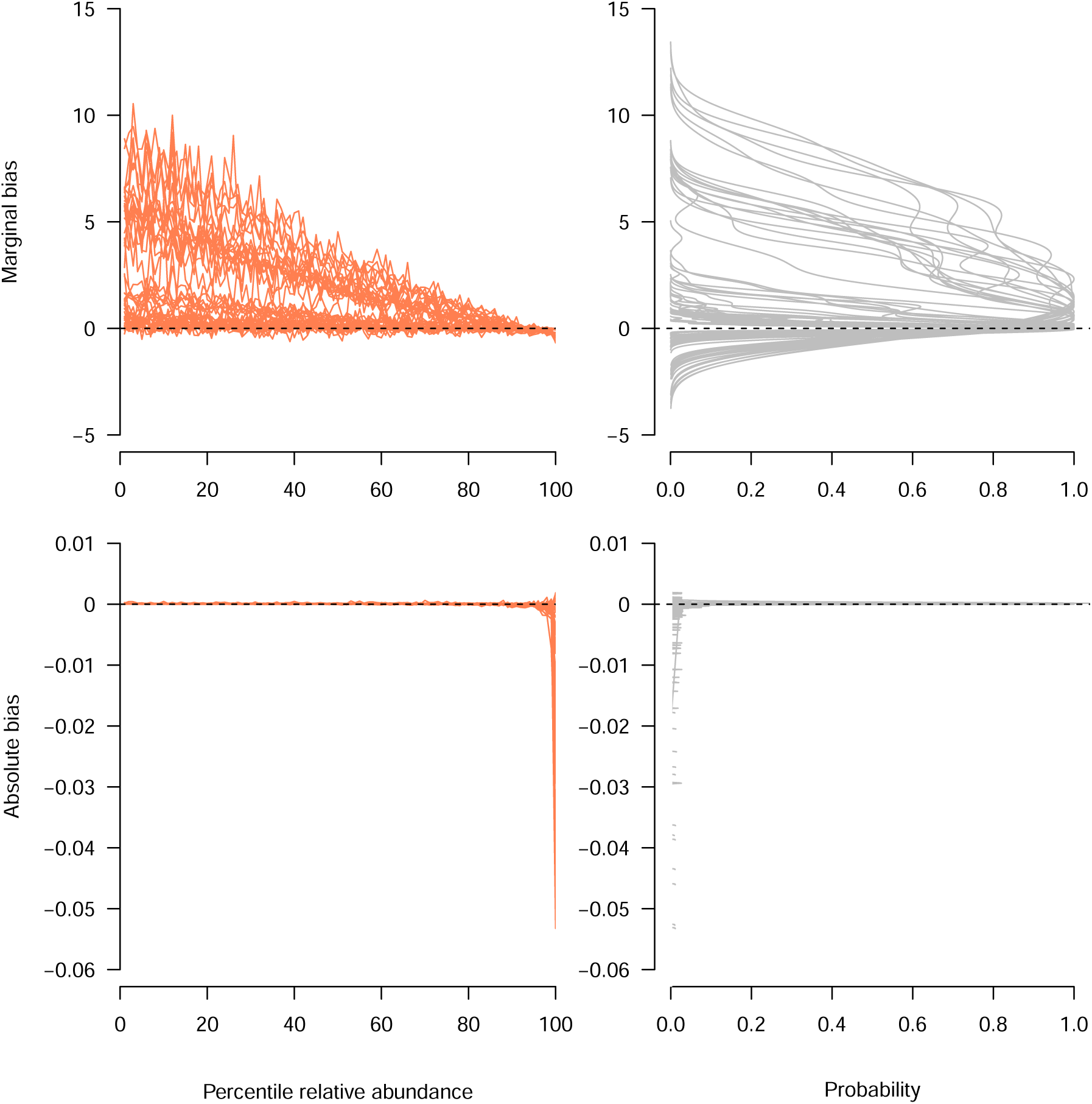
Bias of DMM as implemented via VI as a function of feature relative abundance for data sets simulated using a highly skewed rank abundance distribution (Pareto = 0.7). Percentiles of relative abundances for each dataset were calculated and are shown on the x axis of bias plots (left column), thus normalizing for the differences in numbers of features among datasets. Bias (defined as the difference between predicted values and the truth) in *π* parameters is shown on the y axis. Marginal bias was calculated as absolute bias divided by the relative abundance of the focal parameter. Plots in the right column show probability densities for the different bias values shown in the left column. Each line denotes results from a simulated data set. Results shown here are from a representative subset of the datasets simulated as part of our main experiment including 99 datasets with variable counts among replicates (see Methods) and 45 datasets with invariant counts among replicates. All values that we considered as part of our main experiment for number of features, number of observations, number of replicates, precision (*θ*), and effect sizes were included in the subset of the datasets analyzed here.

**Figure S13:**
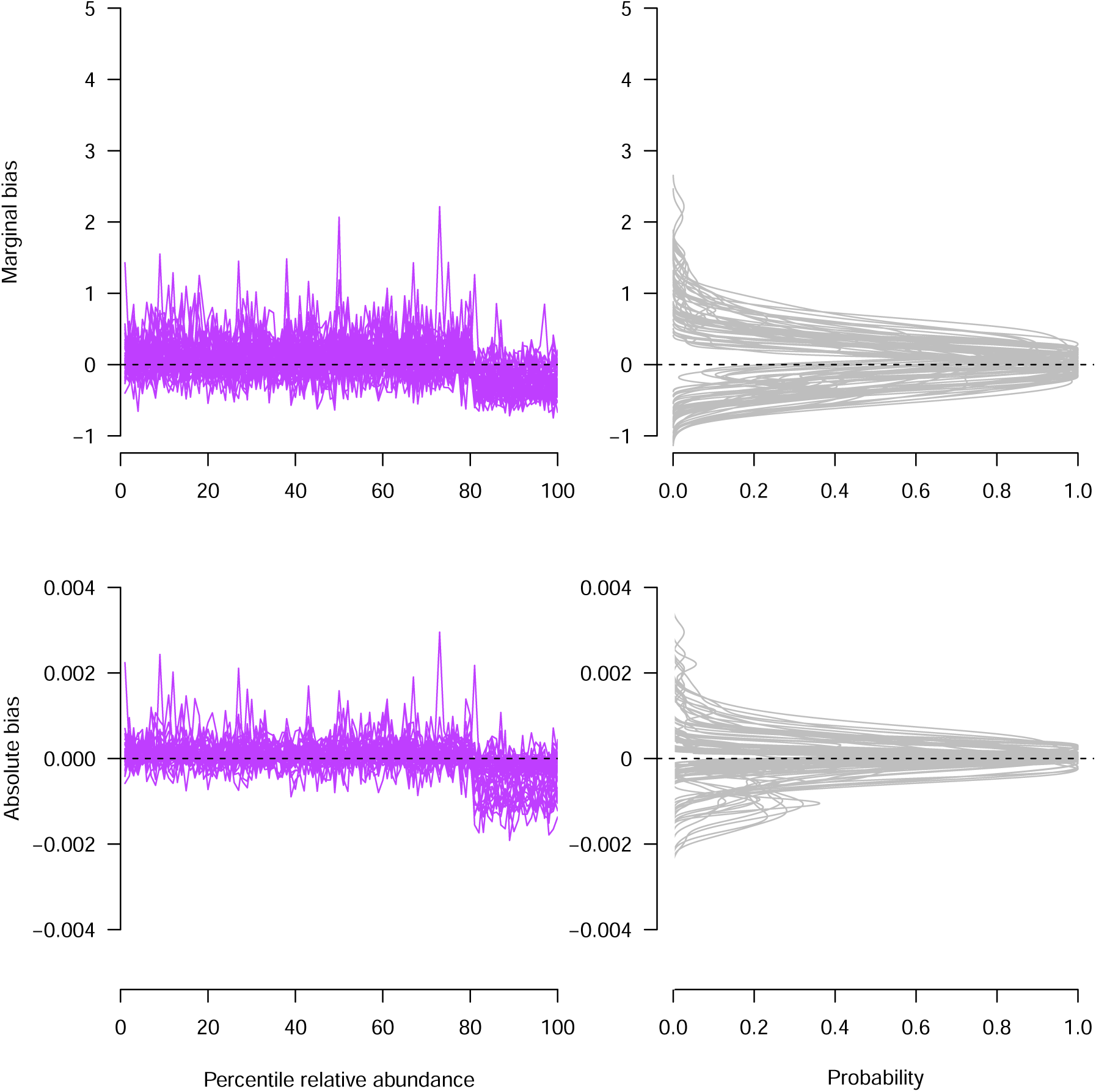
Bias of DMM as implemented via MCMC (via JAGS) as a function of feature relative abundance for data sets simulated using a uniform rank abundance distribution. Percentiles of relative abundances for each dataset were calculated and are shown on the x axis of bias plots (left column), thus normalizing for the differences in numbers of features among datasets. Bias (defined as the difference between predicted values and the truth) in *π* parameters is shown on the y axis. Marginal bias was calculated as absolute bias divided by the relative abundance of the focal parameter. Plots in the right column show probability densities for the different bias values shown in the left column. Each line denotes results from a simulated data set. Results shown here are from a representative subset of the datasets simulated as part of our main experiment including 99 datasets with variable counts among replicates (see Methods) and 45 datasets with invariant counts among replicates. All values that we considered as part of our main experiment for number of features, number of observations, number of replicates, precision (*θ*), and effect sizes were included in the subset of the datasets analyzed here.

**Figure S14:**
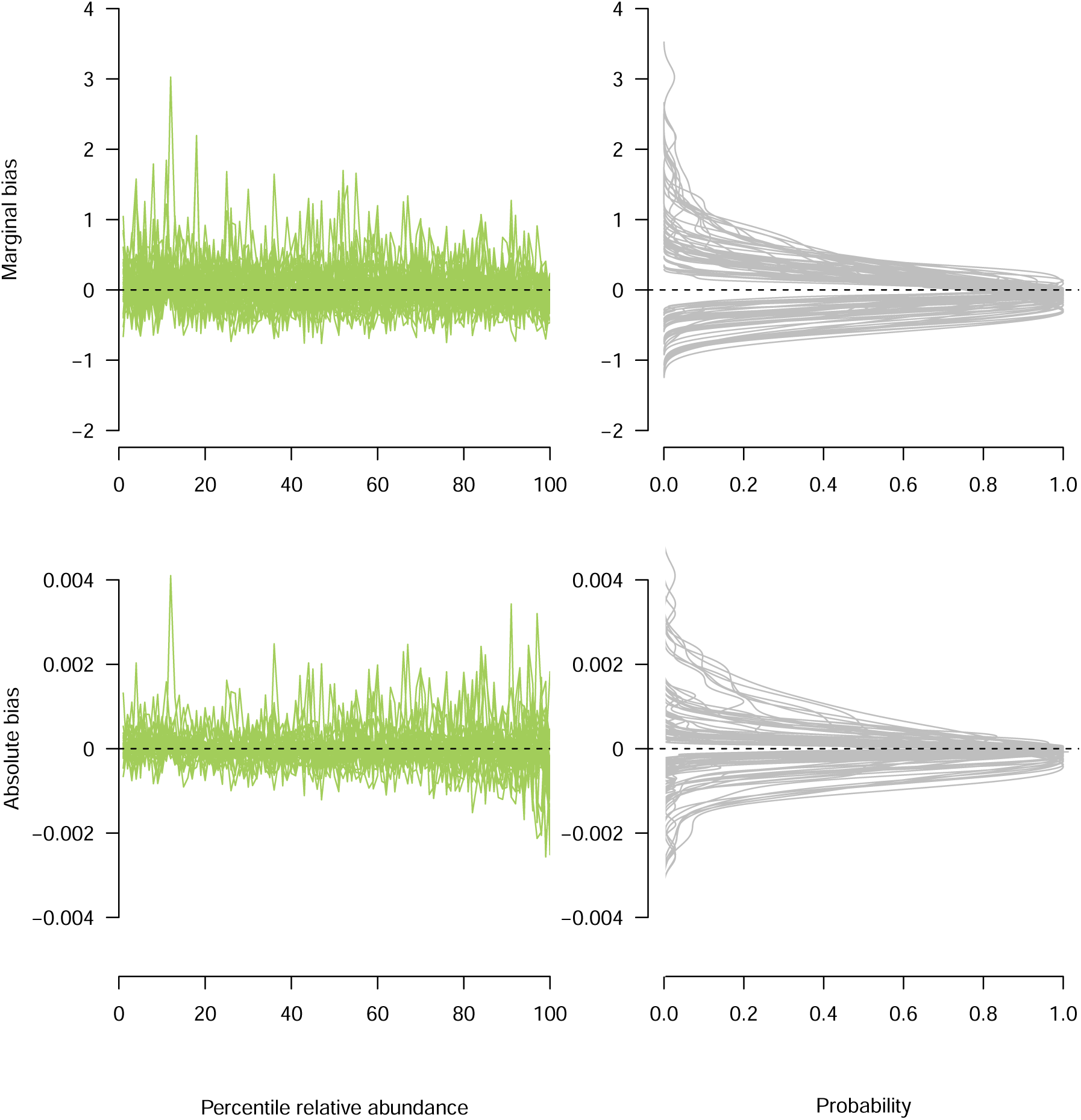
Bias of DMM as implemented via MCMC (via JAGS) as a function of feature relative abundance for data sets simulated using a moderately skewed rank abundance distribution (Pareto = 4). Percentiles of relative abundances for each dataset were calculated and are shown on the x axis of bias plots (left column), thus normalizing for the differences in numbers of features among datasets. Bias (defined as the difference between predicted values and the truth) in *π* parameters is shown on the y axis. Marginal bias was calculated as absolute bias divided by the relative abundance of the focal parameter. Plots in the right column show probability densities for the different bias values shown in the left column. Each line denotes results from a simulated data set. Results shown here are from a representative subset of the datasets simulated as part of our main experiment. Results shown here are from a representative subset of the datasets simulated as part of our main experiment including 99 datasets with variable counts among replicates (see Methods) and 45 datasets with invariant counts among replicates. All values that we considered as part of our main experiment for number of features, number of observations, number of replicates, precision (*θ*), and effect sizes were included in the subset of the datasets analyzed here.

**Figure S15:**
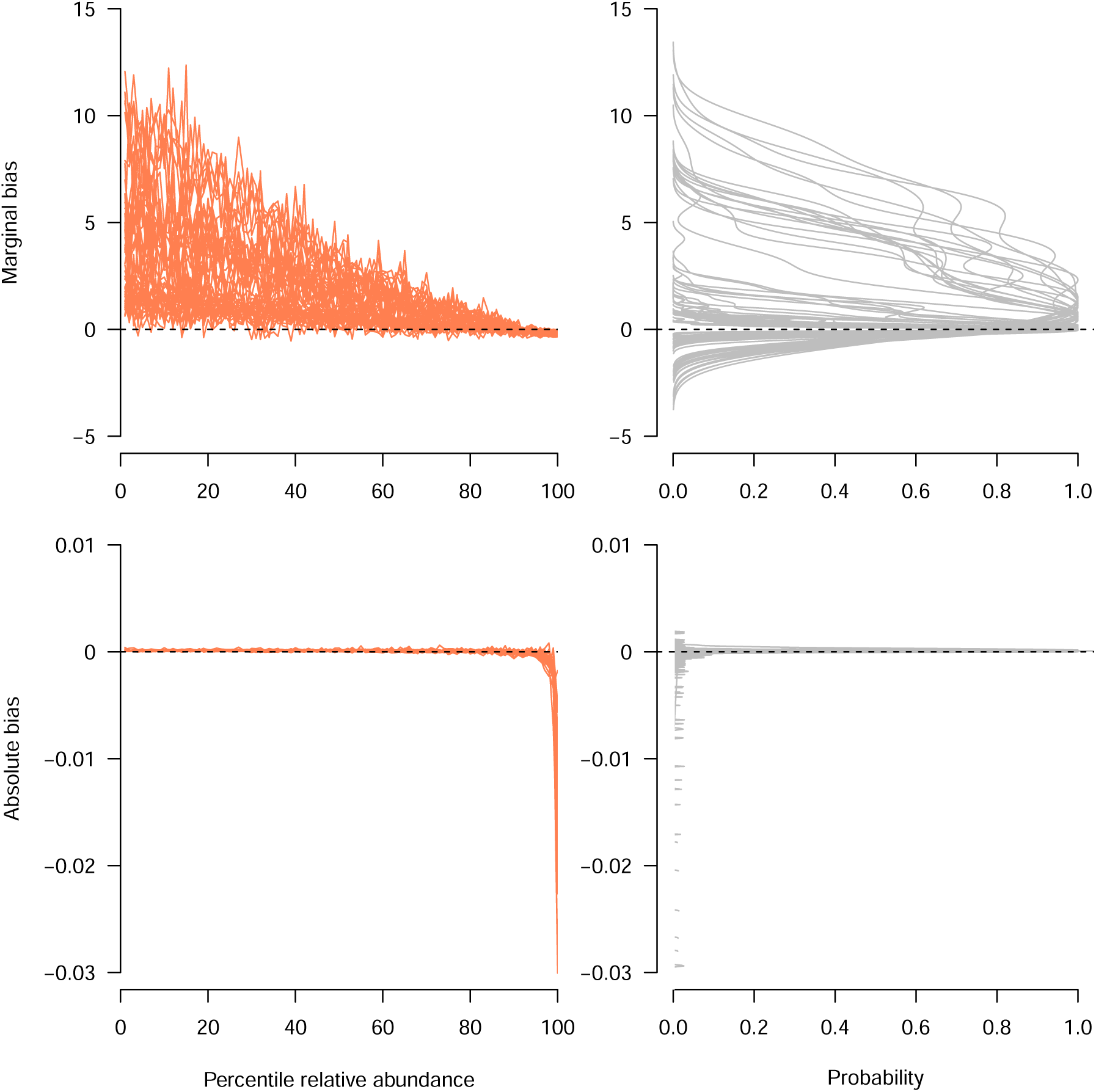
Bias of DMM as implemented via MCMC (via JAGS) as a function of feature relative abundance for data sets simulated using a highly skewed rank abundance distribution (Pareto = 0.7). Percentiles of relative abundances for each dataset were calculated and are shown on the x axis of bias plots (left column) of bias plots (left column), thus normalizing for the differences in numbers of features among datasets. Bias (defined as the difference between predicted values and the truth) in *π* parameters is shown on the y axis. Plots in the right column show probability densities for the different bias values shown in the left column. Plots in the right column show probability densities for the different bias values shown in the left column. Each line denotes results from a simulated data set. Results shown here are from a representative subset of the datasets simulated as part of our main experiment including 99 datasets with variable counts among replicates (see Methods) and 45 datasets with invariant counts among replicates. All values that we considered as part of our main experiment for number of features, number of observations, number of replicates, precision (*θ*), and effect sizes were included in the subset of the datasets analyzed here.

**Figure S16:**
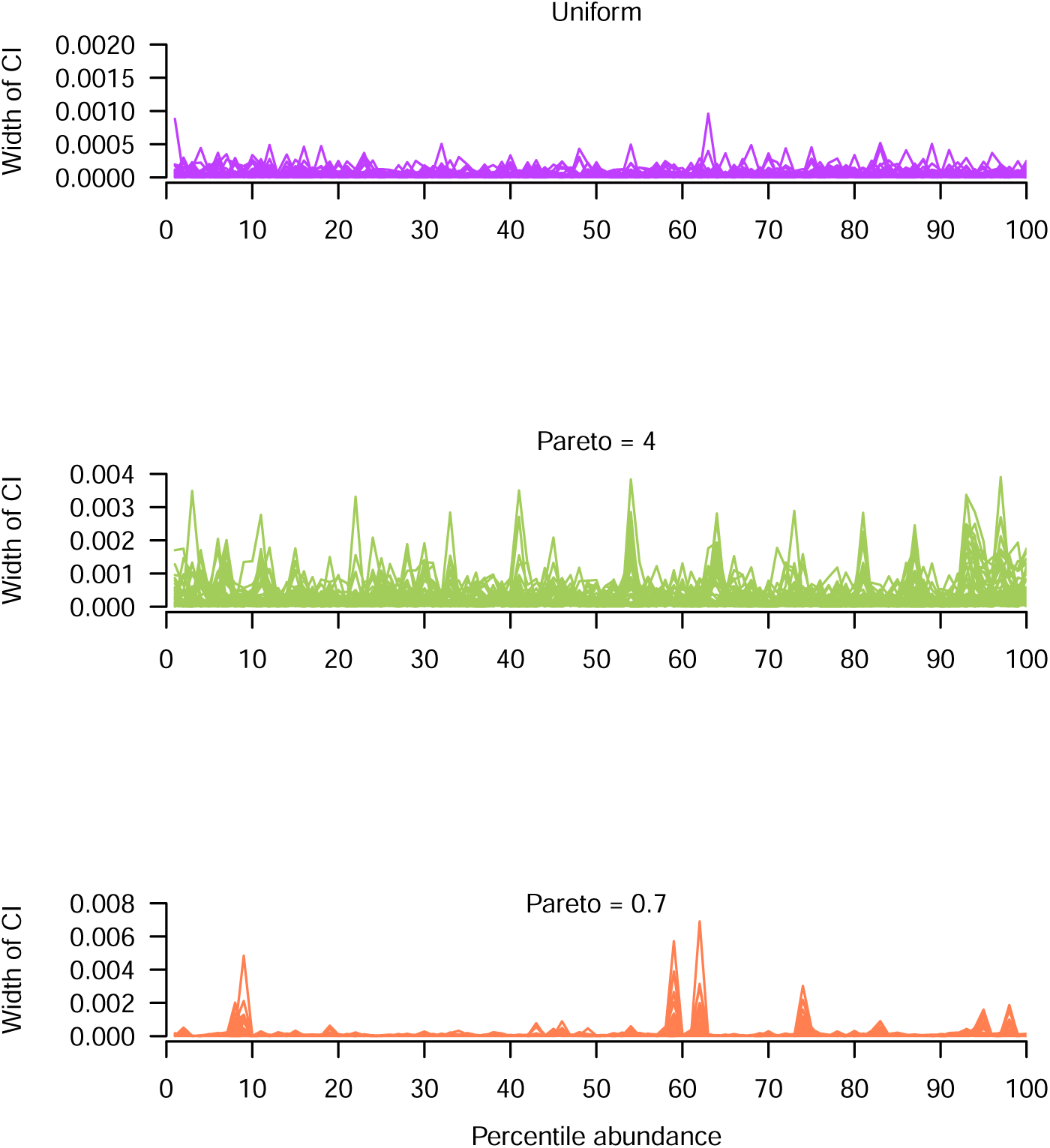
Width of credible intervals for *π* parameters estimated using DMM as implemented via HMC (using the Stan software) as a function of feature relative abundance and rank abundance profile of the data (see Fig. 2). Percentiles of relative abundances for each dataset were calculated and are shown on the x axis of bias plots (left column), thus normalizing for the differences in numbers of features among datasets. Credible interval width (defined as the absolute value of the difference between the 2.5 and 97.5 percentiles of the estimated posterior probability distribution) of *π* parameters is shown on the y axis. Plots in the right column show probability densities for the different bias values shown in the left column. Each line denotes results from a simulated data set. Results shown here are from a representative subset of the datasets simulated as part of our main experiment including 99 datasets with variable counts among replicates (see Methods) and 45 datasets with invariant counts among replicates. All values that we considered as part of our main experiment for number of features, number of observations, number of replicates, precision (*θ*), and effect sizes were included in the subset of the datasets analyzed here.

**Figure S17:**
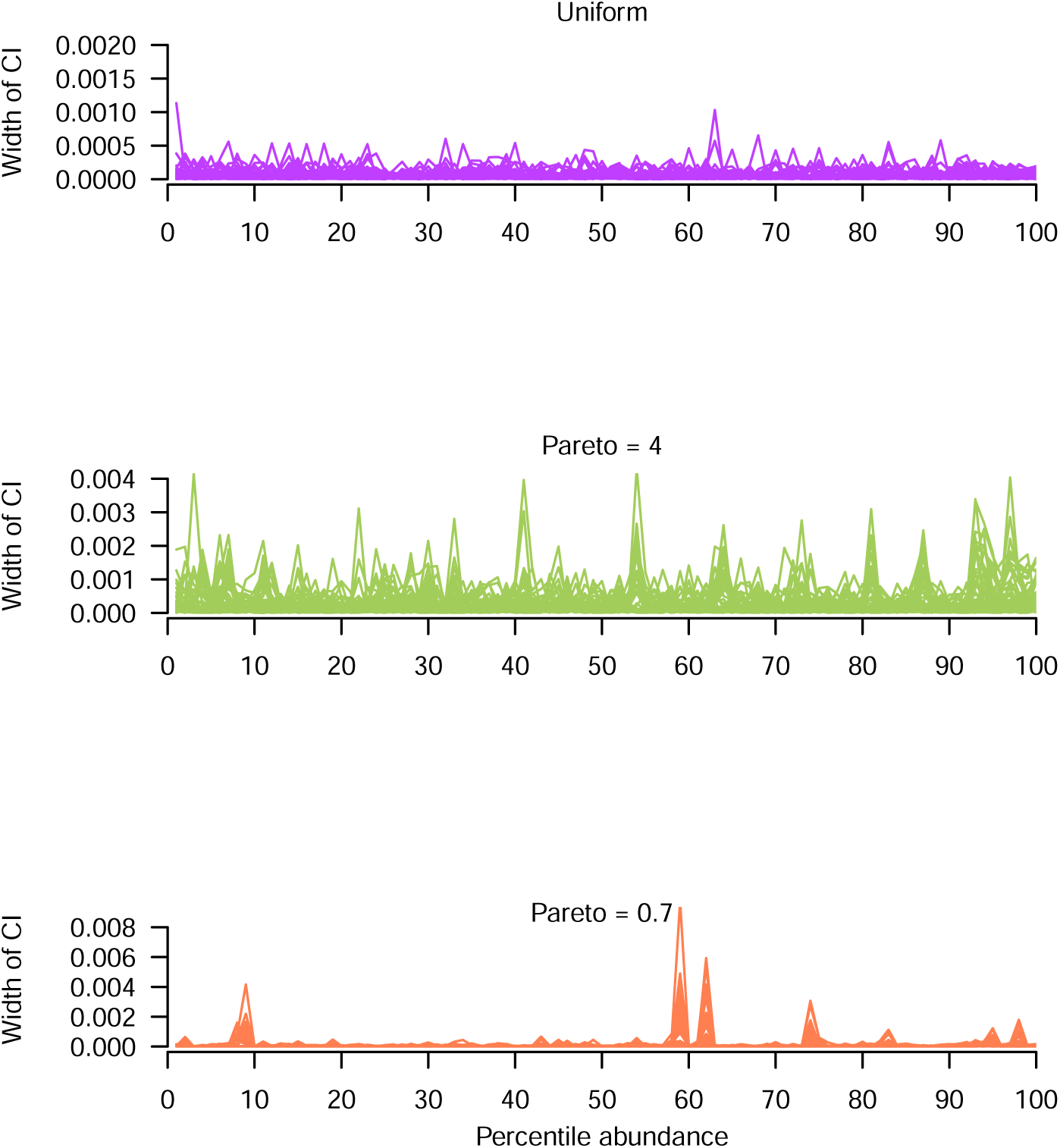
Width of credible intervals for *π* parameters estimated using DMM as implemented via VI (using the Stan software) as a function of feature relative abundance and rank abundance profile of the data (see Fig. 2). Percentiles of relative abundances for each dataset were calculated and are shown on the x axis of bias plots (left column), thus normalizing for the differences in numbers of features among datasets. Credible interval width (defined as the absolute value of the difference between the 2.5 and 97.5 percentiles of the estimated posterior probability distribution) of *π* parameters is shown on the y axis. Results shown here are from a representative subset of the datasets simulated as part of our main experiment including 99 datasets with variable counts among replicates (see Methods) and 45 datasets with invariant counts among replicates. All values that we considered as part of our main experiment for number of features, number of observations, number of replicates, precision (*θ*), and effect sizes were included in the subset of the datasets analyzed here.

**Figure S18:**
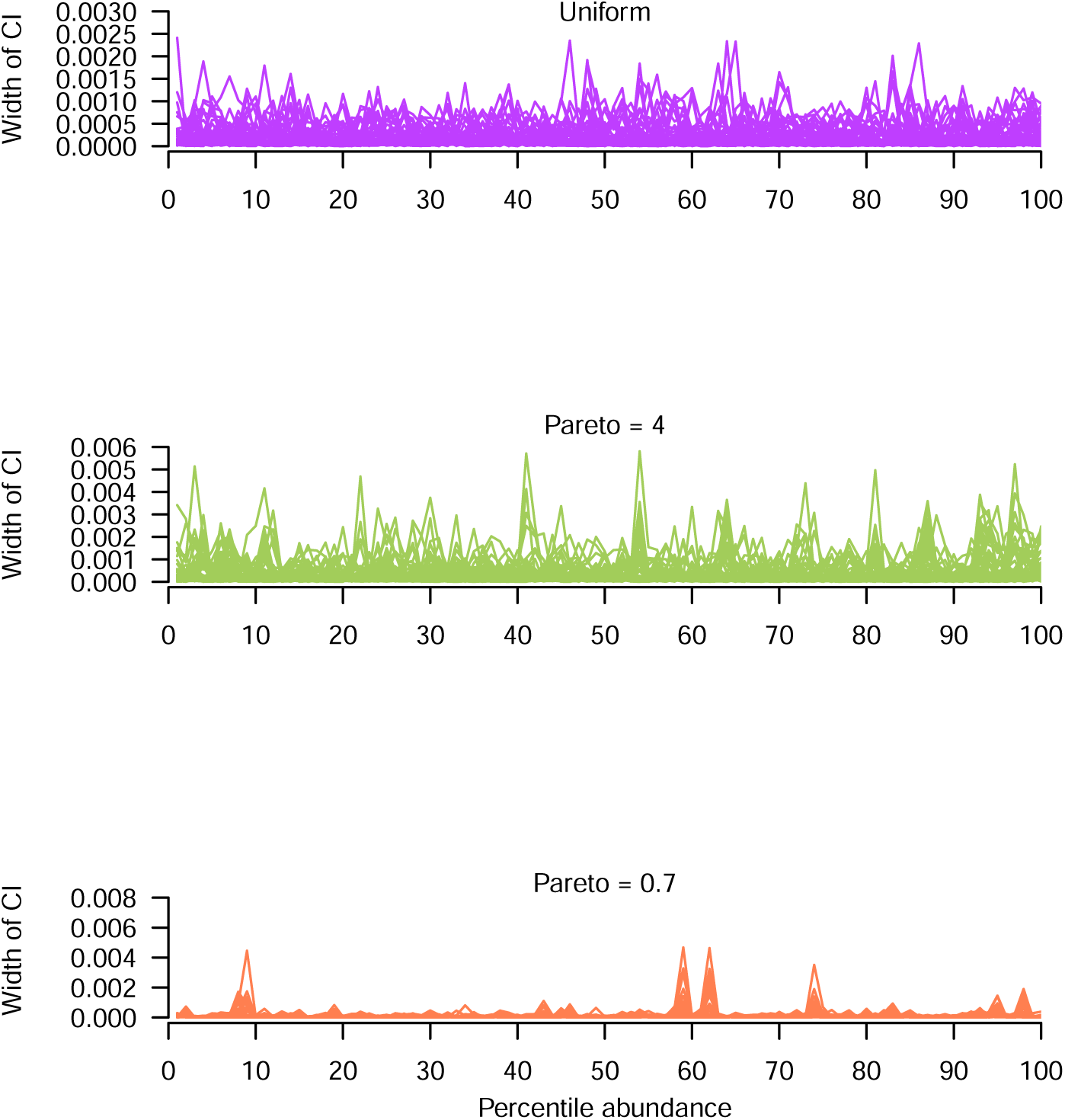
Width of credible intervals for *π* parameters estimated using DMM as implemented via MCMC (using the JAGS software) as a function of feature relative abundance and rank abundance profile of the data (see Fig. 2). Percentiles of relative abundances for each dataset were calculated and are shown on the x axis of bias plots (left column), thus normalizing for the differences in numbers of features among datasets. Credible interval width (defined as the absolute value of the difference between the 2.5 and 97.5 percentiles of the estimated posterior probability distribution) of *π* parameters is shown on the y axis. Plots in the right column show probability densities for the different bias values shown in the left column. Each line denotes results from a simulated data set. Results shown here are from a representative subset of the datasets simulated as part of our main experiment including 99 datasets with variable counts among replicates (see Methods) and 45 datasets with invariant counts among replicates. All values that we considered as part of our main experiment for number of features, number of observations, number of replicates, precision (*θ*), and effect sizes were included in the subset of the datasets analyzed here.

**Figure S19:**
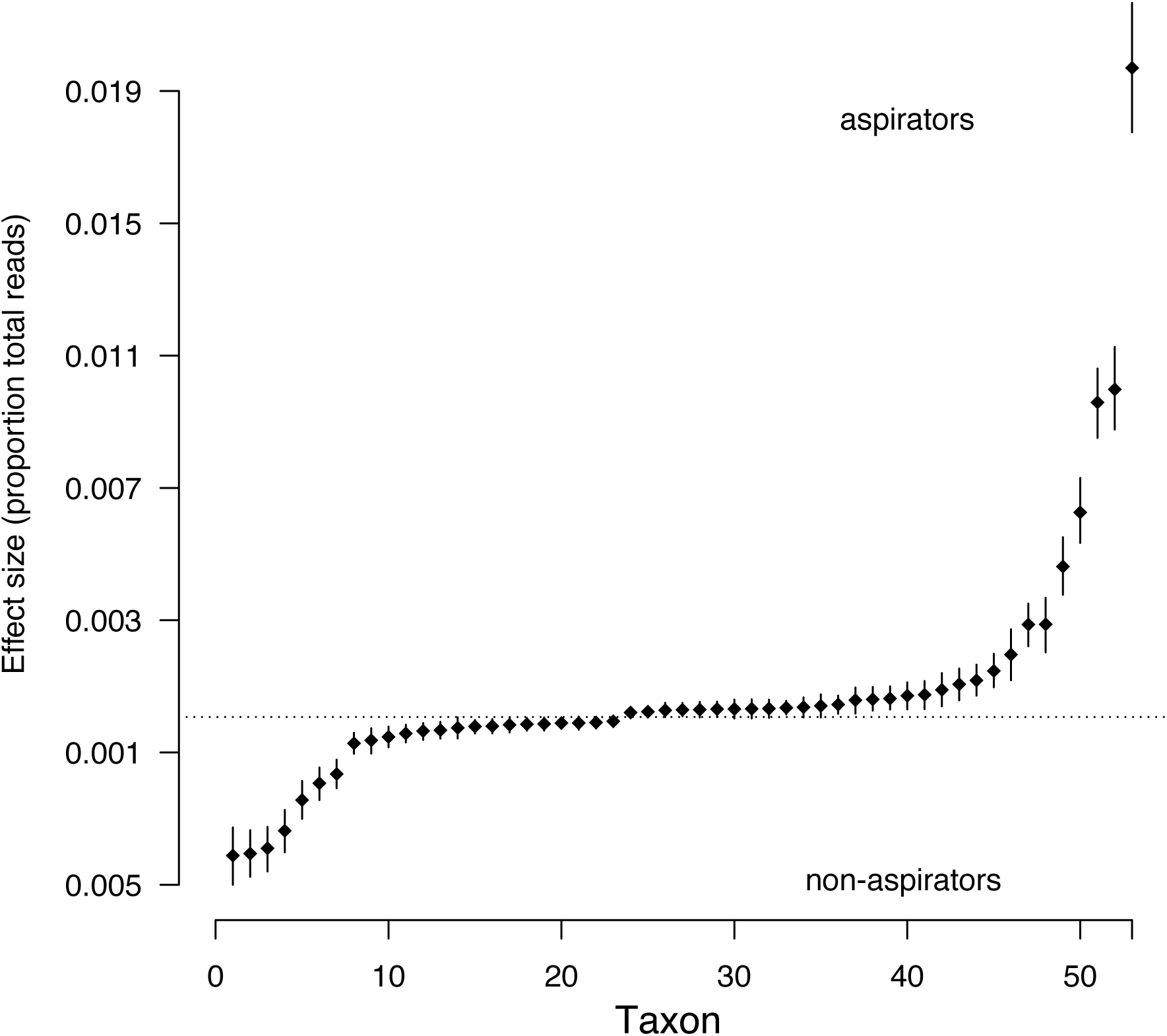
Lung inhabiting bacteria that shifted in relative abundance between aspirating and non-aspirating subjects. Data analyzed were made publicly available by Duvallet et al. (2019). The estimated relative abundance of taxa in non-aspirating patients (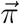 parameters) was subtracted from the relative abundance of taxa in aspirating patients and this difference is shown on the vertical axis. Thus, points on either side of zero (shown as absolute values) correspond with a taxon that was more abundant in that sampling group (non-aspirators below zero, aspirators above zero). Points are means of PPDs of differences; whiskers show the 95% equal tailed probability intervals of PPDs. Bacteria are indexed along the horizontal axis and ordered by effect size.

When a finite number of observations can be ascribed to categories (e.g., observations of taxa or transcripts), the counts of observations of each category can be appropriately modeled using the multinomial distribution. Multinomial parameters define the probability that a given observation belongs to a particular category and these probabilities correspond to the relative abundance of that category in the population that was sampled. Because it accounts for the probability of all categories, the sum of the multinomial parameter vector 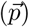 is one. For instance, if 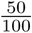 of the birds one observed on a long hike were American robins then the maximum likelihood estimate of the multinomial parameter for robins would be 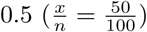 and other parameters would correspond to the relative abundance of the other bird taxa observed. Here we describe how to model multinomial data using a hierarchical Bayesian approach that shares information among replicates via the Dirichlet distribution. The parameters of the Dirichlet distribution allow inference regarding the relative abundance of each category, or feature, within the sampling group.

The goal of the analysis demonstrated here is to identify features (i.e. taxa, transcripts, behavioral preferences) that differ in relative abundance across treatment groups. However, once estimates for feature relative abundance are obtained, these estimates can be passed to additional analyses. We implement modeling using three frameworks (variational inference and Hamiltonian Monte Carlo in Stan, and MCMC [Gibbs and Metropolis-Hastings] sampling in JAGS) to demonstrate the differences and similarities of each.

Be advised that modeling large data sets is computationally expensive, therefore we use a simple, simulated data set for this example. For smaller datasets, say of a few hundred to a thousand features, the model shown here can be run on a desktop system. For larger datasets, computation will take several days, so one may wish to run the model remotely. One trick that can be used to reduce computational expense is to sum uncommon features into a single, composite feature. The counts of this composite feature should be included during modeling, otherwise proportional estimates will be incorrect. Also, run time can be reduced by initializing sampling at values that are likely to be closer to the true values of the parameters to be estimated (e.g., *π* parameters in the Dirichlet could be set to the maximum likelihood estimate of the frequency of that feature across replicates). See documentation for Stan or rjags for information on how to initialize chains.

## Simulation

We start this example by simulating some data. Note that the intensity parameter of the Dirichlet distribution controls the degree of among-replicate variation within the data. Higher values for this parameter lead to less variation among replicates. Also, we add a one to every datum so that there are no zero values within the data. This is necessary because zeros can cause infinite density errors in JAGS, due to their contribution to the Dirichlet probability density function. If zeros exist in one’s data, then add a one to every count.

~~~
*# library(gtools)*
*# library(rstan)*
*# library(rjags)*
*# library(shinystan)*
*# library(VGAM)*

notus <- 50
nsamples <- 5000
nreps <- 100
intensity <- 1
comprop <- **matrix**(0, ncol = notus, nrow = 2)
indprop <- **matrix**(0, ncol = notus, nrow = nreps)

*#Assemblage 1*
comprop[1, ] <- **rdirichlet**(1, **c**(**rep**(15, 5), **rep**(1, notus **-** 5)))
*#Assemblage 2*
comprop[2, ] <- **rdirichlet**(1, **c**(**rep**(1, notus **-** 5), **rep**(15, 5)))

*#Construct data matrix*
com <- **matrix**(0, ncol = notus, nrow = nreps)
**for** (i **in** 1:(nreps **/** 2)) {
indprop[i, ] <- **rdirichlet**(1, comprop[1, ] **\*** intensity)
com[i, ] <- **rmultinom**(1, nsamples, prob = indprop[i, ])
}
**for** (i **in** (1 **+** nreps **/** 2): nreps) {
indprop[i, ] <- **rdirichlet**(1, comprop[2, ] **\*** intensity)
com[i, ] <- **rmultinom**(1, nsamples, prob = indprop[i, ])
}
com <- com **+** 1
nsamples <- nsamples **+** 50
~~~

## Stan model specification

Now we run the model. See the main text for model exposition. First, we load the Stan specification of the model, which, in this case, is in a text file located within the working directory. This can take a few seconds.

~~~
DM <- **stan_model**(“DM.stan”, model_name = “DM”)
*#This file has the following model within it:*

*# // Model specification for Dirichlet-Multinomial*
*# data {*
*#   int<lower=1> N;*
*#   int<lower=1> nreps;*
*#   int<lower=1> notus;*
*#*
*#   int<lower=1> start[N];*
*#   int<lower=1> end[N];*
*#*
*#   int datamatrix[nreps, notus];*
*# }*
*#*
*# parameters {*
*#   real<lower=0> theta[N];*
*#   simplex[notus] pi[N];*
*#   simplex[notus] p[nreps];*
*# }*
*#*
*#*
*# model {*
*#   for(i in 1:N){*
*#     target += exponential_lpdf(theta[i] | 0.001);*
*#     target += dirichlet_lpdf(pi[i] | rep_vector(0.0000001, notus));*
*#     for(j in start[i]:end[i]){*
*#      target += dirichlet_lpdf(p[j] | theta[i]*pi[i]);*
*#      target += multinomial_lpmf(datamatrix[j,] | p[j]);*
*#    }*
*#   }*
*# }*
~~~

## Variational inference in Stan

Now we implement variational inference (VI) to learn parameters of interest. Note how the data are passed in as a named list, the algorithm specified, and the number of samples to be extracted from the estimated posterior specified (“output samples”). For more, see the Stan documentation.

~~~
ptm <- **proc.time**()
fitstan_VI <- **vb**(DM,
       data = **list**(“datamatrix” = com,
               “nreps” = **nrow**(com),
               “notus” = **ncol**(com),
               “N” = 2,
               “start” = **c**(1, nreps**/**2),
               “end” = **c**((nreps**/**2) **-** 1, nreps)
              ),
     algorithm = “meanfield”,
     output_samples = 500,
     check_data = T,
     seed = 123,
     pars <- “pi”)
viTime <- **c**(**proc.time**() **-** ptm)[3]
~~~

Variational inference took 5.959 seconds.

## Hamiltonian Monte Carlo sampling in Stan

Now we implement Hamiltonian Monte Carlo (HMC) using the no U-turn sampling algorithm. Note that the number of chains and cores can be specified (use one core per chain). “warmup” controls model burn in (and should probably be increased for larger data sets). “iter” controls total iterations, so the difference between iter and warmup specifies how many samples of the posterior probability distribution will be extracted. “thin” specifies how many samples to skip before saving another sample (if thin=2 then every other sample will be saved). For more, see the Stan documentation.

~~~
ptm <- **proc.time**()
fitstan_HMC <- **sampling**(DM,
     data = **list**(“datamatrix” = com,
         “nreps” = **nrow**(com),
         “notus” = **ncol**(com),
         “N” = 2,
         “start” = **c**(1, nreps**/**2),
         “end” = **c**((nreps**/**2) **-** 1, nreps)
     ),
     chains=2,
     warmup = 500,
     iter = 1000,
     thin = 2,
     algorithm = “NUTS”,
     cores = 1,
     pars <- “pi”,
     verbose = T)
hmcTime <- **c**(**proc.time**() **-** ptm)[3]
~~~

Hamiltonian Monte Carlo took 278.297 seconds. Note that this time could be reduced by optimizing “warmup” and “iter”, running each chain on a different core, and providing sensible initialization values. When optimizing run time be sure to check model convergence statistics to ensure that convergence upon a stable posterior probability distribution has been achieved.

## Stan estimation diagnostics

Checking model convergence can be done easily for HMC, but at the time of writing there was no simple way to test effectiveness of VI.

For HMC, the number of effective samples and 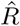 can be checked using the following code.

~~~
**summary**(fitstan_HMC, pars = “pi”, probs = **c**(0.025, 0.975))**$**summary
~~~

The shinystan application is an excellent interface to dig deeper into model performance. See https://mc-stan.org/users/interfaces/shinystan

## Model specification and MCMC samples in JAGS

Now we use a very similar specification of the model for the JAGS software to estimate *π* parameters of the Dirichlet.

Model specification is as follows:

~~~
community.model.level <- “model{
  for(i in 1:N){
   for(j in start[i]:end[i]){
    datamatrix[j,] ∼ dmulti(p[j,], nreads[j])
    p[j,1:notus] ∼ ddirch(pi[i,]*theta[i])
   }
   pi[i,1:notus] ∼ ddirch(alpha)
   theta[i] ∼ dunif(0, 4000)
}
  for(k in 1:notus){
    alpha[k] <- 0.0000001
}
}”
~~~

Compile and run the model.

~~~
ptm <- **proc.time**()

sim.mod.jags <- **jags.model**(
  **textConnection**(community.model.level),
  data = **list**(
   datamatrix = com,
   notus = **dim**(com)[2],
   nreads = **rowSums**(com),
   N = 2,
   start = **c**(1,nreps**/**2),
   end = **c**((nreps**/**2)**-**1,nreps)
),
   n.chains = 2,
   n.adapt = 0
)

*#Adapt model*
iter_needed <- 0
y = FALSE
**while**(y **==** FALSE){
  y <- **adapt**(sim.mod.jags,
          n.iter = 1000,
          end.adaptation = FALSE)
   iter_needed <- 1000 **+** iter_needed
   **if**(iter_needed **>** 4000){**break**}
}

*#Burn in*
**update**(sim.mod.jags,
         n.iter = 3000)

*#Extract samples*
sim.mod.sam <- **jags.samples**(model = sim.mod.jags,
              variable.names = “pi”,
              n.iter = 4000,
              thin = 4)
jagsTime <- **c**(**proc.time**() **-** ptm)[3]
~~~

JAGS took 77.208 seconds. This time could possibly be reduced by optimizing burn in and adaptation and providing sensible initialization values.

To test for MCMC convergence, one can use the functions within the Coda R package. Be advised, that statistics should be calculated parameter-wise when there are many parameters, else memory requirements become burdensome. The following function can be used to accomplish this task.

~~~
*#Compute the Gelman-Rubin and Geweke statistics*
mcmcdiag <- **function**(x, nparams) {
 *#x is an mcmc object*
 *#nparams is number of params in the object*
 Gr <- **vector**(length = nparams)
 GK <- **vector**(length = nparams)
 k <- 1
 a <- **character**(0)

 **while** (k **<=** nparams) {
  m <- x[1**:length**(x)][, k]
  gr <- **gelman.diag**(m)
  **print**(**paste**(“Feature”, k, sep = “ “))
  **print**(“Gelman-Rubin”)
  **print**(gr)
  **if** (gr[[1]][1] **<=** 2) {
   Gr[k] <- “passed”
  } **else**{
   Gr[k] <- “failed”
  }
  gk <- **geweke.diag**(m,
     frac1 = 0.1,
     frac2 = 0.5)
  suspectGK <- **names**(**which**(2 **\* pnorm**(**-abs**(gk[[1]]**$**z)) **<** 0.08))
  **if** (**identical**(a, suspectGK)) {
   GK[k] <- “passed”
   } **else if** (suspectGK **==** “var1”) {
GK[k] <- “failed”
   }
   k <- k **+** 1
  }
  **return**(**list**(Gr,
             GK))
}
diagout <- **mcmcdiag**(**as.mcmc.list**(sim.mod.sam**$**pi), **dim**(com)[2])
~~~

We have noticed that for large datasets (many thousands of parameters), JAGS can require many days to achieve convergence. By comparison, HMC is much faster. To avoid impractically long run times, VI may be the only viable option for extremely large data sets.

## Use of parameter estimates

Now we extract *π* parameters from each sampling group and subtract them. The location of zero within this distribution quantifies the probability of no effect of sampling group. If desired, the mean of this distribution of differences can be extracted and used as a point estimate for the effect of sampling group, though we advocate for using samples characterizing the entire distribution for analyses whenever possible, thus utilizing our measures of uncertainty. We present a simple function to determine if 95% or more of the distribution of differences lies on either side of zero. If so, then we suggest this is high certainty of an effect of sampling group on the relative abundance of that feature.

~~~
calc_certain_diffs <- **function**(mcmc_of_diffs, dimension){
  positives <- **vector**()
  negatives <- **vector**()
  **for**(i **in** 1**:dim**(mcmc_of_diffs)[dimension]){
   **if**(dimension **==** 2){
    perc <- **length**(**which**(mcmc_of_diffs[,i] **>** 0))**/ length**(mcmc_of_diffs[,i])
   }**else**{
    perc <- **length**(**which**(mcmc_of_diffs[i,] **>** 0)) **/ length**(mcmc_of_diffs[i,])
   }
   **if**(perc **>=** 0.95 **|** perc **<=** 0.05){
   positives <- **c**(positives, i)
 }**else**{
   negatives <- **c**(negatives, i)
   }
 }
   **return**(**list**(positives = positives,
           negatives = negatives))
}
 est.pi <- **extract**(fitstan_HMC,”pi”)
 diffs_HMC <- est.pi**$**pi[,1,] **-** est.pi**$**pi[,2,]
 outHMC <- **calc_certain_diffs**(diffs_HMC,2)
 est.pi <- **extract**(fitstan_VI,”pi”)
 diffs_VI <- est.pi**$**pi[,1,] **-** est.pi**$**pi[,2,]
 outVI <- **calc_certain_diffs**(diffs_VI,2)
 diffs_jags <- sim.mod.sam**$**pi[1,,,1: 2] **-** sim.mod.sam**$**pi[2,,,1: 2]
 outJAGS <- **calc_certain_diffs**(**cbind**(diffs_jags[,,1],
                                     diffs_jags[,,2]), 1)
~~~

Next we make a plot to determine which features shifted in relative abundances. Points correspond to estimated differences in feature relative abundance between sampling groups. The blue dots correspond with those features that we expected to shift. Lines extending from each point denote 95% high density intervals, and are colored purple for those features suggested to differ.

~~~
*#Code from Kruschke’s Doing Bayesian Data Analysis book (cited in main text).*
HDIofMCMC = **function**(sampleVec, credMass=0.95) {
*# Computes highest density interval from a sample of representative values*,
*# estimated as shortest credible interval.*
*# Arguments:*
*# sampleVec*
*# is a vector of representative values from a probability distribution.*
*# credMass*
*# is a scalar between 0 and 1, indicating the mass within the credible*
*# interval that is to be estimated.*
*# Value:*
*# HDIlim is a vector containing the limits of the HDI*
sortedPts = **sort**(sampleVec)
  ciIdxInc = **ceiling**(credMass **\* length**(sortedPts))
nCIs = **length**(sortedPts) **-** ciIdxInc
ciWidth = **rep**(0, nCIs)
**for**(i **in** 1: nCIs) {
  ciWidth[i]= sortedPts[i **+** ciIdxInc] **-** sortedPts[i]
}
HDImin = sortedPts[**which.min**(ciWidth)]
HDImax = sortedPts[**which.min**(ciWidth) **+** ciIdxInc]
HDIlim = **c**(HDImin, HDImax)
**return**(HDIlim)
}
notus <- **dim**(com)[2]
colorPoints <- **rep**(“black”, notus)
colorPoints[**c**(1: 5,(notus**-**4): notus)] <- “blue”
*#Plot differences in pis*
plotr <- **function**(x, y, z, whatitis){
**plot**(1: notus, **apply**(x, y, mean),
   cex = 1.5,
   ylim = **c**(**-**0.06,0.06),
   ylab = “Difference in rel. abund.”,
   xlab = “Feature”,
   main = whatitis,
   pch = 16,
   col = colorPoints,
   las = 2)
**abline**(h = 0, col = “red”)
segs <- **apply**(x, y, HDIofMCMC)
colorLines <- **rep**(“black”, notus)
colorLines[z**$**positives] <- “purple”
**segments**(1: notus, segs[1,],
      1: notus, segs[2,],
      col = colorLines)
}
**par**(mfrow=**c**(1,3))
**plotr**(x = diffs_VI, y = 2, z = outVI, whatitis = “VI”)
**plotr**(x = diffs_HMC, y = 2, z = outHMC, whatitis = “HMC”)
**plotr**(x = diffs_jags, y = 1, z = outJAGS, whatitis = “JAGS”)
~~~

DMM can be extended easily to encompass more than two sampling groups. Simply order data (in a matrix or dataframe format) so that replicates from the same sampling groups are neighboring rows. For instance, say one was analyzing measurements from eight sampling groups denoted numerically. One should order the associated data for these sampling groups like so that the data looked like this:

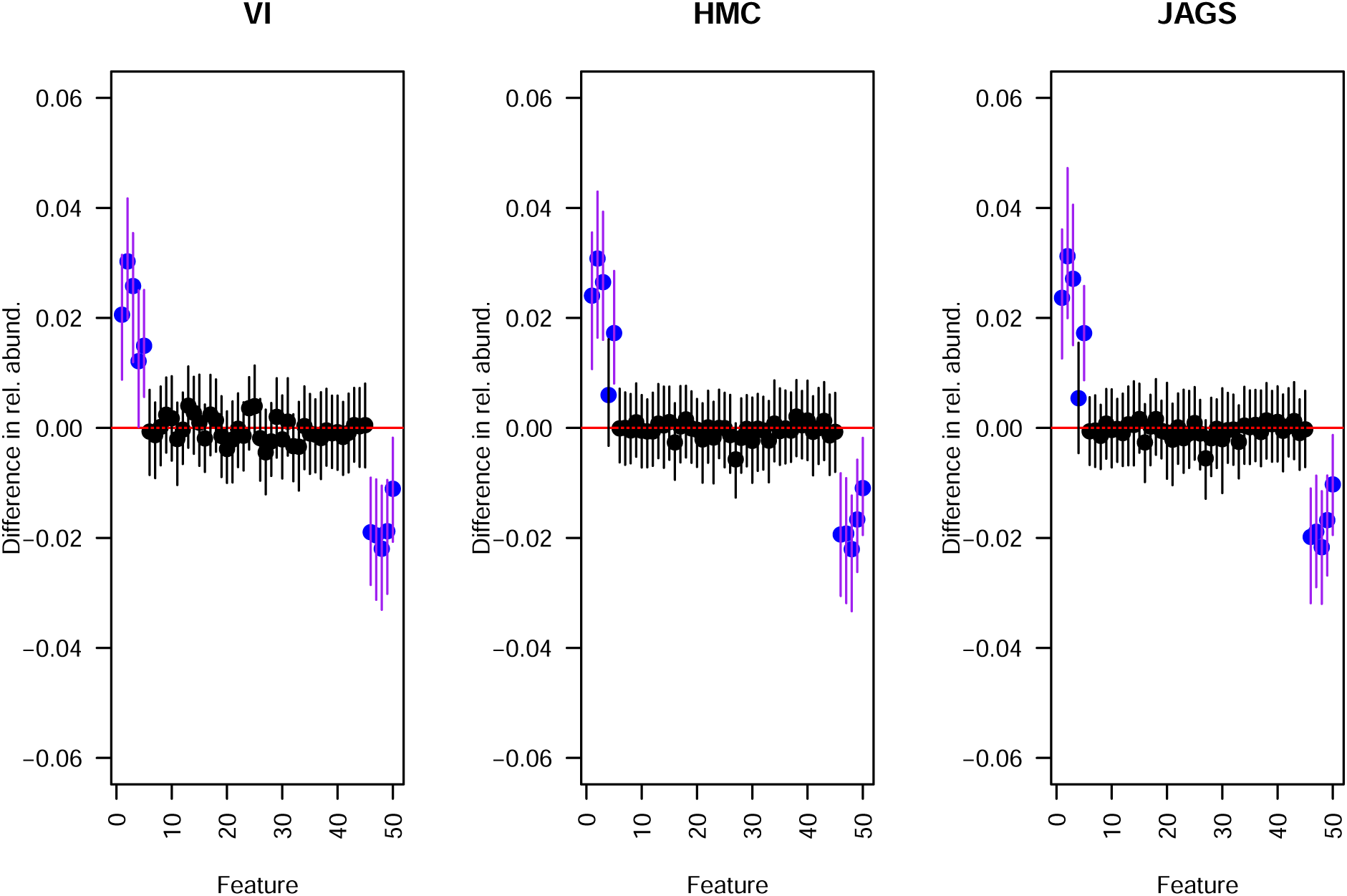

~~~
exampleData <- **round**(**runif**(16,1,1000))
groups <- **c**(**rep**(“group1”,2),
    **rep**(“group2”,2),
    **rep**(“group3”,2),
    **rep**(“group4”,2),
    **rep**(“group5”,2),
    **rep**(“group6”,2),
    **rep**(“group7”,2),
    **rep**(“group8”,2))
**cbind**(exampleData, groups)
## exampleData groups
## [1,] “766” “group1”
## [2,] “347” “group1”
## [3,] “134” “group2”
## [4,] “686” “group2”
## [5,] “90” “group3”
## [6,] “539” “group3”
## [7,] “244” “group4”
## [8,] “799” “group4”
## [9,] “232” “group5”
## [10,] “349” “group5”
## [11,] “751” “group6”
## [12,] “936” “group6”
## [13,] “586” “group7”
## [14,] “201” “group7”
## [15,] “940” “group8”
## [16,] “865” “group8”
~~~

Then one can simply pass in the indices that describe which rows bound which group to the “start” and “end” portions of the function. For our toy example, the start indices would be:

~~~
**c**(1,3,5,7,9,11,13,15)
## [1] 1 3 5 7 9 11 13 15
~~~

and the end indices would be:

~~~
**c**(2,4,6,8,10,12,14,16)
## [1] 2 4 6 8 10 12 14 16
~~~

These values would then be substituted into the model and the “N” parameter changed to reflect the number of sampling groups used (in this case N = 8). Note that you cannot pass in the grouping column if it is included in your data. See above for another example of how data should be formatted.

~~~
fitstan_HMC <- **sampling**(DM,
      data = **list**(“datamatrix” = **as.matrix**(exampleData),
        “nreps” = 16,
        “notus” = 1,
        “N” = 8,
        “start” = **c**(1,3,5,7,9,11,13,15),
        “end” = **c**(2,4,6,8,10,12,14,16)
     ),
     chains=2,
     warmup = 500,
     iter = 1000,
     thin = 2,
     algorithm = “NUTS”,
     cores = 1,
     pars <- “pi”,
     verbose = T)
~~~

## References

Aitchison, J. (1982). The statistical analysis of compositional data. Chapman and Hall, New York, NY. CITE.

Aitchison, J. and Egozcue, J. J. (2005). Compositional data analysis: where are we and where should we be heading? Mathematical Geology, 37(7):829–850.

Akata, K., Yatera, K., Yamasaki, K., Kawanami, T., Naito, K., Noguchi, S., Fukuda, K., Ishimoto, H., Taniguchi, H., and Mukae, H. (2016). The significance of oral streptococci in patients with pneumonia with risk factors for aspiration: the bacterial floral analysis of 16s ribosomal RNA gene using bronchoalveolar lavage fluid. BMC Pulmonary Medicine, 16(1):79.

Bates, D., Mächler, M., Bolker, B., and Walker, S. (2015). Fitting linear mixed-effects models using lme4. Journal of Statistical Software, 67(1):1:48.

Beck, J. M., Young, V. B., and Huffnagle, G. B. (2012). The microbiome of the lung. Translational Research, 160(4):258–266.

Björk, J. R., Hui, F. K. C., O’Hara, R. B., and Montoya, J. M. (2018). Uncovering the drivers of host-associated microbiota with joint species distribution modelling. Molecular Ecology, 27(12):2714–2724.

Blei, D. M., Kucukelbir, A., and McAuliffe, J. D. (2017). Variational inference: a review for statisticians. Journal of the American Statistical Association, 112(518):859–877.

Buckland, M. and Gey, F. (1994). The relationship between Recall and Precision. Journal of the American Society for Information Science, 45(1):12–19.

Bullard, J. H., Purdom, E., Hansen, K. D., and Dudoit, S. (2010). Evaluation of statistical methods for normalization and differential expression in mRNA-Seq experiments. BMC Bioinformatics, 11(1):94.

Carbonetto, P. and Stephens, M. (2012). Scalable variational inference for Bayesian variable selection in regression, and its accuracy in genetic association studies. Bayesian Analysis, 7(1):73–108.

Carpenter, B., Gelman, A., Hoffman, M. D., Lee, D., Goodrich, B., Betancourt, M., Brubaker, M., Guo, J., Li, P., and Riddell, A. (2017). Stan: a probabilistic programming language. Journal of Statistical Software, 76(1):1–32.

Chen, J. and Li, H. (2013). Variable selection for sparse Dirichlet-multinomial regression with an application to microbiome data analysis. The Annals of Applied Statistics, 7(1):418–442.

Claesson, B. A. and Leinonen, M. (1994). *Moraxella catarrhalis* — an uncommon cause of community-acquired pneumonia in Swedish children. Scandinavian Journal of Infectious Diseases, 26(4):399–402.

Coblentz, K. E., Rosenblatt, A. E., and Novak, M. (2017). The application of Bayesian hierarchical models to quantify individual diet specialization. Ecology, 98(6):1535–1547.

Dillies, M.-A., Rau, A., Aubert, J., Hennequet-Antier, C., Jeanmougin, M., Servant, N., Keime, C., Marot, G., Castel, D., Estelle, J., Guernec, G., Jagla, B., Jouneau, L., Laloë, D., Le Gall, C., Schaëffer, B., Le Crom, S., Guedj, M., and Jaffrézic, F. (2013). A comprehensive evaluation of normalization methods for Illumina high-throughput RNA sequencing data analysis. Briefings in Bioinformatics, 14(6):671–683.

Duvallet, C., Larson, K., Snapper, S., Iosim, S., Lee, A., Freer, K., May, K., Alm, E., and Rosen, R. (2019). Aerodigestive sampling reveals altered microbial exchange between lung, oropharyngeal, and gastric microbiomes in children with impaired swallow function. PLOS ONE, 14(5):e0216453.

Eisenberg, E. and Levanon, E. Y. (2013). Human housekeeping genes, revisited. Trends in Genetics, 29(10):569–574.

El-Solh, A. A., Pietrantoni, C., Bhat, A., Aquilina, A. T., Okada, M., Grover, V., and Gifford, N. (2003). Microbiology of severe aspiration pneumonia in institutionalized elderly. American Journal of Respiratory and Critical Care Medicine, 167(12):1650–1654.

Fernandes, A. D., Reid, J. N., Macklaim, J. M., McMurrough, T. A., Edgell, D. R., and Gloor, G. B. (2014). Unifying the analysis of high-throughput sequencing datasets: characterizing RNA-seq, 16s rRNA gene sequencing and selective growth experiments by compositional data analysis. Microbiome, 2:15.

Fordyce, J. A., Gompert, Z., Forister, M. L., and Nice, C. C. (2011). A hierarchical Bayesian approach to ecological count data: a flexible tool for ecologists. PLOS ONE, 6(11):e26785.

Friedman, J. and Alm, E. J. (2012). Inferring correlation networks from genomic survey data. PLOS Computational Biology, 8(9):e1002687. software.

Gelman, A., Carlin, J. B., Stern, H. S., Dunson, D. B., Vehtari, A., Rubin, D. B., Carlin, J. B., Stern, H. S., Dunson, D. B., Vehtari, A., and Rubin, D. B. (2013). Bayesian data analysis. Chapman and Hall/CRC.

Gelman, A. and Rubin, D. B. (1992). Inference from iterative simulation using multiple sequences. Statistical Science, 7(4):457–472.

Geman, S. and Geman, D. (1987). Stochastic relaxation, Gibbs distributions, and the Bayesian restoration of images. In Fischler, M. A. and Firschein, O., editors, Readings in Computer Vision, pages 564–584. Morgan Kaufmann, San Francisco (CA).

Geweke, J. (1991). Evaluating the accuracy of sampling-based approaches to the calculation of posterior moments. Federal Reserve Bank of Minneapolis, Research Department, Minneapolis, MN, USA.

Gloor, G. B., Macklaim, J. M., Pawlowsky-Glahn, V., and Egozcue, J. J. (2017). Microbiome datasets are compositional: and this is not optional. Frontiers in Microbiology, 8. review.

Gloor, G. B. and Reid, G. (2016). Compositional analysis: a valid approach to analyze microbiome high-throughput sequencing data. Canadian Journal of Microbiology, 62(8):692–703.

Grantham, N. S., Reich, B. J., Borer, E. T., and Gross, K. (2017). MIMIX: a Bayesian mixed-effects model for microbiome data from designed experiments. arXiv:1703.07747 [stat]. 1703.07747.

Harrison, J. G., Calder, W. J., Shastry, V., and Buerkle, C. A. (2019). Scripts from “Dirichlet-multinomial modelling outperforms alternatives for analysis of microbiome and other ecological count data”. DOI: 10.5281/zenodo.3558682. Zenodo.

Hoffman, M. D. and Gelman, A. (2014). The no-U-turn sampler: adaptively setting path lengths in Hamiltonian Monte Carlo. Journal of Machine Learning Research, 15:1593–1623.

Holas, M. A., DePippo, K. L., and Reding, M. J. (1994). Aspiration and relative risk of medical complications following stroke. Archives of Neurology, 51(10):1051–1053.

Holmes, I., Harris, K., and Quince, C. (2012). Dirichlet multinomial mixtures: generative models for microbial metagenomics. PLOS ONE, 7(2):e30126. software.

Jiang, L., Schlesinger, F., Davis, C. A., Zhang, Y., Li, R., Salit, M., Gingeras, T. R., and Oliver, B. (2011). Synthetic spike-in standards for RNA-seq experiments. Genome Research.

Johnson, M. A., Drew, W. L., and Roberts, M. (1981). *Branhamella (Neisseria) catarrhalis*–a lower respiratory tract pathogen? Journal of Clinical Microbiology, 13(6):1066–1069.

Jojic, V., Jojic, N., Meek, C., Geiger, D., Siepel, A., Haussler, D., and Heckerman, D. (2004). Efficient approximations for learning phylogenetic HMM models from data. Bioinformatics, 20(suppl_1):i161–i168.

Knight, R., Vrbanac, A., Taylor, B. C., Aksenov, A., Callewaert, C., Debelius, J., Gonzalez, A., Kosciolek, T., McCall, L.-I., McDonald, D., Melnik, A. V., Morton, J. T., Navas, J., Quinn, R. A., Sanders, J. G., Swafford, A. D., Thompson, L. R., Tripathi, A., Xu, Z. Z., Zaneveld, J. R., Zhu, Q., Caporaso, J. G., and Dorrestein, P. C. (2018). Best practices for analysing microbiomes. Nature Reviews Microbiology, 16(7):410–422.

Knights, D., Kuczynski, J., Charlson, E. S., Zaneveld, J., Mozer, M. C., Collman, R. G., Bushman, F. D., Knight, R., and Kelley, S. T. (2011). Bayesian community-wide culture-independent microbial source tracking. Nature Methods, 8(9):761. software.

Krishnamoorthy, K. (2006). Handbook of statistical distributions with applications. Chapman and Hall/CRC, Boca Raton, FL, USA.

Kruschke, J. (2015). Doing Bayesian data analysis: A tutorial with R, JAGS, and Stan. 2nd Edition. Academic Press, Elsevier, London, UK, 2 edition.

Kucukelbir, A., Ranganath, R., Gelman, A., and Blei, D. (2015). Automatic variational inference in Stan. In Cortes, C., Lawrence, N. D., Lee, D. D., Sugiyama, M., and Garnett, R., editors, Advances in Neural Information Processing Systems 28, pages 568–576. Curran Associates, Inc.

Logsdon, B. A., Hoffman, G. E., and Mezey, J. G. (2010). A variational Bayes algorithm for fast and accurate multiple locus genome-wide association analysis. BMC Bioinformatics, 11(1):58.

Love, M. I., Huber, W., and Anders, S. (2014). Moderated estimation of fold change and dispersion for RNA-seq data with DESeq2. Genome Biology, 15:550.

Lunn, D., Jackson, C., Best, N., Thomas, A., Spiegelhalter, D., Jackson, C., Best, N., Thomas, A., and Spiegelhalter, D. (2012). The BUGS book: a practical introduction to Bayesian analysis. Chapman and Hall/CRC.

Lynch, M. D. J. and Neufeld, J. D. (2015). Ecology and exploration of the rare biosphere. Nature Reviews Microbiology, 13(4):217–229.

Mandal, S., Treuren, W. V., White, R. A., Eggesbø, M., Knight, R., and Peddada, S. D. (2015). Analysis of composition of microbiomes: a novel method for studying microbial composition. Microbial Ecology in Health and Disease, 26(1):27663.

Marik, P. E. (2001). Aspiration pneumonitis and aspiration pneumonia. New England Journal of Medicine, 344(9):665–671.

Marion, Z. H., Fordyce, J. A., and Fitzpatrick, B. M. (2018). A hierarchical Bayesian model to incorporate uncertainty into methods for diversity partitioning. Ecology, 99(4):947–956.

Matthews, B. W. (1975). Comparison of the predicted and observed secondary structure of T4 phage lysozyme. Biochimica et Biophysica Acta (BBA) - Protein Structure, 405(2):442–451.

McKnight, D. T., Huerlimann, R., Bower, D. S., Schwarzkopf, L., Alford, R. A., and Zenger, K. R. (2019). Methods for normalizing microbiome data: An ecological perspective. Methods in Ecology and Evolution, 10(3):389–400.

McMurdie, P. J. and Holmes, S. (2014). Waste not, want not: why rarefying microbiome data Is inadmissible. PLOS Comput Biol, 10(4):e1003531.

Monnahan, C. C., Thorson, J. T., and Branch, T. A. (2017). Faster estimation of Bayesian models in ecology using Hamiltonian Monte Carlo. Methods in Ecology and Evolution, 8(3):339–348.

Morton, J. T., Marotz, C., Washburne, A., Silverman, J., Zaramela, L. S., Edlund, A., Zengler, K., and Knight, R. (2019). Establishing microbial composition measurement standards with reference frames. Nature Communications, 10(1):2719.

Mosimann, J. E. (1962). On the compound multinomial distribution, the multivariate \text-beta distribution, and correlations among proportions. Biometrika, 49(1/2):65–82.

Munro, S. A., Lund, S. P., Pine, P. S., Binder, H., Clevert, D.-A., Conesa, A., Dopazo, J., Fasold, M., Hochreiter, S., Hong, H., Jafari, N., Kreil, D. P., Labaj, P. P., Li, S., Liao, Y., Lin, S. M., Meehan, J., Mason, C. E., Santoyo-Lopez, J., Setterquist, R. A., Shi, L., Shi, W., Smyth, G. K., Stralis-Pavese, N., Su, Z., Tong, W., Wang, C., Wang, J., Xu, J., Ye, Z., Yang, Y., Yu, Y., and Salit, M. (2014). Assessing technical performance in differential gene expression experiments with external spike-in RNA control ratio mixtures. Nature Communications, 5:5125.

Niku, J., Brooks, W., Herliansyah, R., Hui, F. K. C., Taskinen, S., and Warton, D. I. (2019). Efficient estimation of generalized linear latent variable models. PLOS ONE, 14(5):e0216129.

Norman M. Jacobs and Harris, V. J. (1979). Acute *Haemophilus* pneumonia in childhood. American Journal of Diseases of Children, 133(6):603–605.

Nowicka, M. and Robinson, M. D. (2016). DRIMSeq: a Dirichlet-multinomial framework for multivariate count outcomes in genomics. F1000Research, 5.

Paulson, J. N., Stine, O. C., Bravo, H. C., and Pop, M. (2013). Differential abundance analysis for microbial marker-gene surveys. Nature Methods, 10(12):1200–1202.

Pawlowsky-Glahn, V. and Egozcue, J. J. (2006). Compositional data and their analysis: an introduction. Geological Society, London, Special Publications, 264(1):1–10.

Pearson, K. (1897). Mathematical contributions to the theory of evolution—on a form of spurious correlation which may arise when indices are used in the measurement of organs. Proceedings of the Royal Society of London, 60(359-367):489–498. classic.

Plummer, M. (2003). JAGS: A program for analysis of Bayesian graphical models using Gibbs sampling.

Plummer, M. (2015). rjags: bayesian graphical models using MCMC. R package version 3-15. https://CRAN.R-project.org/package=rjags.

Quinn, T. P., Erb, I., Richardson, M. F., and Crowley, T. M. (2017). Understanding sequencing data as compositions: an outlook and review. bioRxiv, page 206425.

R Core Team (2019). R: A Language and Environment for Statistical Computing. R Foundation for Statistical Computing, Vienna, Austria.

Raj, A., Stephens, M., and Pritchard, J. K. (2014). fastSTRUCTURE: variational inference of population structure in large SNP data sets. Genetics, 197(2):573–589.

Robinson, M. D., McCarthy, D. J., and Smyth, G. K. (2010). edgeR: a Bioconductor package for differential expression analysis of digital gene expression data. Bioinformatics, 26(1):139–140.

Rosa, P. S. L., Brooks, J. P., Deych, E., Boone, E. L., Edwards, D. J., Wang, Q., Sodergren, E., Weinstock, G., and Shannon, W. D. (2012). Hypothesis testing and power calculations for taxonomic-based human microbiome data. PLOS ONE, 7(12):e52078.

Sachdeva, R., Campbell, B. J., and Heidelberg, J. F. (2019). Rare microbes from diverse Earth biomes dominate community activity. bioRxiv, page 636373.

Salvatier, J., Wiecki, T. V., and Fonnesbeck, C. (2016). Probabilistic programming in Python using PyMC3. PeerJ Computer Science, 2:e55.

Scordato, E. S. C., Wilkins, M. R., Semenov, G., Rubtsov, A. S., Kane, N. C., and Safran, R. J. (2017). Genomic variation across two barn swallow hybrid zones reveals traits associated with divergence in sympatry and allopatry. Molecular Ecology, 26(20):5676–5691.

Shafiei, M., Dunn, K. A., Boon, E., MacDonald, S. M., Walsh, D. A., Gu, H., and Bielawski, J. P. (2015). BioMiCo: a supervised Bayesian model for inference of microbial community structure. Microbiome, 3:8. software.

Shenhav, L., Thompson, M., Joseph, T. A., Briscoe, L., Furman, O., Bogumil, D., Mizrahi, I., Pe’er, I., and Halperin, E. (2019). FEAST: fast expectation-maximization for microbial source tracking. Nature Methods, page 1.

Stan Development Team (2018). RStan: the R interface to Stan. R package version 2.17.3.

Tang, Z.-Z. and Chen, G. (2018). Zero-inflated generalized Dirichlet multinomial regression model for microbiome compositional data analysis. Biostatistics, 00(00):1–16.

Thomson, J., Hall, M., Ambroggio, L., Stone, B., Srivastava, R., Shah, S. S., and Berry, J. G. (2016). Aspiration and non-aspiration pneumonia in hospitalized children with neurologic impairment. Pediatrics, 137(2):e20151612.

Thorsen, J., Brejnrod, A., Mortensen, M., Rasmussen, M. A., Stokholm, J., Al-Soud, W. A., Sørensen, S., Bisgaard, H., and Waage, J. (2016). Large-scale benchmarking reveals false discoveries and count transformation sensitivity in 16s rRNA gene amplicon data analysis methods used in microbiome studies. Microbiome, 4(1):62.

Tkacz, A., Hortala, M., and Poole, P. S. (2018). Absolute quantitation of microbiota abundance in environmental samples. Microbiome, 6(1):110.

Tourlousse, D. M., Yoshiike, S., Ohashi, A., Matsukura, S., Noda, N., and Sekiguchi, Y. (2017). Synthetic spike-in standards for high-throughput 16s rRNA gene amplicon sequencing. Nucleic Acids Research, 45(4):e23–e23.

Tsilimigras, M. C. B. and Fodor, A. A. (2016). Compositional data analysis of the microbiome: fundamentals, tools, and challenges. Annals of Epidemiology, 26(5):330–335. review, very good.

van den Boogaart, K. G. and Tolosana-Delgado, R. (2013). Analyzing Compositional Data with R. Springer Publishing Company, Incorporated.

Wang, Y., Naumann, U., Eddelbuettel, D., Wilshire, J., Warton, D., Byrnes, J., Silva, R. d. S., Niku, J., Renner, I., and Wright, S. (2019). mvabund: statistical methods for analysing multivariate abundance data.

Weiss, S., Van Treuren, W., Lozupone, C., Faust, K., Friedman, J., Deng, Y., Xia, L. C., Xu, Z. Z., Ursell, L., Alm, E. J., Birmingham, A., Cram, J. A., Fuhrman, J. A., Raes, J., Sun, F., Zhou, J., and Knight, R. (2016). Correlation detection strategies in microbial data sets vary widely in sensitivity and precision. The ISME journal, 10(7):1669–1681.

Weiss, S., Xu, Z. Z., Peddada, S., Amir, A., Bittinger, K., Gonzalez, A., Lozupone, C., Zaneveld, J. R., Vázquez-Baeza, Y., Birmingham, A., Hyde, E. R., and Knight, R. (2017). Normalization and microbial differential abundance strategies depend upon data characteristics. Microbiome, 5:27.

White, A. E., Dey, K. K., Mohan, D., Stephens, M., and Price, T. D. (2019). Regional influences on community structure across the tropical-temperate divide. Nature Communications, 10(1):2646.

Yang, Y., Chen, N., and Chen, T. (2017). Inference of environmental factor-microbe and microbe-microbe associations from metagenomic data using a hierarchical Bayesian statistical model. Cell Systems, 4(1):129–137.e5.

## Supplemental References

Caporaso, J. G., Kuczynski, J., Stombaugh, J., Bittinger, K., Bushman, F. D., Costello, E. K., Fierer, N., Peña, A. G., Goodrich, J. K., Gordon, J. I., Huttley, G. A., Kelley, S. T., Knights, D., Koenig, J. E., Ley, R. E., Lozupone, C. A., McDonald, D., Muegge, B. D., Pirrung, M., Reeder, J., Sevinsky, J. R., Turnbaugh, P. J., Walters, W. A., Widmann, J., Yatsunenko, T., Zaneveld, J., and Knight, R. (2010). QIIME allows analysis of high-throughput community sequencing data. Nature Methods, 7(5):335–336.

Harrison, J., Beltran, L. P., Buerkle, C. A., Cook, D., Gardner, D., Parchman, T. L., and Forister, M. L. (2019). A suite of rare microbes interacts with a dominant, heritable, fungal endophyte to influence plant trait expression. bioRxiv, page 608729.

Jost, L. (2007). Partitioning diversity into independent alpha and beta components. Ecology, 88(10):2427–2439.

Marion, Z. H., Fordyce, J. A., and Fitzpatrick, B. M. (2015). Extending the concept of diversity partitioning to characterize phenotypic complexity. The American Naturalist, 186(3):348–361.

Wang, Y., Naumann, U., Wright, S. T., and Warton, D. I. (2012). mvabund–an R package for model-based analysis of multivariate abundance data. Methods in Ecology and Evolution, 3(3):471–474.

Westfall, P. H. and Young, S. S. (1993). Resampling-based multiple testing: examples and methods for p-value adjustment. John Wiley & Sons. Google-Books-ID: nuQXORVGI1QC.

